# Space partitioning by self-organized epithelial networks

**DOI:** 10.64898/2026.09.16.750084

**Authors:** Ronny Tonato Zambrano, Ayla Biallas, Frédérique Mittler, Alice Nicolas, Xavier Gidrol, Sophie Achard, Lionel Hervé, Guillaume Godefroy, Maxim Y. Balakirev

## Abstract

Branching epithelia build supracellular networks that must simultaneously ensure connectivity, mechanical integrity, and efficient space partitioning, yet a general framework for how such networks form beyond the endothelial lineage has been lacking. Here we show that epithelial cells from multiple branching organs spontaneously self-organize into extended reticulate networks in simplified environments, revealing a conserved network-forming capacity. Combining wide-field lensless holographic imaging, deep-learning segmentation (EpiNet), and graph-theoretic analysis, we resolve a reproducible sequence of contact initiation, clustering, percolation, and post-percolation relaxation and quantify its geometry across scales. Actomyosin contractility controls both the growth of connectivity and the relaxation dynamics that set network geometry, tuning the effective cost of forming connections and thereby selecting between tree-like and reticulate topologies. After percolation, epithelial networks behave as active tension networks that progressively refine space partitioning toward centroidal, near-optimal configurations while maintaining a characteristic mesh size through continuous edge nucleation. These findings establish epithelial network formation as a generic, physically regulated mode of tissue self-organization and provide a quantitative framework linking single-cell mechanics, network topology, and space-partitioning dynamics, with implications for branching morphogenesis, organoid models, and tissue engineering.

## INTRODUCTION

Supracellular networks are ubiquitous in tissues, reflecting evolutionary optimization of transport, energy use, and mechanical fitness ^1–14^. Their architectures fall into two recurrent classes: tree-like (loopless) and reticulate (looped). These represent alternative solutions to a universal problem: how to establish connectivity while partitioning space efficiently across scales. Tree-like architectures are typically the terminal outcome of branching morphogenesis, following principles such as Murray’s law and allometric scaling that minimize transport cost and optimize flow ^8,10,15–18^. Reticulate networks, by contrast, characterize the early morphogenesis of the trachea, vasculature, lymphatics, pancreas, and bile ducts ^19–25^, and recur in pathologies such as cancer ^26^. Once dismissed as redundant intermediates, loops are now recognized to confer robustness and tolerance to local failure, an advantage that is especially valuable at early developmental stages ^3,5,7,8,14–16,18,27–30^.

Which architecture prevails is set by morphogenetic regulation acting through geometric and mechanical constraints. Morphogen gradients, tissue boundaries, space-filling requirements, and large-scale forces impose global partitioning rules that favor hierarchical, tree-like solutions ^4,6,9,12,19,31–33^, and theory indicates that the connection costs associated with such global regulation are central to network performance and likely act as key evolutionary determinants ^3,5,7,8,13,14,16,18,27–30,34–36^. When these constraints are relaxed, reticulate topologies emerge spontaneously ^5,14,29,30,36^. Consistently, diverse cell types align into transient net-like patterns *in vitro* ^37–52^ through local traction-feedback interactions with the extracellular matrix (ECM) ^53–58^. Notably, cells from branching epithelia of the vasculature, breast, kidney, and lung can potentially assemble into supracellular nets, although only endothelial cells have been shown to form continuous, lumenized tubular networks^59–63^.

How living tissues achieve optimal space partitioning from cellular to tissue scales nonetheless remains poorly understood. In particular, it is unclear what sets the characteristic length scale of a network, which cellular or supracellular parameters encode its topology, how the effective cost of forming and maintaining connections is defined, and what drives the large-scale collective remodeling seen at late morphogenetic stages, when networks behave as mechanically coupled continua ^19,22,23,53,64,65^. Endothelial *in vitro* networks have become a cornerstone for studying angiogenesis, tissue engineering, and self-organization ^56,59–63,66–75^, but no comparable model exists for epithelial branching beyond the endothelial lineage, leaving these questions largely unexplored for epithelia.

Here we address this gap. We show that supracellular reticulate self-assembly is a common property of epithelial cells from branching organs and, using holographic imaging with neural-network segmentation and graph-theoretic analysis, follow network formation as a percolation-like process organized into distinct topological stages. After percolation, connectivity stabilizes while geometry continues to evolve through active relaxation toward increasingly uniform space partitioning at nearly constant mesh density. This process is actively regulated by actomyosin, which sets both the effective cost of connections and the collective relaxation dynamics of networks treated as mechanical continua. Together, these results establish epithelial networks as a tractable active-matter system and provide a physical framework linking single-cell mechanics, network topology, and tissue-scale morphogenesis.

## RESULTS

### Reconstitution of branching morphogenesis by epithelial cells *in vitro*

We first asked whether epithelial cells retain an intrinsic capacity for branching morphogenesis in the absence of mesenchymal instruction, using human pancreatic ductal epithelial (HPDE) cells ^76^. Within Matrigel, HPDE cells formed spheroidal aggregates, a fraction of which developed central lumens lined by a polarized epithelial layer, resembling early pancreatic organoids ^77^. Optimizing Matrigel concentration and medium composition identified conditions that triggered spontaneous symmetry breaking and branch initiation, under which aggregates developed highly branched, lobulated morphologies reminiscent of pancreatic architecture **(Figure 1A)** ^78^. These branches hollowed through a caspase-dependent mechanism, revealed by cleavage of a fluorogenic caspase substrate, consistent with apoptosis-mediated lumen formation in mammary, prostate, and pancreatic epithelia ^79–81^.

**Figure 1.**
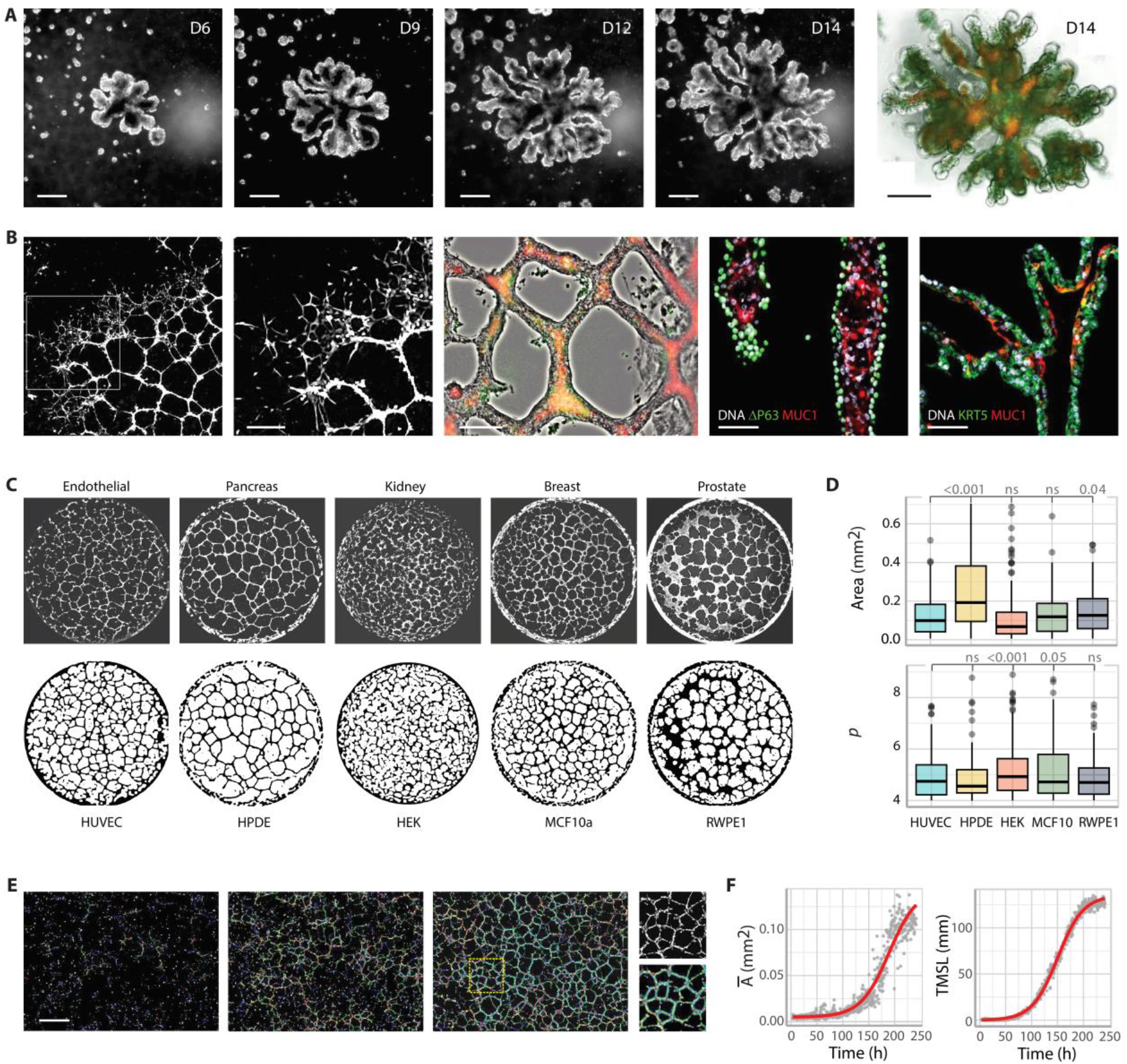
*In vitro* reconstitution of epithelial branching morphogenesis and supracellular networks. **(A)** Time course of HPDE branching morphogenesis in 3D Matrigel (days 6–14; bright-field). Right, live staining with LysoTracker Deep Red and the fluorogenic caspase-3/7 reporter CellEvent Green. **(B)** Quasi-2D epithelial network formation. Left, snapshot with zoom highlighting self-similar architecture (Video S1). Centre, mature network undergoing lumen formation revealed by LysoTracker Deep Red and CellEvent staining. Right, immunostaining of polarized epithelial tubes for ΔNp63, MUC1 and KRT5. **(C)** Representative epithelial networks formed by cell types of distinct tissue origin, visualized by Calcein-AM staining in 96-well plates; bottom, corresponding binarized images used for quantitative analysis. (**D)** Quantitative morphological characterization of epithelial networks. (**E)** Phase-reconstructed holographic image of a growing epithelial network analyzed using Angiogenesis Analyzer, with master segments (yellow) and meshes (cyan) overlaid. (**F)** Time-resolved quantification of epithelial network self-organization derived from Angiogenesis Analyzer outputs: average mesh area (Ᾱ) and the total major segment length (TMSL). Scale bars: 250 µm unless otherwise indicated; **B** (right), 50 µm; **E**, 1 mm.

Because 3D branching was infrequent and heterogeneous, limiting quantification, we reconstituted morphogenesis in quasi-two-dimensional (2D) conditions. Plated at the Matrigel–medium interface, HPDE cells spontaneously formed extended reticulate networks **(Figure 1B)**. Sparse, non-uniform seeding produced hierarchically organized networks, with thick, fused branches near the center and finer branches at the periphery, that expanded as a coordinated, wave-like front into unoccupied space **(Figure 1B, Supplementary Video S1)**, whereas uniform, denser seeding yielded isotropic networks closely resembling HUVEC endothelial capillary networks **(Figure 1B)**. Over days, these epithelial networks (ENs) matured and developed lumens by apoptosis-mediated hollowing, acquiring well-defined apicobasal polarity with basolateral ΔNp63 ^82^ and luminal MUC1 ^83,84^ **(Figure 1B, right panels)**. HPDE cells thus provide a tractable model of pancreatic epithelial self-assembly in both 3D and 2D.

To test whether this capacity is general, we screened twenty non-cancerous cell types of diverse origin **(Supplementary Table S1)**. Stable reticulate networks formed exclusively from epithelial cells derived from organs that undergo branching morphogenesis in vivo, including the kidney, mammary gland, and prostate. In contrast, cells from non-branching tissues failed to generate persistent networks **(Figure 1C, Supplementary Table S1)**. Network architecture was strikingly conserved across branching-epithelial lines, with mean mesh area Ᾱ of 0.1–0.2 mm². The dimensionless shape index *p* = *P*/√*A*, defined as the ratio of the perimeter (P) to the square root of the mesh area (A), converged to 4.0–4.5 **(Figure 1D)**. This metric quantifies polygon compactness, with higher values indicating increasingly irregular shapes and lower values corresponding to more convex, perimeter-minimizing geometries ^85–92^. Accordingly, EN meshes exhibit moderately regular polygonal shapes. Assembly consistently began with cells aligning into linear chains that branched and interconnected into a spanning 2D continuum; relative to HUVEC networks, ENs assembled more slowly (days rather than <24 h) but were far more stable, persisting for weeks. Together, these observations across multiple cell types suggest that reticulate network formation is a conserved intrinsic property of branching epithelia.

### Large-scale imaging of EN dynamics

Capturing this behavior across scales, from single cells to millimeter-scale networks over weeks, lies beyond conventional fluorescence microscopy and requires simultaneous imaging of cellular and supracellular dynamics. We therefore used an in-house multi-wavelength lensless video-holography system providing label-free, long-term imaging over a 29.4 mm² field of view ^93–96^; numerical reconstruction yielded high-contrast phase images suitable for robust segmentation and tracking, and its compact format allowed continuous imaging under physiological conditions **(Figure 1E, Supplementary Figure S1; Materials and Methods)** ^93^. Monitoring assembly for up to three weeks **(Figure 1E)**, we first quantified morphology with the Angiogenesis Analyzer ^66^, which skeletonizes binarized images into node- and edge-based descriptors: during connectivity establishment the mean mesh area Ᾱ grew, indicating coarsening, before stabilizing, while total major segment length (TMSL) rose during expansion and plateaued once the network spanned the surface **(Figure 1F)**.

This approach, however, performed poorly during early assembly and did not capture network topology, motivating a dedicated framework to convert dynamic ENs into spatiotemporal graphs and interrogate connectivity, geometry, and force-driven remodeling across scales.

### EpiNet: deep learning–based graph reconstruction of epithelial network assembly

We developed EpiNet, a deep-learning framework that segments phase-reconstructed holographic images and converts them into graphs for geometric analysis. Unlike existing pipelines, which perform unevenly across the diverse morphodynamic regimes of epithelial patterning, EpiNet maps uniformly from isolated cells to fully reticulated networks.

In the workflow **(Figure 2A)**, images are segmented and skeletonized by fast parallel thinning, then converted into undirected, unweighted graphs *G* = (*V*, *E*) whose vertices are junctions or branch points and whose edges are intercellular connections, analyzed with a NetworkX-based pipeline ^97^. Segmentation uses a U-Net convolutional network ^98^, whose encoder–decoder architecture with skip connections preserves fine features and converges stably **(Figure 2A, framed panel)**, making it well suited to the strong heterogeneity in thickness, connectivity, and topology of assembling ENs.

**Figure 2.**
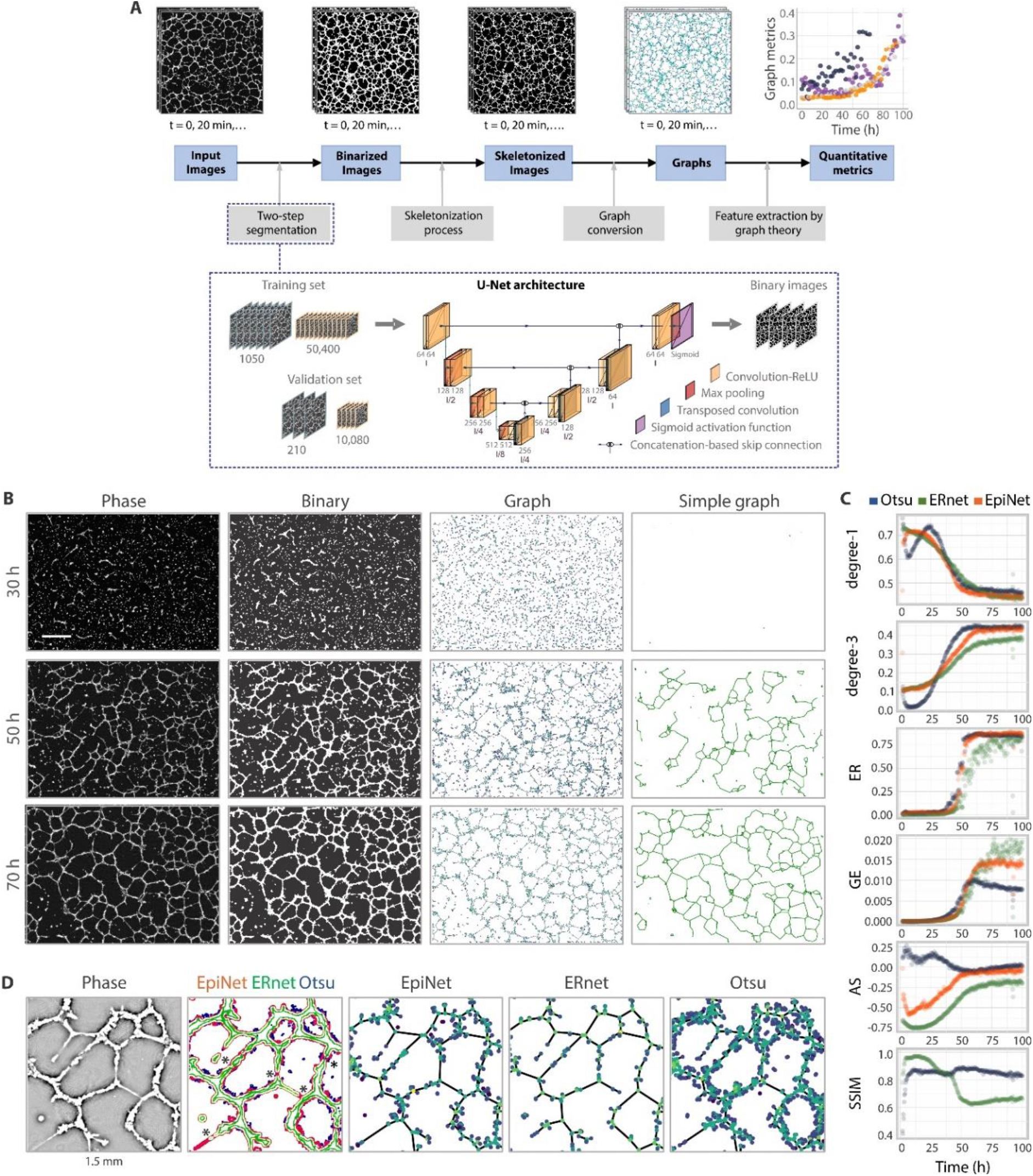
EpiNet analytical framework for epithelial network quantification. **(A)** EpiNet workflow for spatiotemporal analysis of epithelial networks, integrating holographic imaging, deep-learning–based segmentation and graph-theoretic analysis. Inset, U-Net architecture used for image segmentation. The encoder and decoder comprise blocks of two 3 × 3 convolutional layers followed by ReLU activation (orange). Encoder blocks include 2 × 2 max-pooling layers for down-sampling (red), while decoder blocks use 2 × 2 transposed convolutions for up-sampling (dark blue-green) and skip connections to the corresponding encoder features. A sigmoid activation is applied to the final output (magenta). **(B)** Representative epithelial network morphologies at three stages of growth processed by EpiNet, from phase-reconstructed holographic images to binarized representations and corresponding planar graphs. Nodes are color-coded by degree (isolated nodes, violet; degree-1, dark blue; degree-2, blue; degree-3, cyan) (Video S2). Scale bar, 1 mm. **(C)** Time-resolved graph-based quantification of epithelial network morphology obtained using three segmentation methods: Otsu (blue), ERnet (green) and EpiNet (red). Metrics include the fraction of degree-1 and degree-3 nodes, edge ratio (ER), global efficiency (GE) and assortativity (AS). Similarity between EpiNet segmentation and reference methods was quantified using the structural similarity index (SSIM; ERnet relative to Otsu in blue and to ERnet in green). **(D)** Visual comparison of segmentation outputs highlighting the limitations of ERnet in resolving thin branches (asterisks) and the over-segmentation and spurious node generation produced by Otsu thresholding - both resolved by EpiNet. Field 1.5 mm x 1.5 mm.

EpiNet was trained on holographic time-lapse datasets of HPDE networks spanning the full spectrum from sparse cells to dense reticulate structures **(Figure 2B, Supplementary Video S2)**. Because no single algorithm performed well across all stages, with some excelling at early single-cell branching and others at coarsened multicellular architectures, we curated ground-truth masks from the consensus of five independent algorithms **(Supplementary Figure S2; Materials and Methods)**. This approach avoided discontinuities in the derived metrics and provided uniform segmentation quality across stages. The model was trained with the AdamW optimizer and binary cross-entropy loss ^99,100^, reserving ∼17% of the data for validation (210 of 1260 frames; 10,080 of 60,480 cropped regions).

EpiNet converged rapidly and matched or exceeded the curated targets, reliably segmenting both single cells and complex assemblies at every stage **(Figure 2B)**. Degree-coded graph visualizations captured the progressive emergence of connectivity and loops, and pruned subgraphs proved especially effective for tracking reticulation and percolation.

From these graphs we quantified node degree (the fractions of degree-1 and degree-3 nodes), edge ratio (ER, the fraction of edges in the largest connected component), global efficiency (GE, the efficiency of information transfer), and assortativity (AS, the tendency of nodes to link to others of similar degree) **(Figure 2C; Materials and Methods)**, comparing EpiNet to ERnet, a Vision-Transformer method for endoplasmic-reticulum networks ^101^, and Otsu-based adaptive thresholding ^102^. Consistent with visual inspection **(Figure 2B, Supplementary Video S2)**, assembly progressed from a branch-initiation regime dominated by degree-1 nodes and negative assortativity to branch fusion and loop formation marked by rising degree-3 nodes, ER, and GE **(Supplementary Figure S3)**. EpiNet captured these transitions across all stages **(Figure 2C)**, matching ERnet early and outperforming Otsu at later, coarsened stages, and produced smooth trajectories free of the comparators’ noise; assortativity, a marker of hierarchical maturation, was well resolved by EpiNet but not by Otsu, whereas ERnet underestimated connectivity in thin branches **(Figure 2D)**. Structural-similarity (SSIM) analysis ^103^ confirmed that EpiNet combined the strengths of both methods **(Figure 2C, bottom panel)**.

EpiNet thus converts holographic data into spatiotemporal graphs, enabling quantitative characterization of EN morphogenesis at every developmental stage and providing a robust basis for probing the physics of tissue patterning.

### Topology and multiscale assembly of epithelial networks

EpiNet resolves EN topology across space and time, while the quasi-2D geometry constrains the networks to planar graphs, with edges meeting only at vertices ^104,105^. Throughout assembly, the numbers of vertices *V* and edges *E* were comparable, suggesting dominance of short branching events and a mean node degree ⟨*k*⟩ ≈ 2 ^106^, while pruning raised ⟨*k*⟩ to about 2.5, placing the backbone between tree-like (⟨*k*⟩ ≈ 2) and regular planar-mesh (⟨*k*⟩ ≈ 3) topologies **(Supplementary Figure S3; Materials and Methods)**. The physical meaning of nodes and edges evolved as assembly coarsened, with isolated cells and cell–cell contacts merging into multicellular junctions and supracellular cables **(Supplementary Videos S1 and S2)**; because multiple length scales coexist throughout, networks were treated as unweighted graphs, independent of absolute edge length or node size.

Graph metrics showed scale-dependent sensitivity **(Figure 3A)**: local measures such as the average clustering coefficient (AC) tracked short-to intermediate-range organization, including cluster formation and fusion, whereas global measures such as ER and GE responded to large-scale dynamics and rose sharply only once a critical level of connectivity was reached, marking the onset of percolation. Assortativity (AS) remained informative at all scales, revealing coordinated evolution of local and global connectivity.

**Figure 3.**
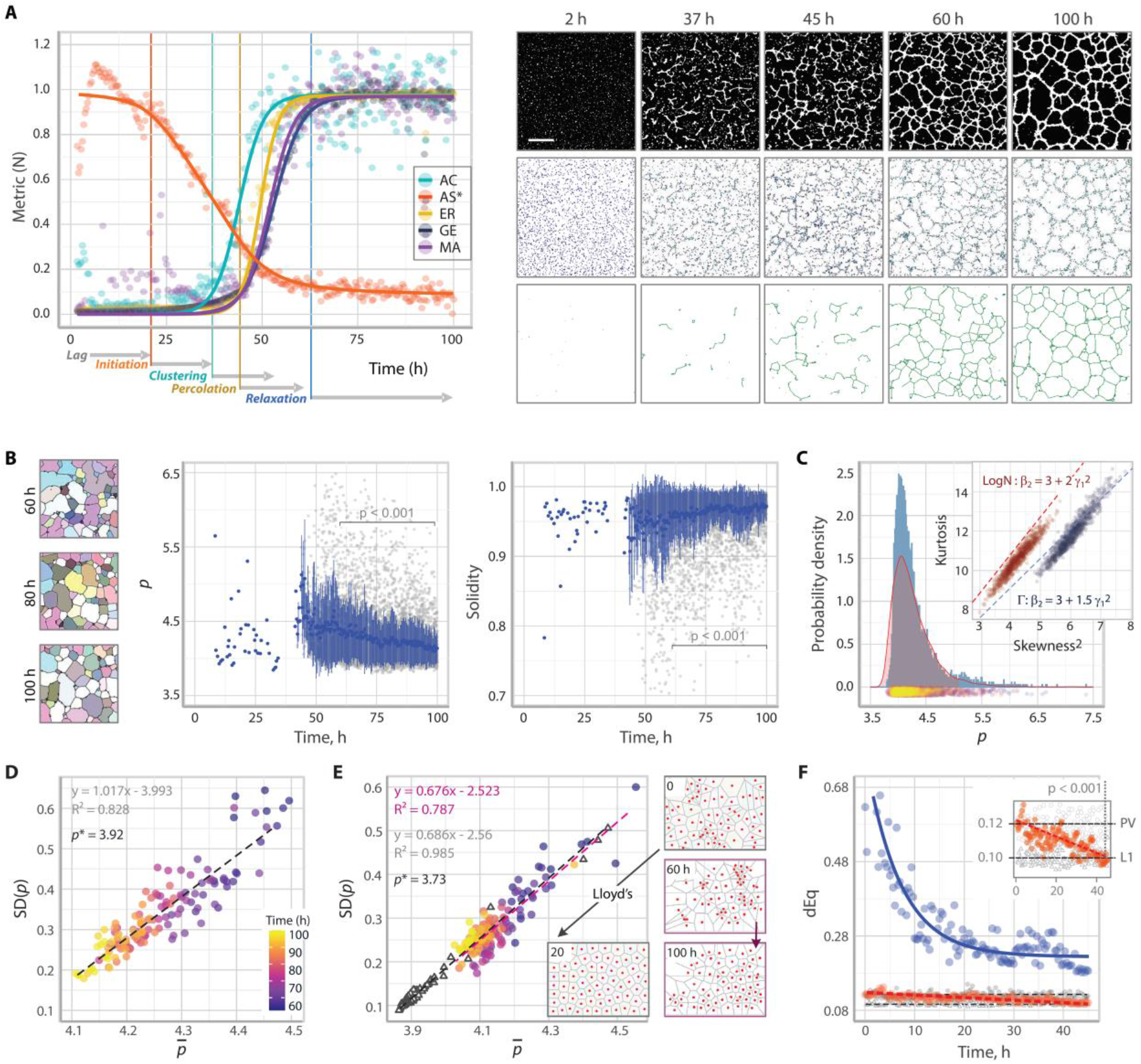
Multiscale kinetics and geometric ordering of epithelial network self-assembly. **(A)** Topological evolution of epithelial networks across scales. Right, representative snapshots at characteristic time points showing binarized images, corresponding EpiNet-derived graphs, and pruned graphs. Scale bar, 1 mm. Left, time-resolved graph metrics normalized to their plateau values: average clustering coefficient (*AC*), assortativity (plotted as the inverted assortativity value *AS*^∗^ for visualization), edge ratio (*ER*), global efficiency (*GE*), and mean mesh area (*MA*). The fraction of area covered by the network (Area), extracted from binary images, is shown for comparison. Solid lines denote fits to sigmoidal logistic functions, yielding asymptotic values (*A_min_*, *A_max_*), midpoints (*x*₀), slopes (*K*), and transition points (*x*₀ ± 2/*K*). The transition points define successive stages of epithelial network assembly (vertical lines, color-coded by the metric determining each transition). **(B)** Geometric relaxation of epithelial networks quantified by the mesh shape index *p* and solidity. Grey symbols indicate individual measurements; blue symbols show mean ± standard deviation. Left, representative network morphologies and corresponding mesh geometries reconstructed from simplified graphs at selected time points. **(C)** Probability density functions (PDFs) of *p* for epithelial networks (steel blue) and Poisson–Voronoi tessellations of comparable mesh density (100 generators; red). Mean, median and modal values are 4.24/4.14/4.04 for epithelial networks and 4.27/4.18/4.06 for Poisson–Voronoi tessellations. Lower panel, temporal distribution of epithelial network *p*₀ values (color-coded by time; see panel **D** for color scale). Inset, moment-ratio diagram of bootstrapped PDFs, including approximate boundaries for log-normal (red) and gamma (blue) distributions. **(D)** Mean *p* plotted against its standard deviation over time, fitted by linear regression. The fitted slope corresponds to the coefficient of variation (*CV*; equation shown in grey), and the *x*-intercept at *y* = 0 (*p*^∗^) is indicated in ochre. Data points are color-coded by time. **(E)** Mean–standard deviation plot for Voronoi tessellations constructed from epithelial network centroids (EN–CVT; filled circles, color-coded by time as in **D**), compared with centroidal Voronoi tessellations (CVTs) obtained by 20 Lloyd iterations from three independent Poisson–Voronoi initializations (PV-CVT, triangles). Linear regressions are shown for EN–CVT (magenta) and PV-CVT (grey). Left, representative examples of space partitioning by EN–CVT during relaxation (60 h to 100 h) and by Lloyd optimization (0 to 20 iterations; fig. S3). **(F)** Quantification of space partitioning using the dimensionless quantizer energy (*dEq*), defined as the scaled sum of the moments of inertia of partitioned domains (Materials and Methods). During relaxation, EN–CVT tessellations exhibit a decrease in *dEq* from Poisson–Voronoi values (PV, *dEq* ≈ 0.12) toward those of a one-step CVT (One Lloyds iteration L1, *dEq* ≈ 0.10) (inset). For reference, *dEq* ≈ 0.080187 for a perfect hexagonal lattice. The *dEq*^∗^ values computed directly from epithelial network meshes are shown in blue and fitted by an exponential decay.

Across metrics, EN evolution followed sigmoidal trajectories that plateaued upon formation of a spanning network, signaling approach to a quasi-stationary state **(Figure 3A, left panel)**. Normalizing the metrics resolved five reproducible stages **(Figure 3A)**: (i) a lag phase, with cells spreading as disconnected objects (⟨*k*⟩ ≈ 0); (ii) initiation, with density-dependent alignment into linear connections (⟨*k*⟩ ≈ 1 − 2); (iii) clustering, with interconnected cliques (⟨*k*⟩ ≥ 2); (iv) percolation, a rapid rise in ER and GE producing a system-spanning mesh at maximal reorganization rate **(Supplementary Figure S4A)**; and (v) relaxation, during which connectivity stabilizes while geometry continues to evolve. Because this phase governs the refinement and stabilization of the mature network architecture beyond connectivity establishment, we focused our subsequent analysis on this regime.

### Geometric relaxation and space-partitioning dynamics

During this regime, ENs underwent progressive geometric rearrangement while maintaining nearly constant global connectivity **(Figure 3A)**, reminiscent of 2D foams and vertex-based tissue models, where fixed connectivity coexists with ongoing polygonal remodeling through force balance and tension redistribution ^90,92,107–118^. Initially irregular meshes progressively regularized into more uniform, convex polygons **(Figure 3A,B)**. The mean shape index *p*, defined as *p* = *P*/√*A*, decreased from ≈ 4.5 toward ≈ 4.1 during relaxation, while increasing solidity independently confirmed enhanced convexity **(Figure 3B)**.

We compare the distribution of *p* to Poisson–Voronoi tessellations ^119–122^, a canonical model of random space partitioning. Both closely matched, in mean, median, and mode **(Figure 3C)**, though EN distributions were more skewed and heavy-tailed, reflecting residual coarsening and scale heterogeneity absent from purely random tessellations. Both EN and Voronoi *p* distributions were well fit by generalized gamma functions ^123,124^, consistent with random partitioning from isotropically distributed seeds. The mean (*p*) and standard deviation (*SD*(*p*)) of *p* distribution scaled approximately linearly, giving a nearly constant coefficient of variation (CV ≈ 1) **(Figure 3D)**, a relation reported for 2D epithelial monolayers and attributed to geometric constraints in jammed systems ^125^. Extrapolation set a lower bound *p*^∗^ ≈ 3.9, comparable to *p* values of 2D dry foams ^91^ and above the regular hexagonal limit (*p*^ℎ*ex*^ ≈ 3.72) **(Figure 3D)**.

To probe space uniformization, we compared EN relaxation with Lloyd’s centroidal Voronoi tessellation (CVT) algorithm, which minimizes quantizer energy, defined as the summed second moments of inertia of the domains, by repositioning generators at cell centroids ^126–130^ **(Materials and Methods)**. Strikingly, when EN mesh centroids were used as generators, the resulting EN-CVT *p* distributions traced the same regression toward the near-hexagonal limit (*p*^∗^ ≈ 3.73) as Lloyd iterations, with decreasing variance and comparable CV ≈ 0.7 **(Figure 3E, Supplementary Video S3)**. This was accompanied by a decrease in the dimensionless quantizer energy (*dEq*) from Poisson–Voronoi toward partially ordered CVT values **(Figure 3F)**. Computed directly on the EN meshes (modified quantizer energy, *dEq*^∗^), this energy decreased roughly threefold **(Figure 3F)**. Furthermore, the EN *dEq*^∗^ approached the *dEq* of the corresponding EN-CVTs, indicating that relaxation drives ENs toward CVT-like space partitioning, as also reported for epithelial monolayers ^131^. The concomitant reduction in shape variability and dynamical slowing are consistent with approach to the mechanically stabilized states ^85–87,89,92,125,132^, so that in supracellular ENs the shape index *p* and *dEq*^∗^ act as order parameters tracking geometric optimization.

### Actomyosin cytoskeleton controls single-cell dynamics and network architecture

Having established how ENs assemble by percolation and then relax toward optimized space partitioning, we next asked what controls both processes. Because actomyosin contractility is a central driver of branching morphogenesis and supracellular network formation *in vivo* and a key parameter in tissue models ^6,133–144^, we tested whether it controls EN behavior by shaping cell geometry, polarity, motility, and force transmission. Low, non-cytotoxic latrunculin A abolished network formation, confirming an essential role for F-actin polymerization **(Figure 4A)**. We then perturbed upstream regulators of the actomyosin cytoskeleton, including non-muscle myosin II (NM2; blebbistatin ^145^), ROCK (Y27632 ^146^), MLCK (ML7 ^147^), and the polarity regulator Cdc42 (ML141 ^148^), at non-cytotoxic doses **(Figure 4B)**. Total actin and NM2 were unchanged, whereas ROCK and MLCK inhibition lowered regulatory-light-chain phosphorylation (pMLC), a canonical readout of contractile activity ^149–152^ **(Figure 4C)**.

**Figure 4.**
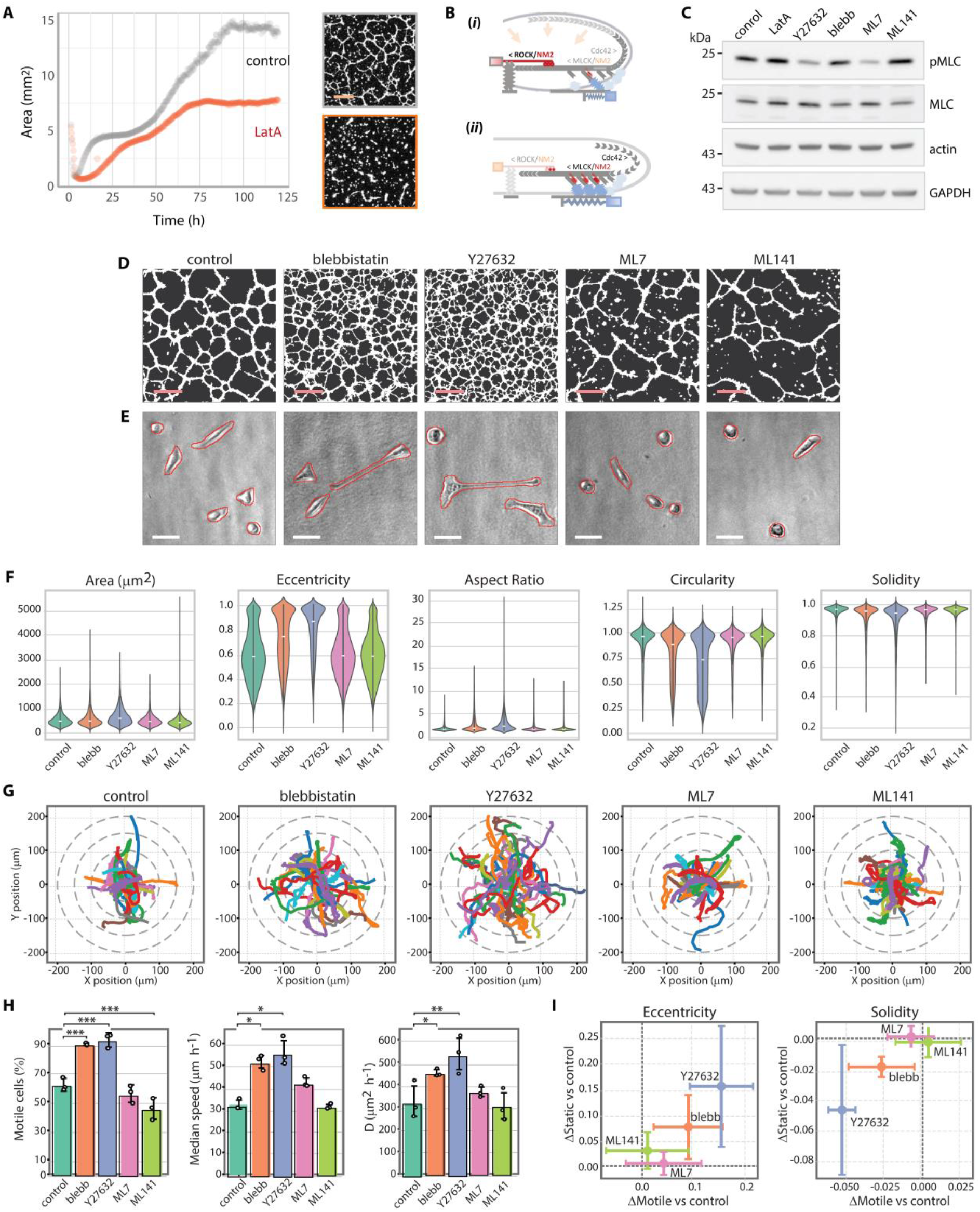
Actomyosin regulation of single-cell morphology and dynamics. **(A)** Time course of area occupation by self-assembling epithelial networks quantified from binary images under control conditions (grey) and upon Latrunculin A (LatA, 50 nM) treatment (red). Right, representative final network morphologies. Scale bar, 1 mm. **(B)** Schematic of the molecular mechanism highlighting major upstream pathways perturbed in this study. Two limiting regimes are illustrated: **(i)** high contractile tension and pronounced retrograde actin flow mediated by the ROCK–NM2 axis, with reduced cell–substrate traction upon Cdc42 or MLCK inhibition, associated with rounded cell morphologies; **(ii)** reduced contractility and retrograde flow upon ROCK or NM2 inhibition, associated with flattened, polarized cells and increased migration efficacy. **(C)** Effects of actomyosin perturbations on key cytoskeletal markers, including actin, myosin light chain (MLC), and phosphorylated MLC (pMLC), assessed by western blotting. **(D)** Binary images illustrating the effects of actomyosin inhibitors on epithelial network morphology. Scale bar, 1 mm. **(E)** Representative bright-field confocal images of single HPDE cells overlaid with Cellpose segmentation masks used for quantitative morphology analysis (Video S4). Scale bar, 50 μm. **(F)** Quantitative analysis of single-cell morphology under the different inhibitor treatments. **(G)** Representative single-cell trajectories extracted using the TrackMate algorithm and used to quantify cell motility. **(H)** Histograms summarizing the effects of actomyosin perturbations on key motility parameters: motile fraction, effective diffusion coefficient, and median displacement speed. **(I)** Wasserstein-distance analysis of cell eccentricity and solidity distributions, showing that inhibitor-induced morphological changes similarly affect motile and non-motile cell populations.

All treatments reshaped EN morphology, but distinctly **(Figure 4D)**: NM2 or ROCK inhibition produced denser, more reticulated networks, whereas MLCK and, more strongly, Cdc42 inhibition impaired long-range connectivity and network integrity. To connect these phenotypes to single-cell behavior, we engineered HPDE cells co-expressing membrane (Myr-mGL ^153^) and nuclear (mCherry-H2B ^154^) markers **(Materials and Methods)**, which faithfully reproduced parental self-organization and allowed simultaneous readout of shape and motility **(Figure 4E, Supplementary Video S4)**; TrackMate ^155^ tracking and Cellpose ^156^ segmentation then yielded trajectories and shape descriptors at high temporal resolution **(Figure 4E-I)**.

NM2 or ROCK inhibition caused pronounced elongation and polarization (higher aspect ratio and eccentricity, lower circularity and solidity), whereas MLCK or Cdc42 inhibition barely affected shape **(Figure 4F)**. Shape distributions were quasi-bimodal, with the elongated population enriched under blebbistatin or Y27632 **(Figure 4F, Supplementary Figure S5A)**. Consistent with elongation as a migratory hallmark, blebbistatin and Y27632 raised the motile fraction, mean speed, and effective diffusion coefficient, whereas ML7 and ML141 reduced the motile fraction (ML7 modestly but reproducibly) **(Figure 4G,H, Supplementary Figure S5B)**. Wasserstein-distance ^157^ comparison of motile and static cells showed that blebbistatin and Y27632 promote polarization independently of motility **(Figure 4I, Supplementary Figure S5E)**, indicating that polarization precedes and enables migration rather than following it. Although blebbistatin, Y27632, and ML7 all target NM2 activity, their effects differed markedly, reflecting distinct molecular targets and modes of action ^158–164^.

These observations align with reports that low-dose Y27632 or blebbistatin increase aspect ratio while lowering traction force on compliant substrates ^165^, reorganizing actin into aligned, long-axis stress-fiber-like bundles. Reducing tension thus does not prevent EN assembly; rather, altered polarity and cytoskeletal organization tune migration and contact formation, reshaping network topology. Enhanced polarization under reduced contractility, with preserved actin polymerization ^165,166^, promotes spreading and motility across diverse cell types ^160,161,167–169^, whereas the sparse, poorly connected networks under ML7 are consistent with impaired peripheral NM2 activation ^170^, altered adhesion dynamics ^171^, and destabilized lamellipodia ^172,173^, and ML141 suppresses migration and connectivity by disrupting Cdc42-dependent polarity and actin polymerization ^148,174–176^. Coordinated regulation of actin polymerization, polarity, and contractility is therefore a central control of EN self-assembly.

### Network self-organization under actomyosin cytoskeleton modulation

We next examined how modulation of the actomyosin cytoskeleton propagates from single-cell behavior to collective self-organization, linking changes in polarization, motility, and encounter rate to assembly kinetics, percolation, and network topology.

EpiNet reconstruction **(Supplementary Video S5)** with sigmoidal fitting confirmed that every condition followed the same five-stage sequence identified above **(Figure 3A)**, comprising lag, initiation, clustering, percolation, relaxation, but with markedly different kinetics **(Figure 5A,C)**. ROCK or NM2 inhibition accelerated assembly, shortening the lag phase to ∼6 h versus ∼20 h in controls, whereas Cdc42 or MLCK inhibition delayed initiation (∼25 h and ∼22 h). This supports a hierarchical cascade in which single-cell polarization and motility control early alignment and contact formation, setting the timescale for subsequent connectivity transitions.

**Figure 5.**
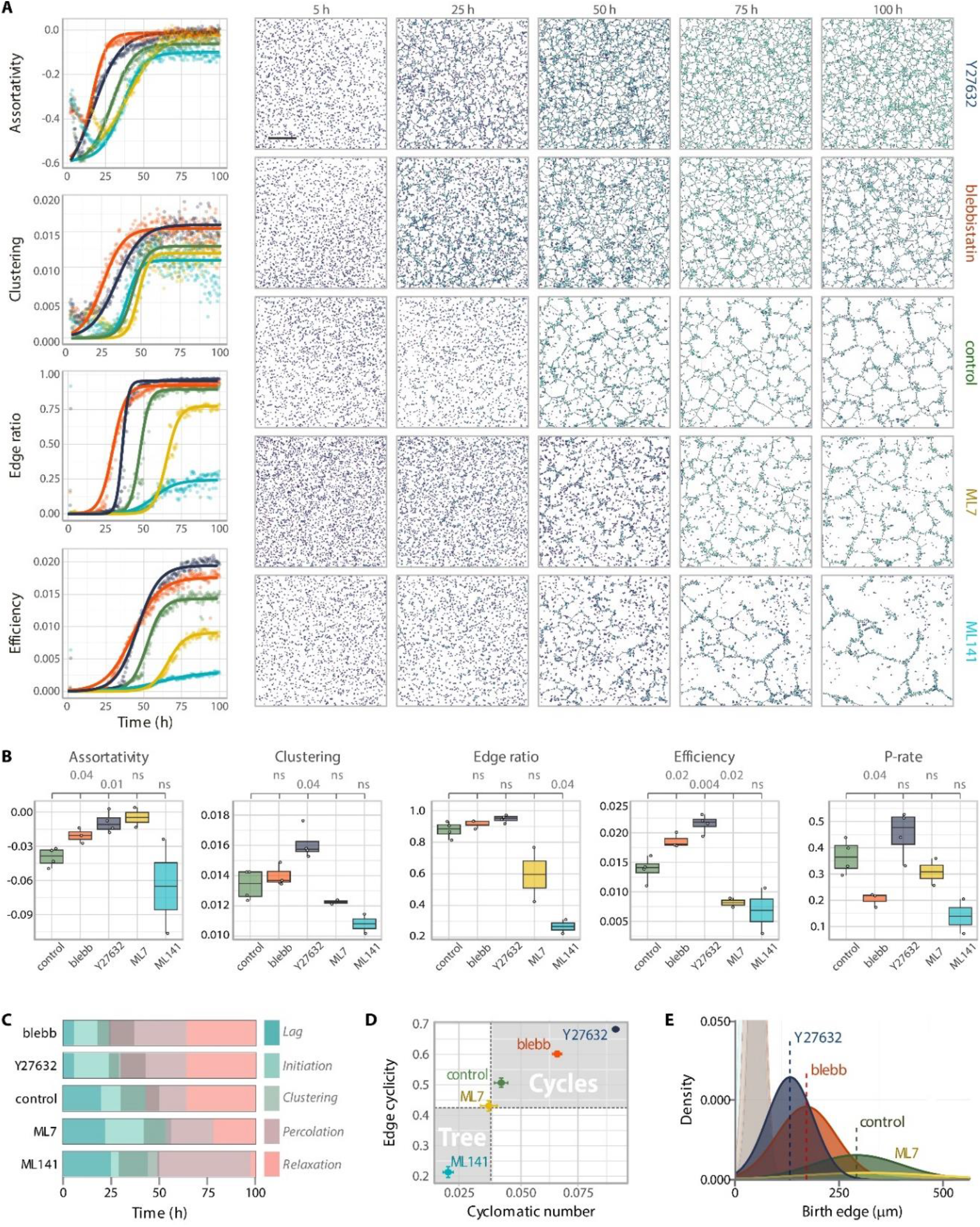
Actomyosin regulation of epithelial network self-assembly. **(A)** Effects of actomyosin perturbations on epithelial network (EN) topology and assembly kinetics. Right, spatiotemporal evolution of ENs reconstructed as EpiNet graphs (node and edge color code as in Fig. 2B) under the indicated treatment conditions. Left, time-resolved graph metrics quantifying network assembly; datapoints are color-coded by condition and fitted with sigmoidal functions (as in Fig. 3A) to extract characteristic times and transition points defining EN assembly stages (summarized in Fig. 5C). Scale bar, 1 mm. **(B)** Final (plateau) values of the indicated graph metrics measured at the end of the assembly kinetics across replicates. The percolation rate (P-rate) is defined as the slope *K* of the sigmoidal fit to the edge ratio (*ER*) metric. **(C)** Duration of the characteristic stages of EN assembly under the different actomyosin perturbations, derived from the fitted transition points shown in **(A)**. **(D)** Effect of actomyosin modulation on final EN topology, classified as tree-like or reticulate based on the cyclomatic number and edge cyclicity of the mature networks. **(E)** Cycle birth statistics quantified as the distribution of maximal edge lengths closing individual cycles. The distributions are bimodal, with a short-length peak (∼60 μm), comparable to a single-cell scale, and a second peak corresponding to space-partitioning meshes (>100 μm). The latter peak is abolished upon ML7 and ML141 treatment, indicating suppression of large-scale loop formation. In **(A)**, **(B)**, **(D)**, and **(E)**, identical color codes denote treatment conditions.

Accordingly, Y27632 and blebbistatin increased cell–cell encounter frequency, nucleating short chains and local loops with an early rise in assortativity **(Figure 5A, Supplementary Video S5)**. Assortativity rose before other motility-dependent metrics, indicating that local alignment precedes large-scale displacement during early contact formation, consistent with polarization preceding motility **(Figure 4I)**. At later stages, connectivity reflected both the edge-formation rate (set by speed and motile fraction) and the stability of new contacts (set by the balance of cell–cell adhesion and edge tension), and the assembly timescale varied inversely with single-cell motility **(Figure 5A,C)**.

Nascent assemblies seeded growth, coarsening, and ultimately percolation. Matching their faster initiation, ROCK or NM2 inhibition rapidly crossed the percolation threshold (*t_perc_* ≈ 37 h for Y27632; ≈ 31 h for blebbistatin), yielding spanning meshes of high final connectivity (ER ≈ 0.95 and 0.92) **(Figure 5A-C)**. Controls percolated later (*t_perc_* ≈ 49 h) but efficiently (ER ≈ 0.89), forming a planar graph of slightly lower reticulation (⟨*k*⟩ ≈ 2.55 for the simplified graph, versus ⟨*k*⟩ ≈ 2.76 and 2.76 for Y27632 and blebbistatin; **Supplementary Figure S3**), reflecting the larger meshes and higher degree-2 fraction of control networks. MLCK inhibition sharply delayed percolation (*t_perc_* ≈ 69 h) and plateaued just above threshold (ER ≈ 0.75), shifting the architecture from reticulate mesh to fragmented, tree-like **(Figure 5D,E)**, and Cdc42 inhibition abolished percolation entirely, leaving isolated branches. Global efficiency discriminated these regimes most sharply, from highly reticulate, efficient networks (*GE_Y_*_27632_ > *GE_blebbistatin_* > *GE_control_*) to minimally reticulated trees (*GE_ML_*_7_ ≫ *GE_ML_*_141_) **(Figure 5A-E)**.

Percolation kinetics also revealed a ROCK/NM2 difference unseen at the single-cell level: despite earlier initiation, blebbistatin percolated more slowly (*K_perc_* ≈ 0.25) than Y27632 (*K_perc_* ≈ 0.53) or control (*K_perc_* ≈ 0.35) **(Figure 5A,B)**. Because effective percolation depends on both edge formation and edge stability, this likely reflects reduced stability of nascent contacts under direct NM2 inhibition ^177^. Indeed, ROCK and NM2 act through overlapping but distinct pathways: ROCK can tune actin turnover and tension independently of NM2 ^178^, while some NM2 effects on adhesion and contact stability are ROCK-independent ^160,163,172^.

Overall, modulation of the actomyosin cytoskeleton produced a sharp topological switch from branched, tree-like to densely reticulated meshes, captured by cyclomatic number, edge cyclicity, and cycle-birth statistics **(Materials and Methods)**: cycles collapsed under ML7 and ML141 but proliferated under blebbistatin and Y27632 **(Figure 5D,E)**, where shorter characteristic birth-edge length and smaller mesh area accompanied higher encounter rates and connectivity, while reduced cortical tension stabilized extended interfaces, increasing total network length, as addressed below.

Together, these results suggest that the actomyosin cytoskeleton controls EN topology through coupled kinetic and energetic processes that regulate polarization, motility, encounter rate, and the cost of edge formation, thereby shaping the balance between tree-like and reticulate architectures.

### Actomyosin controls space-partitioning by epithelial network

Beyond topology, we examined post-percolation space partitioning. Where a spanning network formed, the mean mesh area remained nearly constant, exhibiting only weak growth and thus indicating suppressed coarsening **(Figure 6A)**. Actomyosin inhibition lowered the stationary mesh area (Ᾱ*_control_* ≫ Ᾱ*_blebbistatin_* > Ᾱ*_Y_*_27632_) **(Figure 6A)**, consistent with the shorter characteristic birth-edge length **(Figure 5D)**. Yet substantial dynamics persisted during relaxation, quantified by frame-difference (*FD*) analysis of binarized images **(Figure 6b, Supplementary Figure S4)**: displacement amplitudes were largest under ROCK and NM2 inhibition and decayed over time, converging for control and Y27632, whereas blebbistatin slowed relaxation, consistent with the structural instability noted above **(Figure 6B)**.

**Figure 6.**
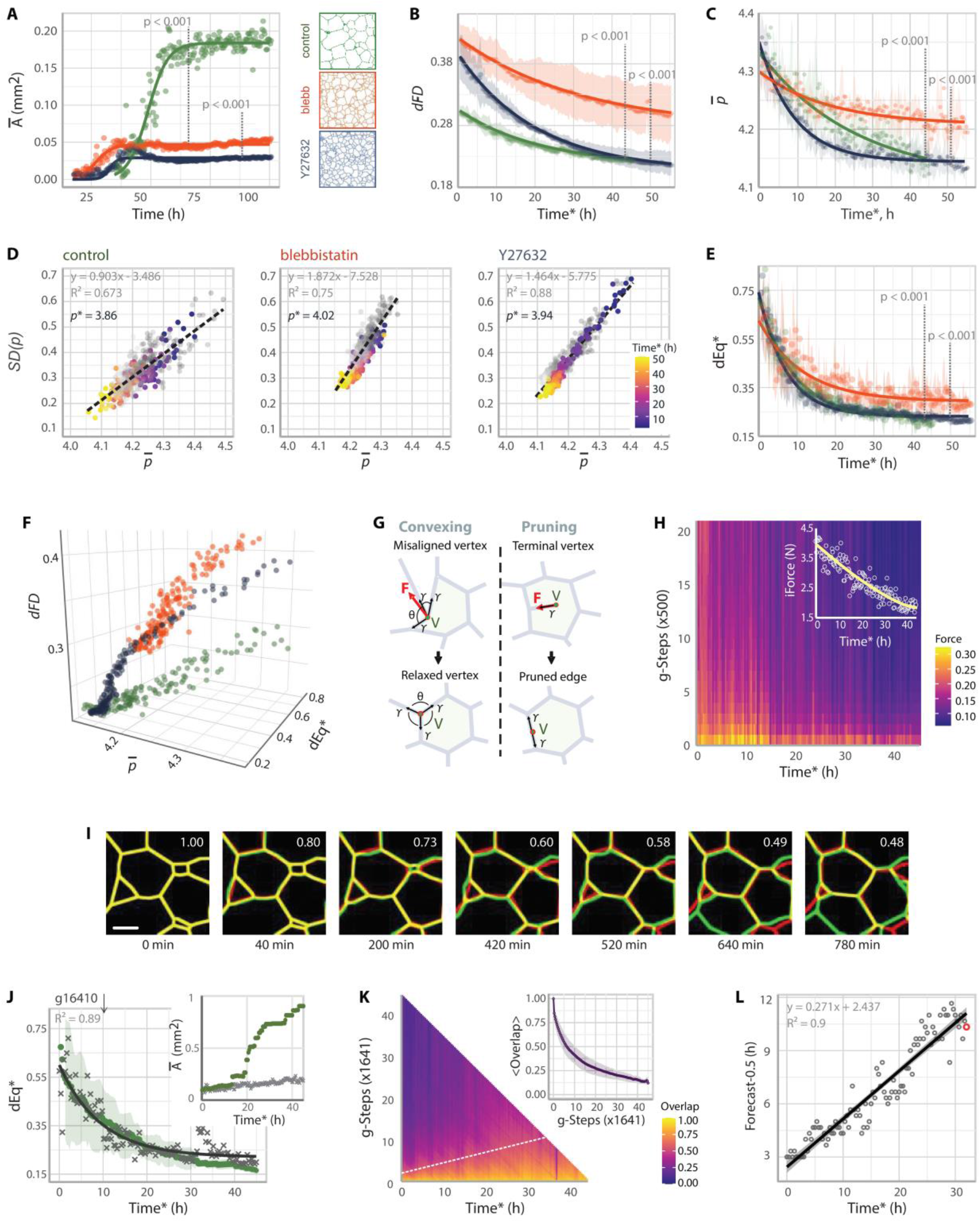
Regulation of epithelial network relaxation and space partitioning. **(A)** Effect of ROCK–NM2 inhibition on mean mesh area during EN self-assembly and relaxation. Representative final EN morphologies are shown; treatment conditions are color-coded as in Fig. 5. **(B)** Frame-difference (FD) analysis of EN dynamics during relaxation. The pixel-wise difference between consecutive binarized frames was normalized by the corresponding moving average to obtain a dimensionless metric (*dFD*) reporting the fraction of pixels undergoing change. Replicate means (filled circles) and s.d. (error bars) are color-coded as in **(A)**. Time is measured from the onset of the relaxation stage (Time*, Fig. 5C). Solid lines show exponential fits. **(C)** Geometric relaxation of epithelial networks quantified by the temporal evolution of the mean mesh shape index *p*. Symbols represent replicate means and error bars indicate standard deviation; solid lines show exponential fits. **(D)** Mean–standard deviation analysis of *p* distributions for the indicated conditions, as described in Fig. 3D. Linear regressions are shown, with the slope corresponding to the coefficient of variation (*CV*) and the extrapolated zero-variance intercept (*p*^∗^). One representative dataset is color-coded according to relaxation time (Time*); remaining datasets are shown in grey for clarity. **(E)** Quantitative analysis of space partitioning based on the modified dimensionless quantizer energy *dEq*^∗^. Symbols represent replicate means and error bars indicate s.d.; solid lines show exponential fits. **(F)** Relaxation phase space linking EN geometry and dynamics through three dimensionless parameters: *dFD* **(B)**, *p* **(C)**, and *dEq*^∗^ **(E)**. Mean values are shown. **(G)** Schematic of tension-driven vertex rearrangements underlying network relaxation. Each incident edge exerts a force of magnitude *γ* along its local tangent. The vector sum of these forces at angles θ determines the net force acting on the vertex and approaches zero at mechanical equilibrium. Such rearrangements increase mesh convexity and promote the pruning of unstable dangling edges. **(H)** Energy landscape of a control EN during relaxation probed using Surface Evolver. The heatmap shows the magnitude of residual force imbalance as a function of experimental relaxation Time* and gradient-descent steps. Inset, integrated residual force after 10^4^ gradient-descent steps, reporting the distance of the experimental network from a local mechanical minimum at each Time* point. **(I)** Comparison of experimental (green) and simulated (red) network evolution quantified by network overlap after rescaling gradient-descent steps to physical time. Corresponding overlap values are indicated on the images. The appearance of a new edge (in green, 640 min) illustrates a typical domain-splitting event observed experimentally but not captured by the model (Video S7). Scale bar, 360 μm. **(J)** Simulation of space-partitioning dynamics using STRING. Experimental EN geometries sampled at beginning of relaxation (Time* = 0, 20, and 40 min) were evolved by gradient descent and the corresponding *dEq*^∗^ values were computed. Replicate means (green circles) and s.d. are shown. Simulated and experimental (grey crosses; exponential fit shown as solid black line) relaxation kinetics were aligned by rescaling gradient-descent steps to physical time using ordinary least squares, mapping 16 410 steps to 10 h. Inset, evolution of mean mesh area Ᾱ in simulations (green) and experiments (grey). **(K)** Prediction horizon of STRING simulations quantified from network overlap measurements (as in **(I)**). The white dashed line shows the trend in overlap half-time *t*_0.5_. Inset, decrease in network overlap with simulation steps averaged across all initial conditions. **(L)** Increase in prediction horizon *t*_0.5_ as a function of network relaxation state (Time*) corresponding to the white dashed line in **(K)**. The red circle corresponds to the example shown in panel **(I)**.

Geometric analysis (as in **Figure 3**) showed that disordered networks evolved toward more regular polygonal domains, with a decreasing mean shape index (*p*), and toward more efficient space partitioning, reflected by reduced quantizer energy (*dEq*^∗^) **(Figure 6C-E)**. Nevertheless, blebbistatin sustained a higher level of disorder, characterized by slower relaxation kinetics and elevated *p* (and extrapolated *p*^∗^), *CV*, and *dEq*^∗^ **(Figure 6C-E)**, summarized in a combined geometry-dynamics plot **(Figure 6F)**. Thus, whereas ROCK inhibition raises connectivity with only modest effects on geometric ordering and stability, blebbistatin promotes a more disordered, fluid-like state.

Time evolution of EN geometry reflects the force balance governing relaxation. By analogy with dry foams and epithelial monolayers ^116–118,134,137,179^, and because inter-domain pressure differences and elastic deformations are negligible here, force balance is dominated by effective line tension along the edges. At steady state the actively injected, stored mechanical energy *E_st_* is therefore approximately interfacial: in a quasi-2D description, *E_st_* ∼ *γ L_tot_*, with *γ* the effective line tension and *L_tot_* the total network length. Since mean mesh area scales inversely with the square of edge density, Ᾱ ∼ (1/ *L_tot_*)^2^ ∼ (*γ* / *E_st_*)^2^, lowering contractility should reduce Ᾱ by decreasing *γ* **(Figure 4B)**; assuming constant *E_st_*, the observed 4–5-fold reduction in Ᾱ under actomyosin inhibitors **(Figure 6A)** implies an approximately twofold decrease in effective line tension *γ*.

Vertex-angle equilibration toward Plateau-like configurations, frequent T0 and occasional T1 rearrangements, and pruning of dangling edges together indicate tension-driven remodeling **(Figure 6G, Supplementary Video S6)**. To test whether line-tension equilibration alone could account for the ordering, we simulated relaxation with the STRING model in Surface Evolver, which evolves geometry by gradient descent of interfacial energy ^180,181^. The energy landscapes showed a transition from stressed configurations, characterized by force-imbalanced vertices and numerous dangling edges, toward local minima with reduced force imbalance **(Figure 6H)**, indicating that much of the assembly-generated disorder can be removed through local tension equilibration and vertex rearrangement.

Rescaled to physical time, STRING reproduced the key hallmarks of post-percolation relaxation **(Figure 6I-L)**, including vertex-angle equilibration, edge-length remodeling, pruning of unstable branches, and T0 transitions accompanying mesh collapse **(Figure 6I, Supplementary Video S7)**. It also captured the qualitative decrease in *dEq*^∗^ **(Figure 6J)**. However, agreement between simulations and experiments was limited by the persistence of biological remodeling, yielding a prediction horizon that depended on the network’s initial relaxation state. Overlap was rapidly lost immediately after percolation, when remodeling remained high (*t*_0.5_ ≈ 3 ℎ), but persisted much longer late in relaxation (*t*_0.5_ ≈ 11 ℎ), when EN dynamics became increasingly dominated by tension equilibration **(Figure 6K,L)**.

Beyond this prediction horizon, the comparison exposed STRING’s principal limitation. Because the model minimizes interfacial energy on a fixed topology, it predicts foam-like coarsening, whereas experimental networks continue to remodel by generating new edges long after percolation. As in foams ^115,116,179^, STRING enlarged some domains at the expense of others, increasing size heterogeneity and reducing mesh number while mean area grew **(Figure 6J, inset, Supplementary Video S8).** As a result, decreases in *dEq*^∗^ reflected increasing convexity rather than improved spatial uniformity. Consistently, simulations initialized from convex Poisson–Voronoi tessellations exhibited a rise in *dEq*, confirming that line-tension minimization alone does not optimize space partitioning **(Supplementary Figure S6)**.

Experimental networks instead held mesh density nearly constant while improving uniformity. Time-lapse imaging revealed frequent edge-nucleation events - new edges emerging orthogonally from existing ones into transient T-junctions that regressed or stabilized to split domains **(Figure 6G; Supplementary Video S6 and S8)**. Such events resemble branch initiation during vascular and epithelial morphogenesis ^1,4,20,57,182,183^ and may arise from active mechanochemical patterning or local mechanical instabilities ^133,184–188^. Regardless of origin, domain splitting provides negative feedback to tension-driven coarsening, replenishing small domains and preserving the characteristic network scale.

Post-percolation EN dynamics therefore arise from two coupled processes: a tension-driven relaxation, accurately captured by STRING over its prediction horizon, that promotes force equilibration and geometric ordering; and an active domain-splitting process, that counteracts coarsening and preserves network scale. Together, these processes homogenize space partitioning while maintaining a persistent reticulate architecture.

## DISCUSSION

Branching morphogenesis requires tissues to coordinate connected transport, mechanical integrity, and space partitioning across many scales ^1–11,14,63^, yet how single-cell behavior yields organized supracellular networks remains unclear. Combining a general epithelial network (EN) *in vitro* model with large-scale imaging and graph-geometric analysis, we show that epithelial networks assemble, select their topology, and relax toward stationary, mechanically stable, space-efficient states, providing a quantitative framework linking cell dynamics, network topology, and tissue geometry.

### An experimental platform for epithelial network morphogenesis

The model developed here exploits the self-organization of epithelial cells at the ECM–medium interface. Examining 20 cell types revealed that this approach supports robust, long-lived reticulate networks only in cells derived from branching epithelial lineages. This reductionist system provides access to the physical principles behind network assembly, while preserving tissue-scale properties that are difficult to recapitulate in conventional *in vitro* models, including ECM-embedded aggregates and organoids, where network size, geometry, and connectivity are not typically maintained ^1,32,77,78,143,189–191^.

A stringent test of such a model is lumen formation, given that lumenogenesis and branching are interrelated and follow tissue-specific temporal programs ^1,6,22,62,73,79,81,192–194^. Continuous hollow-tube networks had been demonstrated *in vitro* only for endothelia ^59–63^, although lumen formation was also documented for breast, bronchial and kidney net-like assemblies ^43,44,47,49,195–197^ and is a typical feature of branching organoids ^78,80,198–204^. Using HPDE cells ^76^ as a representative epithelial model, we show that EN formation captures key stages of epithelial morphogenesis, including budding, branching, and quasi-2D reticulation, within a single platform **(Figure 1)**, followed by maturation into polarized hollow tubes with basolateral ΔNp63 and luminal MUC1 expression ^82–84^. Lumen formed through apoptosis-mediated cavitation ^79–81^, as observed during breast, prostate, and pancreatic morphogenesis. ENs thus provide accessible intermediates of ductal morphogenesis for quantitative analysis in quasi-2D and 3D.

### Multiscale analysis of epithelial network morphogenesis

Capturing this morphogenesis is challenging, as it spans microns to centimeters over timescales from minutes to months. Fluorescence microscopy ^205^, the standard approach, is limited by field of view, labeling, photobleaching, and transparency, so dynamics are usually inferred from fixed snapshots rather than followed live ^22,33,64,192,206–212^. To image continuously, we developed an in-house multi-wavelength lensless video-holography system ^213^ giving label-free imaging over a 29.4 mm² field at minute-scale resolution ^93–96^. Analysis posed a second challenge: existing pipelines are largely tied to fluorescence, with limited graph metrics ^23,70,101,214,215^, and even those formalizing images as geometric networks ^23,26,66,101,113,216,217^ rarely resolve the connectivity measures needed for topology and space partitioning. We therefore built EpiNet, a U-Net-based framework ^98^ that segments phase-reconstructed images into planar graphs ^104,105^. Trained from single cells to supracellular cords, EpiNet outperforms existing methods and, treating ENs as unweighted planar graphs, resolves connectivity from cell contacts to supracellular cables independently of edge length or node size, as in previous studies relating network topology to function. ^2,9,23,31,101,106,216,218–220^ **(Figure 2)**. Morphogenesis can thus be followed as an evolving graph capturing connectivity, mesh organization, and geometric order.

### Network assembly and connectivity formation

Graph analysis revealed that EN evolution follows reproducible sigmoidal trajectories, with the largest changes confined to a narrow window characteristic of a percolation transition ^221,222^ observed in endothelial and other biological networks ^7,14,68,75,219,223–231^ **(Figure 3A**, **5)**. By perturbing the actomyosin cytoskeleton, we further showed that this transition is not simply density driven, but instead reflects a connectivity threshold ^226^ set jointly by cell density, polarization and motility, such that identically seeded cultures percolate only where motility permits **(Figure 4, 5)**.

Graph metrics sensitive to distinct structural scales, resolve the process into four regimes **(Figure 3A**, **5A)**: initiation, where local alignment produces sparse, negatively assortative trees; clustering, where multicellular clusters form loops; percolation, where the giant connected component grows abruptly as the mesh coarsens; and relaxation, where connectivity stabilizes near a plateau. The transition from tree-like to reticulate architectures depended on the ability of cells to form and maintain new connections. Thus, polarity, protrusive activity, and encounter dynamics appear to determine whether networks remain near a branching threshold or progress toward loop-rich organization.

Modulation of the actomyosin cytoskeleton via ROCK–NM2 axis altered both the kinetics and topology of network maturation. Increasing polarity and motility with Y27632 or blebbistatin accelerated contact nucleation, increased assortativity, shortened the lag phase, and advanced percolation, whereas inhibiting actin polymerization or protrusion formation with ML7 or ML141 arrested networks in a fragmented tree-like state **(Figure 5)**. These findings suggest that the actomyosin cytoskeleton sets the mechanical cost of forming and maintaining supracellular connections, thereby selecting between tree-like and reticulate architectures, consistent with its role in the *in vivo* plexus-to-tree transition of the developing pancreas ^22,23,192^.

### Geometric relaxation and mesh regularization

Once percolated, the topology becomes relatively invariant and morphogenesis proceeds through local remodeling. This places ENs within the broader class of planar random tessellations, which includes foams, granular packings, epithelial sheets, and Voronoi tessellations ^86,90,92,109,111,112,115,116,118,125,131,138,232^. Many such systems evolve through a limited repertoire of local rearrangements, such as T0/T1 transitions, domain fusion, and division, toward statistically stationary states that can be described by maximum-entropy ensembles subject to geometric constraints.^112,115,233,234^. In ENs, global force balance along supracellular cables constitutes an additional constraint that progressively restricts the accessible geometric configurations while preserving connectivity, thereby increasing geometric order. We quantified this ordering through the dimensionless regularity index *p* = *P*/√*A*, widely used in vertex models ^85–90,92^. EN mesh *p* distributions closely resembled those of Poisson–Voronoi tessellations ^119–122^ and followed generalized gamma statistics^123,124^, as expected for isotropic random partitions. During relaxation, the mean *p* (*p*) decreased while convexity increased, indicating progressive regularization of mesh geometry. Consistent with fixed-shape gamma statistics ^123–125^, we further observed the linear relation between the mean *p* and its standard deviation (*SD*(*p*)), supporting the view that relaxation progressively narrows the ensemble of accessible geometries **(Figure 3D**, **6D)**.

This linear *p*–*SD*(*p*) relation provides a compact statistical description of geometric regularization, paralleling observations in foams, granular packings, and epithelial sheets ^91,125,235,236^. The extrapolated lower bound *p*^∗^ represents the shape regularity limit reached as shape variability tends to zero, corresponding to the most ordered tessellation compatible with the system constraints. For centroidal Voronoi tessellations, this limit corresponds to the regular hexagonal lattice (*p*^ℎ*ex*^ ≈ 3.72; **Figure 3E**). In ENs, larger *p*^∗^ values indicate that biological constraints, including force balance, topology, and active remodeling, shift a geometric optimum away from the hexagonal limit. Thus, *p*^∗^ reflects the constraint-dependent geometric offset from the ideal regular tessellation. Notably, the observed values of *p*^∗^ ≈ 3.9 **(Figure 3D**, **6D)** were comparable to *p* reported for 2D dry foams ^91^, suggesting that relaxed ENs approach a tension-balanced geometries despite persistent biological remodeling. This interpretation is further supported by the dynamical slowing accompanying the decrease in *p* revealed by frame-difference analysis **(Figure 6B,F)**, as well as by the improved predictive performance of the STRING model late in EN relaxation **(discussion below; Figure 6L)**.

### Space partitioning and centroidal organization

Geometric regularity, however, does not imply efficient space partitioning: foams with low *p* can be significantly disordered ^91,115,116^ and, therefore, suboptimal in second-moment energy *dE_q_* **(see also Supplementary Figure S6)**. We therefore compared EN relaxation with Lloyd relaxation of Poisson–Voronoi tessellations, for which iterative centroid relocation yields centroidal Voronoi tessellations (CVTs; **Supplementary Video S3**) that minimize quantizer energy ^107,127,129,130,237–239^, and uniformize space partitioning approaching a near-hexagonal, hyperuniform lattice with *p* ≈ 3.72 and *dE_q_* ≈ 0.0808 ^107,130,240^. Although experimental ENs remained far from the hexagonal optimum, EN-CVTs reconstructed from the same mesh centroids followed the Lloyd trajectory toward *p* ≈ 3.72 and progressively lower *dE_q_* **(Figure 3E,F)**. During EN relaxation, the gap between the experimental quantizer energy *dEq*^∗^and its corresponding EN-CVT value progressively narrowed, indicating that EN morphology becomes increasingly CVT-like **(Figure 3F)**, as also reported for epithelial monolayers ^131^. Thus, relaxation does not only regularize individual meshes but also improves space partitioning by increasing the geometric consistency between mesh boundaries and centroid positions. Nevertheless, ENs are less efficient than the corresponding CVTs, reflecting the constraints inherent to a living tissue.

### Force balance, coarsening, and the limits of line-tension minimization

Line-tension equilibration promotes the geometric regularization of ENs. EN relaxation proceeds through vertex-angle equilibration, T0/T1 rearrangements, and pruning of unstable edges, consistent with active tension redistribution along supracellular cables **(Figure 6G, Supplementary Video S6)** ^139–141,188,231^. Inhibiting actomyosin contractility with Y27632 or blebbistatin weakens the mechanical constraints imposed by global force balance, thereby impairing mesh regularization process, as reflected by higher coefficients of *p* variation, larger lower *p*^∗^ bounds, and displacement amplitudes (*dFD*) **(Figure 6B-F)**, paralleling observations from vertex models ^85,86,90,118,241–243^. Blebbistatin had a stronger effect than Y27632, consistent with their distinct mechanisms ^145,146,192,244–250^. Both reduced mesh size while increasing mesh density and total network length **(Figure 6A)**, as observed in endothelial networks ^67,72,251–253^ potentially reflecting reduced supracellular tension and stabilization of larger interfacial areas ^108,134,135,137,152,188,231,254–258^.

Yet pure line-tension minimization proves insufficient for geometric ordering. Relaxing experimental meshes with the STRING framework in Surface Evolver ^180,181^, as in vertex-model studies ^86^, reproduced the observed *dE*^∗^ kinetics but mainly through mesh regularization and edge pruning rather than uniformization **(Figure 6I,J)**. When initialized from convex Poisson–Voronoi tessellations, the same framework reduced partition uniformity rather than increasing it **(Supplementary Figure S6, Video S8)**. Like in foams ^91,115,116,179,259^, these STRING effects likely reflect coarsening and T0/T1^234^–driven polydispersity.

Instead, in ENs coarsening stays limited and mesh density nearly constant, implying an active stabilizing process. CVTs ensure constant mesh density through a fixed number of generators, while vertex models do so via a preferred-area term linked to cell elasticity ^85–87,90,92,117,118,138,260,261^; but in ENs, where meshes are ECM domains bounded by supracellular cords, its origin remains unknown.

### Domain splitting as a counterbalance to coarsening

Domain splitting by edge nucleation provides a plausible mechanism for limiting coarsening and stabilizing network scale. Time-lapse imaging revealed new edges sprout orthogonally from existing cables into transient T-junctions that regress or stabilize, subdividing larger domains **(Figure 6G, Supplementary Video S6)**. Such T-junction formation is a branching hallmark, driven by morphogen-controlled collective migration (e.g. VEGF, FGF) ^1,4,6,20,57,182,183^ and actomyosin-dependent mechanoregulation ^184,185^: cord tension suppresses perpendicular protrusions ^186^, and intercellular tension negatively regulates angiogenic sprouting ^251,253^. Accordingly, the elevated sprouting under actomyosin inhibition could itself densify networks despite reducing geometric regularization, as we observed with blebbistatin and Y27632 treatments **(Figure 5,6)**. Contractility may also drive branching mechanically, as tension focusing could break symmetry orthogonally to cables through mechanisms such as T1-like transitions, buckling, or ECM cracking, producing T-junction-rich patterns reminiscent of leaf venation or drying-mud fractures ^115,262,263^.

By subdividing domains, edge nucleation opposes tension-driven coarsening, as confirmed by theoretical works where sprouting sustains a quasi-stationary state ^188,231^; theory likewise shows fusion–division yields more homogeneous, statistically steadier networks than T0/T1 rearrangements alone ^234^. Uniform partitioning would then require size-dependent negative feedback, whereby the probability of edge nucleation increases with domain size to balance coarsening. Characterizing this feedback is a key next step. Actomyosin activity, which regulates mesh size, coarsening, and branching frequency and is repeatedly implicated in sprouting in vivo, may therefore regulate the balance between coarsening and splitting, coupling assembly to subsequent geometric refinement.

### Limitations of the study

Several limitations temper these conclusions. The system is deliberately reductionist, being quasi-two-dimensional and mesenchyme-free, and omits key in vivo features such as 3D geometry, stromal signaling, interstitial flow, and heterotypic interactions. Whether the connectivity-then-geometry sequence generalizes to native tissues remains an open question. Mechanistic inference relies on pharmacological perturbations of actomyosin that may have off-target effects, the proposed size-dependent feedback remains inferred rather than directly demonstrated, and the graph-based analysis provides only a coarse-grained representation of force balance.

### Toward a physical framework of epithelial network morphogenesis

Taken together, our results support a two-phase view of epithelial network morphogenesis: connectivity formation followed by constrained geometric relaxation. Cell polarization and motility first drive a connectivity-based percolation transition assembling a spanning network; connectivity then holds fixed while supracellular cables, balancing tension at vertices, relax the geometry through angle equilibration, junction elimination, and rising polygonal regularity, as in 2D foams and vertex models. Unlike passive foams, however, ENs preserve a finite mesh size because active domain splitting continually counteracts coarsening. The resulting architecture is not a static equilibrium but a dynamically maintained nonequilibrium steady state, in which connectivity, line-tension relaxation, coarsening, and domain splitting together produce homogeneous, yet suboptimal, space partitioning. EN morphogenesis thus links percolation, constrained tessellation, and active tissue mechanics, suggesting how developmental programs generate robust supracellular architectures through coupled topological, geometric, and mechanical regulation. Extending these principles to 3D tissues, and identifying the molecular mechanisms behind, is a clear path forward.

## MATERIALS AND METHODS

### Cell culture (cell passage in monolayer culture)

Unless otherwise stated, all cell lines were maintained at 37°C in a humidified 5% CO2 atmosphere (hereafter, cell incubator). The human pancreatic duct epithelial cell line HPDE/H6c7 (HPV18-E6E7-immortalized; #ECA001-FP) was obtained from Kerafast and cultured in Keratinocyte SFM medium (K-SFM, Gibco^TM^ #17005042) containing 0.05 mg ml⁻¹ bovine pituitary extract (BPE), 5 ng ml⁻¹ epidermal growth factor (EGF), and 1% penicillin-streptomycin (PS) (complete K-SFM). Cells at passages 5-7 were used for all experiments.

GFP-labeled human umbilical vein endothelial cells (GFP-HUVECs, Angio-Proteomie #CAP0001GFP) were obtained from Angio-Proteomie and cultured in complete EndoGM medium (Angio-Proteomie #CAP02). Cells at passages 2-4 were used for all experiments.

RWPE-1 cells (HPV18-immortalized human adult prostate epithelial cells; ATCC #CRL-11609) were purchased from ATCC and cultured in the same medium as HPDE cells. Cells at passages 5-7 were used for all experiments.

SuperTopFlash (STF) HEK293 cells (immortalized human embryonic kidney cells; #EJH014) were purchased from Kerafast and cultured in DMEM/Ham’s F-12 (Gibco^TM^ #11320033) supplemented with 20% bovine calf serum (ATCC #30-2030) and 200 µg ml⁻¹ G418. Cells at passages 5-7 were used for all experiments.

MCF10A cells (immortalized human mammary epithelial cells; ATCC #CRL-1573) were purchased from ATCC and cultured in DMEM/Ham’s F-12 (Gibco^TM^ #11320033) supplemented with 20 ng ml⁻¹ EGF, 10 µg ml⁻¹ insulin, 0.5 µg ml⁻¹ hydrocortisone, 100 ng ml⁻¹ cholera toxin, 5% horse serum (Gibco^TM^ #16050130), and 1% PS. Cells at passages <10 were used for all experiments.

### Three-dimensional (3D) Cell Assay

This assay was designed to evaluate epithelial self-organization and branching-like morphogenesis in a matrix-rich environment.

An overlay protocol was used to culture HPDE cells in 3D: single cells were seeded on a gelled layer of growth-factor-reduced Matrigel (Corning #CLS354230), approximately 1-2 mm thick. Multi-well plates were pre-chilled at 4°C and placed on a cold Corning^TM^ XT Starter Ice-free Cooler during Matrigel dispensing. Matrigel was diluted to concentrations of 3 mg ml⁻¹ or higher with ice-cold medium, and the required volume was dispensed using a positive-displacement pipette with pre-cooled tips (care was taken to avoid bubble formation). Plates were kept at 4°C for 15 min to allow homogeneous settling of Matrigel, then transferred to the cell incubator for 1 h at 37°C for gelation.

A single-cell suspension of HPDE cells was prepared in 3D assay medium (3D-AM) and seeded onto the Matrigel layer at 2-10 x 10^3^ cells cm⁻^2^. The 3D-AM consisted of 50% complete K-SFM and 50% DMEM supplemented with GlutaMAX^TM^, pyruvate, and 10% fetal bovine serum (FBS, CliniSciences #FBS-12A), plus 50 µg/ml Matrigel (added immediately before use).

Cells were cultured for several days. Aggregate formation was monitored by bright-field imaging every 1-2 days on an Axioimager Z1 Apotome fluorescence microscope (Zeiss). Objectives of 10× and 20× were used to image individual cells and local aggregates; a 1.25× objective was used for larger structures. For live fluorescence imaging, 3D-AM was supplemented with 0.5 µM LysoTracker^TM^ Deep Red dye (LTDR, Thermo Fisher Scientific #L12492) and 0.5x CellEvent^TM^ Caspase-3/7 Green Reagent (CE, Thermo Fisher Scientific #R37111) to detect apoptosis ^264^.

### Quasi-two-dimensional (quasi-2D) Cell Assay

This assay was used for quantitative analysis of supracellular network formation under reduced geometric complexity.

A modified 3D assay was used for quasi-2D culture: cells were seeded on top of a gelled Matrigel layer without a 5% Matrigel overlay. In 96-well black PhenoPlates (Perkin Elmer #6055300), 50 µl of 5 mg ml⁻¹ Matrigel was used to form the gel bed. Then, 40 µl/well of single-cell suspension in growth medium (the same medium used for routine passaging; 10^4^ to 10^5^ cells ml⁻¹) was seeded on top of the Matrigel and incubated for 3 h to allow attachment. Wells were then completed with 210 µl of quasi-2D assay medium (q2D-AM), composed of 50% growth medium and 50% DMEM supplemented with GlutaMAX^TM^, pyruvate, and 10% FBS, but without Matrigel (replaced by 0.4% methyl-cellulose, MeC).

Live imaging of multicellular structures was performed as described for the 3D assay. For end-point viability analysis, 4 µM Calcein acetoxymethyl ester (Calcein AM) and 10 µM verapamil were added to wells, followed by 1 h incubation in the cell incubator. Imaging was then performed with a GFP filter set.

For immunofluorescent analysis, the multicellular structures (ENs) were fixed with 4% formaldehyde for 1 h at room temperature and then processed to embed either in paraffin blocks or in OCT medium. The samples were cut using a microtome (Leica) and the slices were stained with DAPI (0.8 µg ml⁻¹), anti-MUC1 (1:100, Biolegend #BLE355602), anti-ΔNp63 (1:100, Abcam #ab203826) or anti-KRT5 (1:100, #ab52635) primary antibody, followed by Cy5-conjugated anti-mouse secondary antibody (1:1000, Jackson ImmunoResearch #115-175-146, for anti-MUC1) or FITC-conjugated anti-rabbit secondary antibody (1:1000, Jackson ImmunoResearch #111-095-144, for anti-ΔNp63 and anti-KRT5).

### Western blot analysis

For Western blot analysis, HPDE cells were cultured on circular Matrigel pads measuring 2 cm in diameter and approximately 0.4 mm in thickness. To prepare each pad, a 140 µl drop of ice-cold Matrigel (5 mg ml⁻¹) was dispensed in the center of a well of a 6-well plate (non-treated polystyrene, Falcon #351146, pre-chilled to 4 °C) and spread evenly with the tip of a P200 pipette to form a 2-cm circular area, taking care to avoid air bubbles and prevent overspreading. A paper template with printed 2-cm circles was placed under the plate to guide the spreading. The plate was then placed at 4 °C for 15 min to allow the Matrigel to settle uniformly, and subsequently transferred to a cell incubator at 37 °C for 1 h to allow Matrigel gelation. Then, 150 µl of 1.5 × 10^6^ cells ml⁻¹ single cell suspension in complete K-SFM medium was carefully added in the center of the Matrigel pad to form a settle drop entirely covering the pad surface and the plate was placed in cell incubator for 3 hours to allow cell attachment. After that, the wells were complemented with 3 ml q2D-AM. After 24 hours in cell incubator, cells were treated with 50 nM latrunculin A, or 10 µM Y-27632, or 1 µM blebbistatin, or 5 µM ML-7, or 2 µM ML141 for another 24 hours. Then the cells were recovered from Matrigel by enzymatic digestion. Culture medium was aspirated and replaced with 1 ml Advanced DMEM F12 containing 5 µl Collagenase Type 2 (10^4^ U ml⁻¹, Gibco™ #17101-015) and 20 µl Dispase (50 U ml⁻¹, Corning #CLS354235). Plates were incubated for 1 h 30 min in the cell incubator. From 1 h of incubation onward, the Matrigel suspension was gently aspirated and dispensed using a P1000 pipette whose tip had been pre-coated with 0.1% BSA, and this step was repeated one to two times to facilitate matrix dissociation while minimizing mechanical stress. The resulting suspension was transferred to a 5-ml microcentrifuge tube. Wells were then rinsed with 1 ml Advanced DMEM/F12 containing 0.1% BSA, and the rinse was pooled with the initial suspension in the same tube. Samples were centrifuged for 4 min at 1300 rpm, the supernatant was carefully discarded, and the cell pellet was washed once with PBS.

Cellular proteins were extracted using RIPA lysis buffer (Sigma #R0278) supplemented with cOmplete™ protease inhibitor cocktail (Roche #11836153001), 10 mM ortho-phenanthroline, 30 mM N-ethylmaleimide, 5 mM sodium ortho-vanadate, and 5 mM sodium fluoride. After quantification with a BCA protein assay kit (Pierce #23225), an equal amount of protein was run on NuPAGE Novex Bis-Tris Gel (Thermo Fisher Scientific #NP0322BOX) in MES buffer and then transferred onto the nitrocellulose membrane. The membranes were probed using anti-ppMLC (1:2000, Cell Signaling #3674T), anti-MLC2 (1:2000, Cell Signaling #3672S) anti-actin (1:2000, Cell Signaling #4970S), and anti-GAPDH (1:5000, Santa Cruz Biotechnology #sc-25778) antibodies at 4°C overnight and then with secondary anti-rabbit horseradish peroxidase-conjugated antibody (1:10000, Jackson ImmunoResearch #111-035-144) for 1 h at room temperature.

### HPDE bicolor reporter cell line (HPDE-bic)

HPDE bicolor reporter cells (HPDE-bic) were generated by HDR-mediated knock-in of a reporter vector at the AAVS1 locus using the CRISPR/Cas9 Safe Harbor Knock-in System. The reporter vector encoded a bicistronic transcript under the control of the CMV promoter and contained an mCherry-H2B reporter gene (amplified from Addgene plasmid #20972 ^154^) and an IRES-dependent Myr-mGL construct encoding an N-terminal myristoylation signal peptide fused to the mGreenLantern fluorescent protein (coding sequence amplified from the mGreenLantern plasmid ^153^). The reporter sequence was flanked by AAVS1 homology arms to enable HDR-mediated integration at the AAVS1 locus. The reporter vector and the pCas-Guide-AAVS1 plasmid, which encodes CRISPR/Cas9 and an AAVS1-specific sgRNA, were co-transfected into HPDE cells using FuGENE® HD Transfection Reagent (Promega, #E2311). Cells expressing the Myr-mGL green membrane marker and the mCherry-H2B red nuclear marker were selected, and stable clones were obtained by repeated sorting using a BD FACSMelody™ Cell Sorter.

### Time-lapse imaging of single cells

This assay was used to link single-cell shape and motility dynamics to network-level morphogenesis under control and inhibitor conditions.

Single-cell dynamics was analyzed in 96-well black PhenoPlate (Perkin Elmer). HPDE-bic cells were seeded using 40 µl of 8 × 10^3^ cells ml⁻¹ cell suspension following the protocol described above (quasi-2D Cell Assay). After cells were adhered, wells were completed to a total volume of 300 µl of q2D-AM, supplemented where indicated with final concentration Y-27632 (10 µM), blebbistatin (1 µM), ML-7 (5 µM), or ML141 (2 µM). Cells were then grown for 24 h in cell incubator prior to imaging.

Live imaging was performed on a spinning-disk confocal microscope equipped with an environmental chamber maintained at 37 °C with controlled humidity and CO₂ levels (EclipseTi-E Nikon inverted microscope equipped with a CSUX1-A1 Yokogawa confocal head, an Evolve EMCCD camera from Roper Scientific, Princeton Instruments). Images were acquired using a 10× objective (field of view: 683 × 683 µm; pixel size = 1.32 µm). Time-lapse sequences were collected over 16 h at 7-min intervals. Brightfield and mCherry fluorescence channels were recorded to visualize cell morphology and mCherry-H2B-labeled nuclei, respectively.

### Cell tracking and migration analysis

Nuclear tracking was performed on mCherry–H2B fluorescence images. Image sequences were exported as TIFF stacks and processed in ImageJ using the TrackMate plugin^155^ for automated cell tracking. Extracted trajectories were then filtered with a custom classification pipeline in Python to distinguish motile from static cells and to exclude trajectories arising from noise or tracking artefacts. Classification used per-frame displacement (Euclidean distance between consecutive positions), total displacement, spreading radius around the trajectory center of mass, and trajectory duration, and all parameters were validated by visual inspection.

Migration characteristics were quantified for cells classified as motile. Instantaneous velocity was calculated as the Euclidean distance between consecutive nuclear positions divided by the acquisition interval and expressed in µm h⁻¹. Mean squared displacement (MSD) was obtained by averaging squared displacements over increasing time lags, and diffusion coefficients (D) were derived by linear fits to MSD(τ) = 4Dτ (**Supplementary Figure S5**).

### Cell morphology analysis

Cell segmentation was performed on brightfield images using two pretrained Cellpose models: transformer-based Cellpose-SAM^265^, which accurately detected elongated cell morphologies, and Cellpose2-LC4^266^, which recovered additional cells missed by SAM. Masks generated by both models were merged and spatially associated with their corresponding fluorescently tracked nuclei, enabling integration of morphological data with migration classification (motile versus static). Cell morphology parameters were subsequently quantified from the resulting segmentation masks. All analyses were implemented in custom Python scripts.

### EpiNet framework

#### Lensless imaging of epithelial network self-assembly

Lensless imaging was performed with a Cytonote 6W microscope ^93–95,267^ (Iprasense, Montpellier, France) by parallel monitoring of cells in six wells of a 6-well plate (Falcon #351146) directly in the cell incubator. The cells were seeded on 2-cm circular Matrigel pads essentially as described in section “Western blot analysis”, except that the concentration of HPDE cells in single cell suspension varied from 10^4^ to 5 × 10^5^ cells ml⁻¹. The principal results described in the article (**Figure 2** to **Figure 6**) were obtained with cell seeding concentration 3 × 10^5^ cells ml⁻¹. Immediately after the wells were complemented with 3 ml q2D-AM, the plate was transferred on the microscope inside the cell incubator and the acquisitions started.

#### Image acquisition and reconstruction

We used an Iprasense system equipped with six parallel RGB LED sources spatially filtered through 50 µm diameter pinholes and positioned 50 mm from the samples. Diffraction patterns were recorded by a CMOS detector (3,840 × 2,748 pixels, pixel pitch 1.67 µm; field of view 6.4 × 4.6 mm²) placed at a sample-to-sensor distance of 2 mm, with an integration time of 300 ms. Three diffraction patterns corresponding to the RGB channels were acquired sequentially within approximately 1 s. Holographic images were acquired every 20 min over 7–27 days, yielding 500–2000 image triplets per sample.

In this lensless configuration, no optical imaging elements were used, and only intensity measurements were recorded at the sensor plane. Optical path difference (OPD) images in the sample plane were subsequently reconstructed using the previously described algorithm^268^.

The reconstruction pipeline comprised three main steps **(Supplementary Figure S1)**. First, an initial reconstruction was obtained by solving an inverse problem initialized with a zero field. Second, a convolutional neural network (CNN) was used to refine the reconstruction and correct phase unwrapping artefacts. Third, a second inverse problem was solved, initialized with the CNN output, yielding the final reconstruction. The code was implemented in MATLAB, with an execution time of ∼20 s per frame on a GPU. Further details of the algorithm can be found elsewhere^269,270^.

A critical step consisted in determining the appropriate sample–detector distance “z” parameter. This distance (typically 2 mm) was optimized once per culture condition using a representative frame by visual inspection of reconstruction quality, with z scanned between 1.5 and 3 mm in 0.05 mm increments.

**Supplementary Figure S1.**
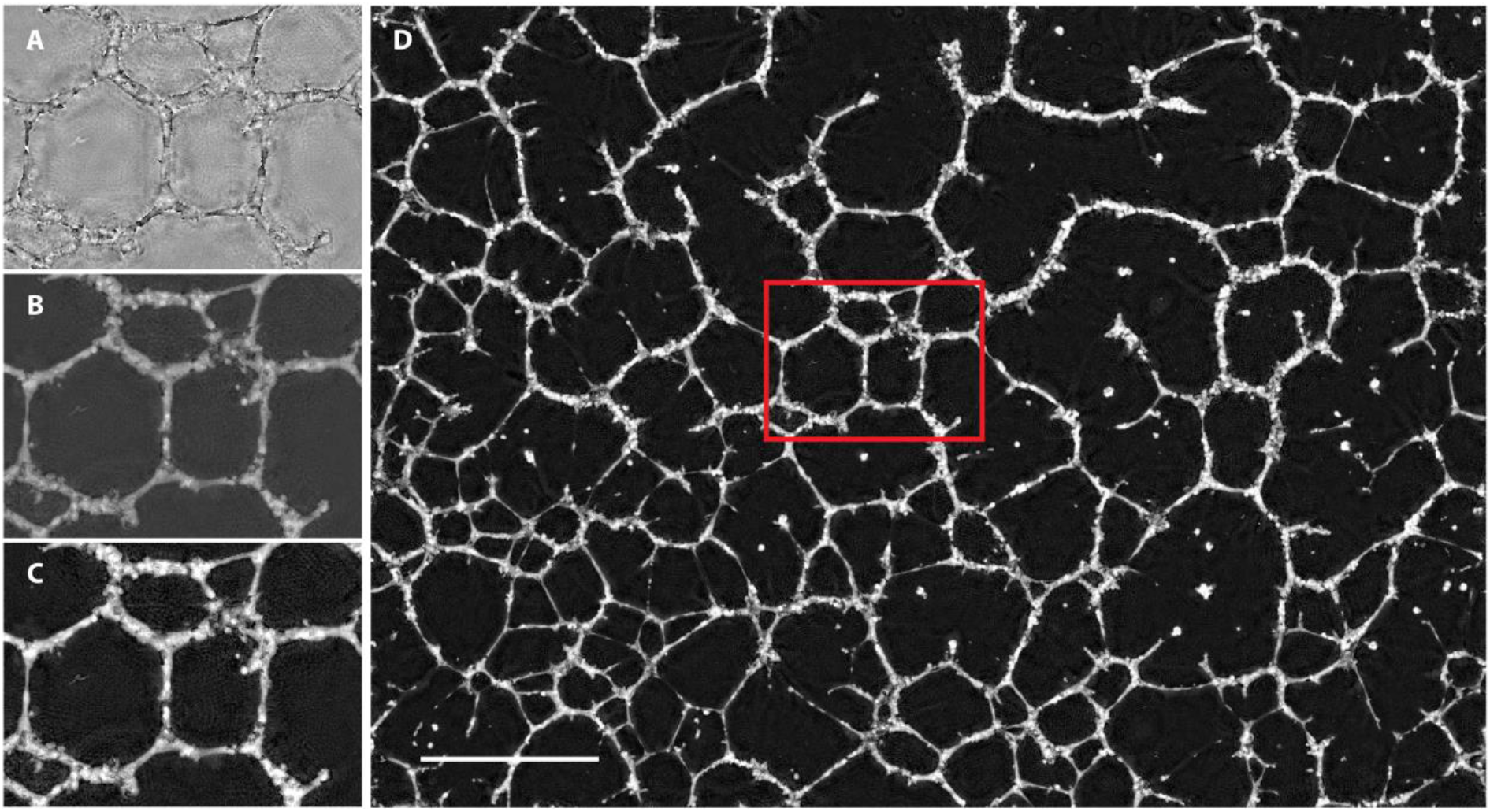
Holographic reconstruction pipeline. (A) Optical path difference (OPD) map obtained after solving the first inverse problem. (B) OPD map after inference using the phase-unwrapping convolutional neural network (CNN). (C) Final OPD map obtained after solving the second inverse problem. (D) Full-field reconstruction of the final OPD map. Scale bar, 1 mm.

#### EpiNet segmentation: architecture, training, and validation

Due to the heterogeneity of the images—arising from diverse initial conditions, particularly variations in intensity and network-formation behavior observed in the time-lapse videos—it was not feasible to define a single generic segmentation method applicable to all conditions analyzed in this study. Segmentation was therefore performed using a deep learning model trained on ground-truth annotations generated with a combination of image-processing tools and state-of-the-art models, each selected according to the image conditions in which they performed best.

Our dataset was first divided into two subsets. The first subset was used to construct ground-truth segmentation data for training a deep learning–based model and validating its performance. The second subset was reserved for testing the model and assessing its generalization to unseen data.

To generate the ground-truth segmentation data for the first subset, six phase-reconstructed time-lapse videos of ∼1500 frame each were used, representing different conditions of epithelial cell growth resulting in different network morphologies (referred to as conditions 1–6). Several segmentation methods were explored, which can be grouped into five segmentation strategies. These states encompass morphological operations, classical image-processing algorithms, and recent deep-learning–based approaches. Specifically, we employed Otsu’s thresholding ^271^, the ERnet model ^101^, an adaptive thresholding algorithm ^272,273^, and the Contrast Limited Adaptive Histogram Equalization (CLAHE) method ^274^. The ERnet model used in this work is based on a pre-trained SwinIR network ^101^, available on the corresponding GitHub repository (https://github.com/charlesnchr/ERnet-v2/releases/tag/v2.0).

For two conditions, CLAHE was applied to enhance image contrast. Adaptive thresholding was also used, computing the threshold for each pixel based on the mean intensity of its local neighborhood minus a constant *C*. The neighborhood size was set to 349 pixels and *C* = 3. These values were selected after extensive visual inspection to identify the parameter combination yielding the most reliable segmentation quality across the evaluated images.

**Supplementary Figure S2.**
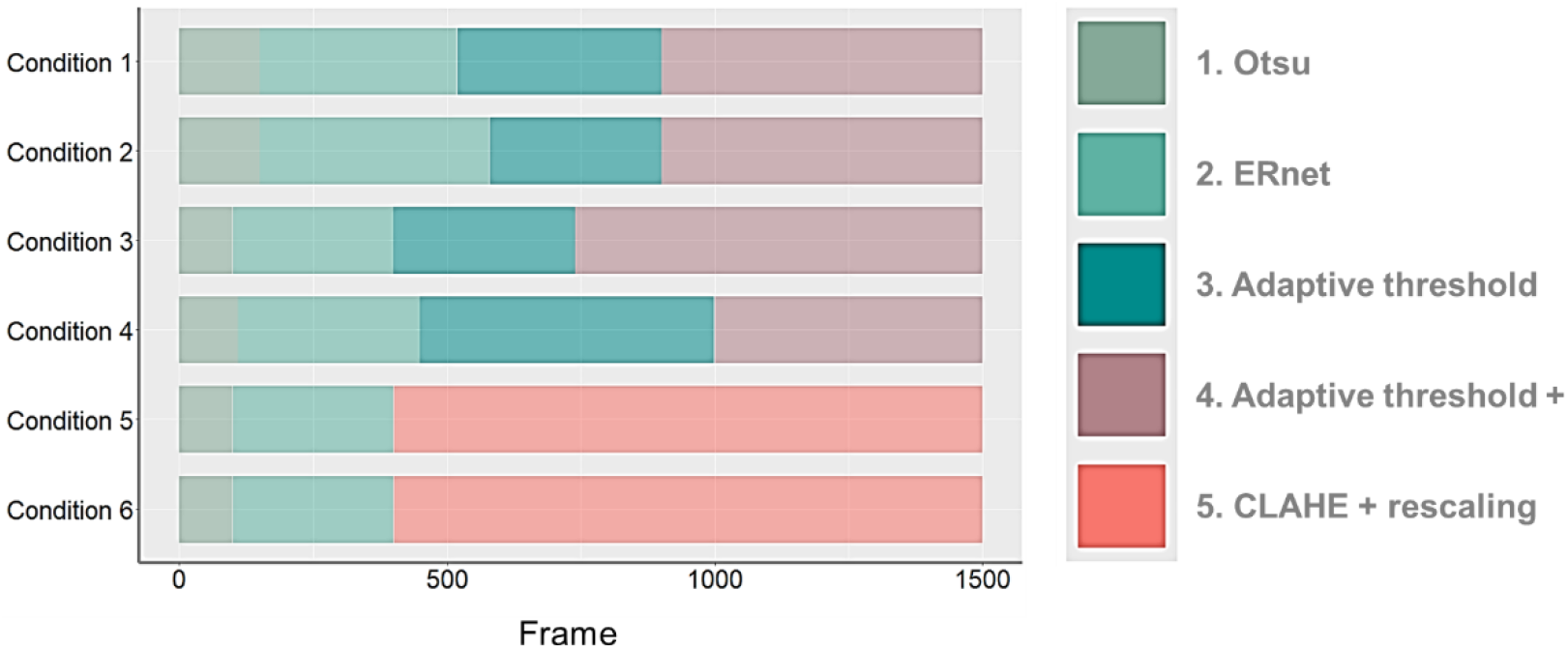
Implementation of segmentation strategies. Six phase-reconstructed time-lapse videos (∼1500 frames per condition; conditions 1–6) were analyzed using five segmentation strategies.

The five segmentation strategies are defined as follows:

1. **Otsu:** Images are normalized using the min–max method and segmented using Otsu’s threshold.
2. **ERnet:** Images are downsampled to half their original resolution, filtered bilaterally, normalized, rescaled, segmented using the pre-trained ERnet model, post-processed with a closing operation, and finally upsampled to their original size.
3. **Adaptive threshold:** Images undergo min–max normalization and intensity rescaling, followed by adaptive thresholding, removal of isolated pixels, and three dilation steps.
4. **Adaptive threshold+:** A bilateral filter is applied prior to normalization and rescaling; adaptive thresholding is followed by isolated-pixel removal, three dilations, and three closing operations.
5. **CLAHE + rescaling:** Images are filtered bilaterally, normalized, enhanced using CLAHE, segmented using Otsu’s threshold, and post-processed with an opening operation.

These segmentation strategies were applied to each video according to a frame-dependent scheme designed to accommodate the temporal evolution of image quality and network-formation dynamics **(Supplementary Figure S2).**

To prepare the input data for model training, each selected image was cropped into non-overlapping patches of size 512 × 512 pixels, yielding 48 crops per image. The model’s performance, however, was evaluated on the original full-resolution images. In total, 1050 training images and 210 validation images were selected, resulting in 50 400 cropped training samples and 10 080 cropped validation samples.

The segmentation model employed in this study is based on the U-Net architecture ^98^, a widely used convolutional neural network for biomedical image segmentation. An overview of the architecture is shown in the inset of **Figure 2A**. U-Net consists of two main components: an encoder that compresses the input image into a lower-dimensional latent representation while preserving relevant features, and a decoder that reconstructs a segmentation map at the original resolution. The encoder follows a contracting path composed of convolutional layers with ReLU activations and max-pooling operations that progressively reduce spatial resolution. The decoder follows an expansive path consisting of transposed convolutions and convolutional layers with ReLU activations, gradually restoring spatial resolution. At each decoding stage, feature maps from the corresponding encoder layer are concatenated to preserve spatial information. A sigmoid activation function is applied to the final output layer to produce a binary segmentation mask.

Model training was performed using the Binary Cross-Entropy (BCE) loss function, defined as (1):

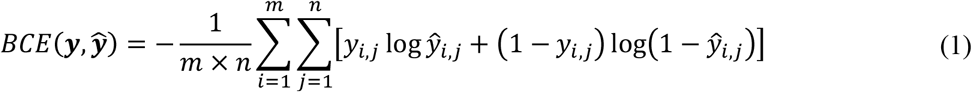

where *m* × *n* is the image size in pixels, *y* = (*y*_1,1_, …, *y_m_*_,*n*_) is the ground-truth image segmentation mask, and *y* = (*y*_1,1_, …, *y_m_*_,*n*_) is the predicted mask.

The model was optimized using the AdamW algorithm ^99^ with a learning rate of 1 × 10^−3^ and a weight decay of 5 × 10^−4^. An exponential learning-rate scheduler was applied to improve parameter updates during training. Mini-batches of size 8 were used. Early stopping was implemented: training was halted if the validation loss did not improve for 10 consecutive epochs. The model was trained for up to 100 epochs, and the version achieving the lowest validation loss was retained as the final model.

#### Graph construction: skeletonization and pruning

Once the images were segmented, the next step consisted of converting them into graph structures. The network was represented as an undirected and unweighted graph *G* = (*V*, *E*), where *V* denotes the set of nodes and *E* the set of edges. To obtain the graph topology, the segmented images were skeletonized using the fast parallel thinning algorithm ^275^. This algorithm iteratively removes boundary pixels from binary shapes while preserving their connectivity, applying two sub-iterations that evaluate local pixel neighborhoods to ensure that endpoints and essential structural elements are retained.

Graphs were then constructed from the resulting skeletons by identifying nodes and edges as the key topological components describing network connectivity. Graph-theoretical tools were subsequently applied to extract structural features and compute metrics characterizing the network architecture. As illustrated in **Figures 2A**, **3A**, and **5A**, skeleton-derived graphs may contain a large number of nodes, which can introduce substantial noise into the temporal evolution of the computed metrics. To mitigate this effect, the graph was simplified through a pruning procedure designed to preserve its essential topology.

Inspired by topological simplification approaches addressing intersection overcounting ^276^ and by angle-based filtering strategies for mesh preservation ^277^, we developed the pruning algorithm described here. The procedure begins by removing all nodes with degree less than 2—where the degree denotes the number of neighbors connected to a given node—thereby eliminating minimally connected elements. Next, nodes with degree-2 are examined together with their neighbors; the intermediate node is removed and its neighbors are directly connected unless this operation would distort the main network structure. To prevent such distortions, the angle *θ* formed between the node to be removed and its two neighbors is computed from their spatial coordinates. Following the criterion of Viana et al. ^277^, nodes are retained when θ ∈ (30^∘^, 150^∘^) and removed otherwise. Finally, any isolated nodes remaining after pruning are discarded.

#### Graph analysis: metrics definition

Quantitative network metrics were computed using the functions available in the Python package NetworkX ^97^. Among all available graph metrics, this study focuses on the most relevant ones for describing and characterizing the behaviours observed during epithelial network formation under the different conditions considered.

For a graph *G* = (*V*, *E*), the number of nodes and edges are denoted by |*V*| and |*E*|, respectively. The local connectivity of the network was quantified by the *mean node degree*, ⟨*k*⟩, defined as the average number of edges incident on a node. For a graph containing |*V*| nodes and |*E*| edges, the degree *k_i_* of node *i* was determined as the number of connected neighboring nodes, and the mean degree was calculated as (2):

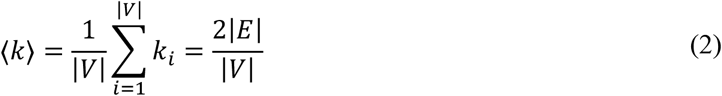

where the second equality follows from the handshaking lemma. Thus, ⟨*k*⟩ provides a global measure of network connectivity independent of graph size. In planar epithelial networks, ⟨*k*⟩ distinguishes tree-like architectures, characterized by values approaching 2, from increasingly reticulated networks enriched in cycles and branching junctions, for which ⟨k⟩ approaches 3. Node degrees and mean degree values were computed directly from graph representations generated by EpiNet using the NetworkX library. Furthermore, the distribution of nodes per degree provides insight into how network connectivity evolves. An increase in high degree nodes may indicate growing centralization, whereas an increase in low degree nodes may suggest a more decentralized or fragmented structure. This behavior is similarly reflected in the percentage of nodes per degree, which is directly related to the absolute number of nodes per degree.

To characterize the structural properties of the networks, the *average clustering coefficient* and the *assortativity coefficient* were evaluated. The average clustering coefficient quantifies the tendency of a network to form triangles by relating triplets of nodes to one another ^101^. A *closed triplet* consists of three nodes connected by three edges, whereas an *open triplet* consists of three nodes connected by only two edges ^278^. The average clustering coefficient (AClc) is defined as (3):

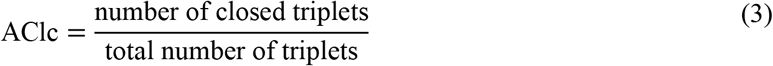

Network assortativity describes the preference of nodes to connect with others that share similar properties—here, specifically, node degree. Conversely, nodes that preferentially connect with dissimilar nodes exhibit *disassortative mixing*. The assortativity coefficient is defined as (4):

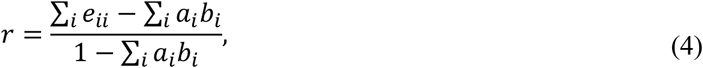

where *e_ij_* is the fraction of edges connecting nodes of type *i* and *j*, *a_i_* is the sum of *e_ij_* over all *j*, and *b_i_* is the sum of *e_ij_* over all *i* ^101^.

To assess macroscopic network organization, the number of *connected components* in the network was computed. Networks may be fully connected or composed of multiple disconnected components ^279^. These components can reflect merging or splitting behavior in evolving networks over time. In networks with many components, the largest connected component typically captures the most representative topological features ^280^. To account for connectivity patterns, two additional metrics related to the largest connected component *Gcc*_0_ of the network were included: the node-assembly ratio and the edge-assembly ratios. The *node-assembly ratio* is defined in (5), where |*V_Gcc_*_0_ | is the number of nodes in *Gcc*_0_. Equation (6) defines the *edge-assembly ratio*, where |*E_Gcc_*_0_ | is the number of edges in *Gcc*_0_. Both metrics range from 0 (completely disconnected) to 1 (fully connected).

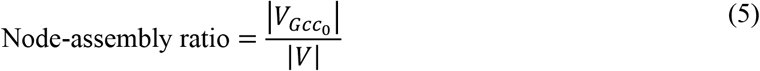

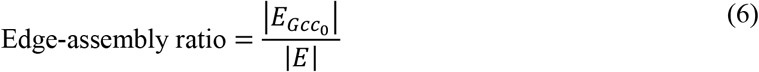

Global and local efficiency measures were used to evaluate higher-order transport efficiency in multicellular networks ^281^. *Local efficiency* quantifies how random node failures affect the number of shortest paths between neighboring node pairs, thereby providing insight into the network’s resilience to random errors through changes in local transportation costs ^281^. *Global efficiency*, in contrast, measures the overall routing efficiency of the network by considering the shortest paths between all node pairs and assessing how easily information can traverse the network ^281–283^.

The efficiency between a pair of nodes is defined as the multiplicative inverse of the shortest path distance between them. The *global efficiency E_glob_* of an undirected graph *G* = (*V*, *E*) is given by (7):

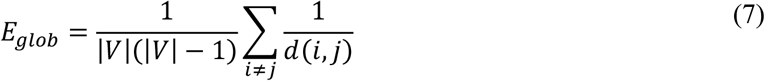

where *d*(*i*, *j*) is the shortest path distance between nodes *i* and *j*, and |*V*| is the total number of nodes in *G*.

Then, we analyzed the behavior of the connected components of the graph. Specifically, we computed the *cyclomatic number*, defined as the number of independent cycles in a graph and commonly used as a measure of *cyclic complexity* in mesh structures.

Let *H* be a connected component of the graph *G* = (*V*, *E*). For this component, the cyclomatic number is defined as (8):

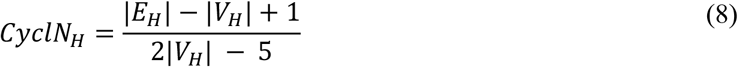

where |*V_H_*|, |*E_H_*| denote the number of nodes and edges of *H*, respectively.

This metric can be normalized using a reference value, which in this case corresponds to the largest connected component of the graph. Thus, we define the normalized cyclomatic number of a connected component *H*as (9):

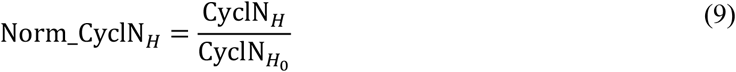

where CyclN*_H_*_0_ is the cyclomatic number of the largest connected component *H*_0_.

In addition to these metrics, we also consider the distribution of cycles within the largest connected component. Let *γ_i_* be the *i*-th cycle of *H*_0_, defined as (10):

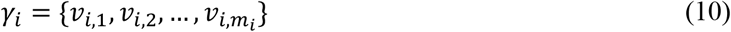

where *v_i_*_,*j*_ denotes the *j*-th node (with *j* ∈ {1, …, *m_i_*}) of the *i*-th cycle in *H*_0_.

We define the total number of cycle nodes in the largest connected component *H*_0_as (11):

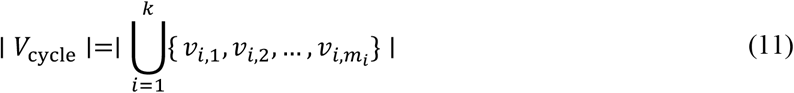

where *k* is the number of cycles in *H*_0_.

Each cycle *γ_i_* contributes the following set of edges (12):

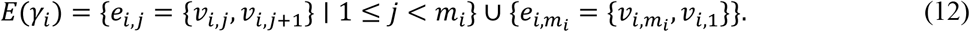

Then, the total number of cycle edges in *H*_0_ is (13):

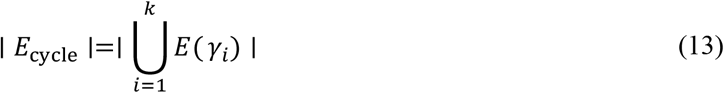

Finally, we compute the ratios of these two metrics with respect to the size of the largest connected component (14,15):

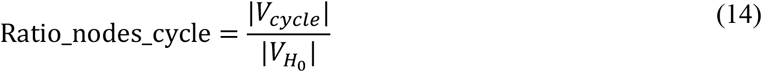

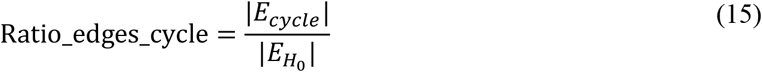

where |*V_H_*_0_ |, |*E_H_*_0_ | denote the number of nodes and edges of the largest connected component *H*_0_.

*Cycle birth statistics* were quantified using an edge-length–ordered filtration of EN planar graphs, where edge lengths were used to define the filtration sequence, while the graph topology remained unweighted. Edge lengths were computed from Euclidean distances between node coordinates and rescaled to physical units using the imaging calibration. For each network, edges were sorted in ascending order of length and sequentially inserted into an initially disconnected graph. At each insertion step, connectivity was updated and an edge was classified as cycle-forming when it connected two nodes already belonging to the same connected component, i.e., when it closed a loop in the growing graph. The corresponding edge length was recorded as the cycle birth length, providing a direct measure of the spatial scale at which mesoscopic loops emerge during network assembly. The resulting distributions were pooled across all networks within each experimental condition and analyzed statistically. This procedure is formally equivalent to a Kruskal-type minimum spanning tree filtration and closely related to the computation of 1-dimensional persistent homology in weighted graphs, where edge-insertion events correspond to birth of H1 topological features in a filtration parameterized by edge length ^284,285^. To account for potential multimodality reflecting distinct structural scales, the pooled cycle birth length distributions were modeled using a two-component Gaussian mixture model (G = 2), from which component means, variances, and mixture weights were extracted to distinguish local rearrangements from mesoscopic space-partitioning cycles **(Figure 5E)**.

### Geometrical analysis of epithelial networks

The relaxation of epithelial networks (ENs) was analyzed geometrically by quantifying the regularity of the network and the uniformity of space partitioning over time. The angle-filtered simplified graphs (see above) were used.

#### Quantification of network regularity by shape index p

For each network image, enclosed mesh domains were identified from binary masks after morphological preprocessing and particle segmentation in Fiji. The geometry of each mesh was represented as a polygon, from which area *A*, perimeter *P*, centroid position, and convexity related metrics such as circularity and solidity were calculated. The shape index *p* was then calculated as *p* = *P*/√*A*. Mean values and standard deviations of *p* distributions were computed for each time point and experimental condition.

#### Quantification of space partitioning by modified dimensionless quantizer energy dEq^∗^

To characterize the efficiency and uniformity of space partitioning by epithelial networks (ENs), we computed the modified dimensionless quantizer energy, (*dEq*^∗^), following the framework developed for dimensionless quantizer energy *dEq* based on Voronoi tessellations and centroidal Voronoi partitions ^107,127,129,130,237–239,286^. Modified in our case means that *dEq* was calculated for space partitioning by ENs and not by CVT. To compute *dEq*^∗^, second moments of EN polygonal mesh domains were calculated.

For a given mesh (*i*), the quantizer energy (*E_i_*) was defined as the polar second moment of area about its centroid, equivalent to the moment of inertia of the domain (16):

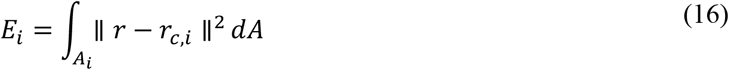

where *A_i_* is the mesh area and *r_c_*_,*i*_ is the centroid position. The integral was evaluated from polygon vertex coordinates using Green’s theorem. The total quantizer energy of the network was then obtained as (17):

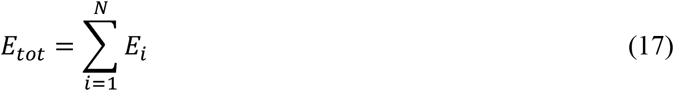

To enable comparison between networks containing different numbers of meshes and different total partitioned areas, a dimensionless quantizer energy was calculated as (18) ^130^:

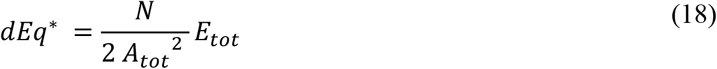

where (*N*) is the number of mesh domains and (*A_tot_*) is the total analyzed partitioned area. This normalization corresponds to the dimensionless quantizer energy *dEq* commonly used in the analysis of Voronoi tessellations and centroidal Voronoi tessellations (CVTs), for which lower values indicate more efficient and more uniform space partitioning ^127,130,286^. In particular, Poisson–Voronoi tessellations exhibit *dE_q_* ≈ 0.12, whereas increasingly ordered CVTs approach a near-hexagonal hyperuniform lattice with *p*_0_ ≈ 3.72 and dimensionless quantizer energy *dE_q_* ≈ 0.0808, close to the value *dE_q_* ≈ 0.080187 of hexagonal lattice ^107,130,240^.

### Surface Evolver modeling of epithelial network relaxation

To model the relaxation dynamics of epithelial networks (ENs), experimentally reconstructed graph geometries were mapped onto a surface-tension–driven mechanical framework implemented in the Surface Evolver STRING model, which performs energy minimization via gradient descent of interfacial line tension.

#### Graph preprocessing and construction of simulation geometries

Simplified angle-filtered graphs (see above) were used for simulations. The network geometries were further processed by:

1. Frame clipping: edges intersecting the boundaries of the defined frame were truncated at the frame limits to avoid artificial boundary artifacts in subsequent mechanical relaxation.
2. Node and edge extraction: intersection points and connectivity relations were retained as vertex and edge sets defining a planar graph representation of the EN.
3. Coordinate reconstruction: vertex coordinates were exported in Cartesian form and converted into Surface Evolver-compatible .fe files, encoding the initial network geometry as a tension-bearing polyhedral network.

#### STRING model implementation

Simulations were performed using the standard STRING model in Surface Evolver ^180,181^, in which all edges are assigned identical line tension. The system evolves according to (19):

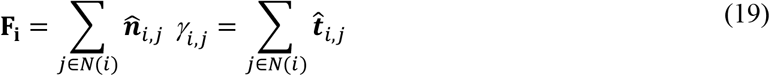

where **F_i_** is the force vector acting on a node (*i*) (the vector sum), *γ_i_*_,*j*_ is a scalar interfacial (line)-tension coefficient (constant and identical in our case), and *t_i_*_,*j*_ is a tension vector. Network configurations are updated via gradient descent of total interfacial energy, leading to progressive reduction of local force imbalance and interfacial length.

No additional bulk energy, area constraint, or perimeter elasticity was imposed, allowing the system to evolve purely under line-tension equilibration. Within this framework, Surface Evolver simulations represent, therefore, a pure tension-equilibration null model.

#### Extraction of simulated network observables

At each iteration of the Surface Evolver relaxation, network configurations were exported and reconstructed as planar graphs. From these configurations, geometric and topological observables were computed, including: vertex connectivity, mesh partitioning, and polygonal domain geometry. To quantify the efficiency of space partitioning, the modified dimensionless quantizer energy, (*dEq*^∗^), was computed for each configuration as described above.

#### Analysis of STRING model performance

To compare STRING simulations with experimentally observed relaxation dynamics, gradient-descent iteration steps were rescaled to physical time using a least-squares alignment procedure. This mapping minimized the discrepancy between simulated and experimentally measured evolution of *dEq*^∗^(*t*), enabling direct comparison of relaxation kinetics across both domains. The predictive performance of STRING simulations was assessed by comparing simulated and experimental network geometries using a *graph overlap* metric, evaluated as a function of simulation horizon and initialization time. The metric was computed by pairwise comparison of network binary images using the AND operator in the Image Calculator tool of Fiji.

### Statistical testing and reproducibility

For comparison between two groups, either the nonparametric Mann–Whitney U test or a two-tailed paired Student’s t-test was used, depending on normality and pairing structure. For comparisons involving more than two groups, a one-way analysis of variance (ANOVA) followed by Tukey–Kramer post hoc pairwise comparisons were performed.

To compare shape index *p* probability distributions between epithelial networks and Poisson–Voronoi tessellations, confidence intervals and significance estimates were obtained by nonparametric bootstrap resampling. Bootstrap procedures were also used where appropriate to estimate uncertainty in derived network metrics and fitted parameters. In particular, bootstrap-generated probability density functions were used to build moment–ratio diagrams, enabling comparison of EN and Poisson–Voronoi statistics in a reduced moment space. These diagrams were additionally used to evaluate deviations from canonical distribution classes, with approximate theoretical boundaries for log-normal and gamma distributions overlaid for reference.

To quantify treatment-induced effects on single cell morphology, Wasserstein distances ^157^ were computed between empirical distributions of single-cell shape parameters under control and inhibitor conditions. This metric was used to quantify global distributional shifts and to cluster inhibitors based on their multivariate morphological signatures.

Relationships between continuous variables (e.g., decay rates, scaling laws, and fitted parameters extracted from temporal or spatial summaries of network evolution) were evaluated using linear, exponential, or sigmoidal regression models. Goodness of fit and monotonic associations were further assessed using Pearson or Spearman correlation coefficients, depending on distributional assumptions and linearity. Statistical analyses and numerical computations were performed in R and Python.

## ACKNOWLEDGMENTS

We thank C. Allier (Janelia Research Campus, Ashburn, VA, USA) for initiating the analysis of cell-cell interactions using graph theory and for insightful discussions during the early stages of this work. We thank V. Collin-Faure (CEA/DRF/IRIG/DIESE/CBM/PROMIT, France) for assistance with HPDE-bic cell sorting and S. Combe (CEA/DRF/IRIG/BGE/BIOMICS, France) for assistance with confocal imaging. We thank the MuLife microscopy facility at IRIG/DBSCI, supported by CEA Nanobio and LabEx GRAL (ANR-10-LABX-49-01) and funded within the Université Grenoble Alpes graduate school CBH-EUR-GS (ANR-17-EURE-0003). We thank B. C. Campbell (Helen and Robert Appel Alzheimer’s Disease Research Institute, Weill Cornell Medicine, New York, NY, USA) for providing the pcDNA3.1-mGreenLantern plasmid.

## Author contributions

Conceptualization: M.Y.B., R.T.Z., G.G.,

Methodology: R.T.Z., A.B., F.M., A.N., S.A., L.H., G.G., M.Y.B.

Investigation: R.T.Z., A.B., F.M., L.H., G.G., M.Y.B.

Software: R.T.Z., L.H., G.G.

Formal analysis: R.T.Z., A.B., A.N., M.Y.B.

Supervision: G.G., L.H., S.A., A.N., X.G., M.Y.B.

Project administration: X.G., M.Y.B.

Funding acquisition: G.G., S.A., A.N., X.G., M.Y.B.

Writing—original draft: R.T.Z., A.B., A.N., L.H., G.G., M.Y.B.

Writing—review & editing:

## Data, code, and materials availability

The main data and code required to evaluate and reproduce the results presented in this paper are provided in the paper and/or the Supplementary Materials, or are described in previously published works appropriately cited in the article.

## REFERENCES

(1) Goodwin, K.; Nelson, C. M. Branching Morphogenesis. Development 2020, 147 (10), dev184499. 10.1242/dev.184499.

(2) Hannezo, E.; Scheele, C. L. G. J.; Moad, M.; Drogo, N.; Heer, R.; Sampogna, R. V.; Van Rheenen, J.; Simons, B. D. A Unifying Theory of Branching Morphogenesis. Cell 2017, 171 (1), 242–255.e27. 10.1016/j.cell.2017.08.026.

(3) Hunt, D.; Savage, V. M. Asymmetries Arising from the Space-Filling Nature of Vascular Networks. Phys. Rev. E 2016, 93 (6), 062305. 10.1103/PhysRevE.93.062305.

(4) Iber, D.; Menshykau, D. The Control of Branching Morphogenesis. Open Biol. 2013, 3 (9), 130088. 10.1098/rsob.130088.

(5) Katifori, E.; Szöllősi, G. J.; Magnasco, M. O. Damage and Fluctuations Induce Loops in Optimal Transport Networks. Phys. Rev. Lett. 2010, 104 (4), 048704. 10.1103/PhysRevLett.104.048704.

(6) Paramore, S. V.; Goodwin, K.; Nelson, C. M. How to Build an Epithelial Tree. Phys. Biol. 2022, 19 (6), 061002. 10.1088/1478-3975/ac9e38.

(7) Ronellenfitsch, H.; Katifori, E. Phenotypes of Vascular Flow Networks. Phys. Rev. Lett. 2019, 123 (24), 248101. 10.1103/PhysRevLett.123.248101.

(8) Ronellenfitsch, H.; Katifori, E. Global Optimization, Local Adaptation, and the Role of Growth in Distribution Networks. Phys. Rev. Lett. 2016, 117 (13), 138301. 10.1103/PhysRevLett.117.138301.

(9) Uçar, M. C.; Kamenev, D.; Sunadome, K.; Fachet, D.; Lallemend, F.; Adameyko, I.; Hadjab, S.; Hannezo, E. Theory of Branching Morphogenesis by Local Interactions and Global Guidance. Nat Commun 2021, 12 (1), 6830. 10.1038/s41467-021-27135-5.

(10) West, G. B.; Enquist, B. J. The Origin of Universal Scaling Laws In. Scaling in biology 2000, 87.

(11) Yu, W.; Marshall, W. F.; Metzger, R. J.; Brakeman, P. R.; Morsut, L.; Lim, W.; Mostov, K. E. Simple Rules Determine Distinct Patterns of Branching Morphogenesis. Cell Systems 2019, 9 (3), 221–227. 10.1016/j.cels.2019.08.001.

(12) Pries, A. R.; Secomb, T. W. Making Microvascular Networks Work: Angiogenesis, Remodeling, and Pruning. Physiology 2014, 29 (6), 446–455. 10.1152/physiol.00012.2014.

(13) Bullmore, E.; Sporns, O. The Economy of Brain Network Organization. Nature reviews neuroscience 2012, 13 (5), 336–349.

(14) Kaiser, F.; Ronellenfitsch, H.; Witthaut, D. Discontinuous Transition to Loop Formation in Optimal Supply Networks. Nat Commun 2020, 11 (1), 5796. 10.1038/s41467-020-19567-2.

(15) Lu, Y.; Hu, D. Optimisation of Biological Transport Networks. EAJAM 2021, 12 (1), 72–95. 10.4208/eajam.180521.130721.

(16) Mengistu, H.; Huizinga, J.; Mouret, J.-B.; Clune, J. The Evolutionary Origins of Hierarchy. PLoS Comput Biol 2016, 12 (6), e1004829. 10.1371/journal.pcbi.1004829.

(17) Murray, C. D. The Physiological Principle of Minimum Work: I. The Vascular System and the Cost of Blood Volume. Proceedings of the National Academy of Sciences 1926, 12 (3), 207–214.

(18) Tekin, E.; Hunt, D.; Newberry, M. G.; Savage, V. M. Do Vascular Networks Branch Optimally or Randomly across Spatial Scales? PLoS Comput Biol 2016, 12 (11), e1005223. 10.1371/journal.pcbi.1005223.

(19) Ghabrial, A.; Luschnig, S.; Metzstein, M. M.; Krasnow, M. A. Branching Morphogenesis of the Drosophila Tracheal System. Annu. Rev. Cell Dev. Biol. 2003, 19 (1), 623–647. 10.1146/annurev.cellbio.19.031403.160043.

(20) Carmeliet, P.; Jain, R. K. Molecular Mechanisms and Clinical Applications of Angiogenesis. Nature 2011, 473 (7347), 298–307. 10.1038/nature10144.

(21) Tanimizu, N.; Kaneko, K.; Itoh, T.; Ichinohe, N.; Ishii, M.; Mizuguchi, T.; Hirata, K.; Miyajima, A.; Mitaka, T. Intrahepatic Bile Ducts Are Developed through Formation of Homogeneous Continuous Luminal Network and Its Dynamic Rearrangement in Mice. Hepatology 2016, 64 (1), 175–188. 10.1002/hep.28521.

(22) Jackson, A. L.; Heilmann, S.; Agerskov, R.; Ebeid, C.; Krivokapic, J. M.; Romero Herrera, J. A.; Semb, H.; Nyeng, P. Real-Time Imaging Reveals New Mechanisms for Pancreatic Ductal Establishment and Remodeling. Journal of Cell Biology 2026, 225 (3), e202409022.

(23) Dahl-Jensen, S. B.; Yennek, S.; Flasse, L.; Larsen, H. L.; Sever, D.; Karremore, G.; Novak, I.; Sneppen, K.; Grapin-Botton, A. Deconstructing the Principles of Ductal Network Formation in the Pancreas. PLoS Biol 2018, 16 (7), e2002842. 10.1371/journal.pbio.2002842.

(24) Kazenwadel, J.; Harvey, N. L. Morphogenesis of the Lymphatic Vasculature: A Focus on New Progenitors and Cellular Mechanisms Important for Constructing Lymphatic Vessels. Developmental Dynamics 2016, 245 (3), 209–219. 10.1002/dvdy.24313.

(25) LaRue, A. C.; Mironov, V. A.; Argraves, W. S.; Czirók, A.; Fleming, P. A.; Drake, C. J. Patterning of Embryonic Blood Vessels. Developmental Dynamics 2003, 228 (1), 21–29. 10.1002/dvdy.10339.

(26) Hahn, A.; Bode, J.; Krüwel, T.; Solecki, G.; Heiland, S.; Bendszus, M.; Tews, B.; Winkler, F.; Breckwoldt, M. O.; Kurz, F. T. Glioblastoma Multiforme Restructures the Topological Connectivity of Cerebrovascular Networks. Sci Rep 2019, 9 (1), 11757. 10.1038/s41598-019-47567-w.

(27) Banavar, J. R.; Colaiori, F.; Flammini, A.; Maritan, A.; Rinaldo, A. Topology of the Fittest Transportation Network. Phys. Rev. Lett. 2000, 84 (20), 4745–4748. 10.1103/PhysRevLett.84.4745.

(28) Durand, M. Structure of Optimal Transport Networks Subject to a Global Constraint. Phys. Rev. Lett. 2007, 98 (8), 088701. 10.1103/PhysRevLett.98.088701.

(29) Corson, F. Fluctuations and Redundancy in Optimal Transport Networks. Phys. Rev. Lett. 2010, 104 (4), 048703. 10.1103/PhysRevLett.104.048703.

(30) Bohn, S.; Magnasco, M. O. Structure, Scaling, and Phase Transition in the Optimal Transport Network. Phys. Rev. Lett. 2007, 98 (8), 088702. 10.1103/PhysRevLett.98.088702.

(31) Uçar, M. C.; Hannezo, E.; Tiilikainen, E.; Liaqat, I.; Jakobsson, E.; Nurmi, H.; Vaahtomeri, K. Self-Organized and Directed Branching Results in Optimal Coverage in Developing Dermal Lymphatic Networks. Nat Commun 2023, 14 (1), 5878. 10.1038/s41467-023-41456-7.

(32) Lang, C.; Conrad, L.; Iber, D. Organ-Specific Branching Morphogenesis. Front. Cell Dev. Biol. 2021, 9, 671402. 10.3389/fcell.2021.671402.

(33) Metzger, R. J.; Klein, O. D.; Martin, G. R.; Krasnow, M. A. The Branching Programme of Mouse Lung Development. Nature 2008, 453 (7196), 745–750.

(34) Raj, A.; Chen, Y. The Wiring Economy Principle: Connectivity Determines Anatomy in the Human Brain. PloS one 2011, 6 (9), e14832.

(35) Ahn, Y.-Y.; Jeong, H.; Kim, B. J. Wiring Cost in the Organization of a Biological Neuronal Network. Physica A: Statistical Mechanics and its Applications 2006, 367, 531–537.

(36) Waszkiewicz, R.; Shaw, J. B.; Lisicki, M.; Szymczak, P. Goldilocks Fluctuations: Dynamic Constraints on Loop Formation in Scale-Free Transport Networks. Phys. Rev. Lett. 2024, 132 (13), 137401. 10.1103/PhysRevLett.132.137401.

(37) Folkman, J.; Haudenschild, C. Angiogenesis in Vitro. Nature 1980, 288 (5791), 551–556. 10.1038/288551a0.

(38) Harris, A. K.; Stopak, D.; Wild, P. Fibroblast Traction as a Mechanism for Collagen Morphogenesis. Nature 1981, 290 (5803), 249–251. 10.1038/290249a0.

(39) Kleinman, H. K.; Martin, G. R. Matrigel: Basement Membrane Matrix with Biological Activity; Elsevier, 2005; Vol. 15, pp 378–386.

(40) Kubota, Y.; Kleinman, H. K.; Martin, G. R.; Lawley, T. J. Role of Laminin and Basement Membrane in the Morphological Differentiation of Human Endothelial Cells into Capillary-like Structures. The Journal of cell biology 1988, 107 (4), 1589–1598. 10.1083/jcb.107.4.1589.

(41) Madri, J. A.; Williams, S. K. Capillary Endothelial Cell Cultures: Phenotypic Modulation by Matrix Components. The Journal of cell biology 1983, 97 (1), 153–165.

(42) Méhes, E.; Biri-Kovács, B.; Isai, D. G.; Gulyás, M.; Nyitray, L.; Czirók, A. Matrigel Patterning Reflects Multicellular Contractility. PLoS Comput Biol 2019, 15 (10), e1007431. 10.1371/journal.pcbi.1007431.

(43) Montesano, R.; Soriano, J. V.; Fialka, I.; Orci, L. Isolation of EpH4 Mammary Epithelial Cell Subpopulations Which Differ in Their Morphogenetic Properties. In Vitro Cell.Dev.Biol.-Animal 1998, 34 (6), 468–477. 10.1007/s11626-998-0080-3.

(44) Montesano, R.; Schaller, G.; Orci, L. Induction of Epithelial Tubular Morphogenesis in Vitro by Fibroblast-Derived Soluble Factors. Cell 1991, 66 (4), 697–711.

(45) Montesano, R.; Orci, L.; Vassalli, P. In Vitro Rapid Organization of Endothelial Cells into Capillary-like Networks Is Promoted by Collagen Matrices. The Journal of cell biology 1983, 97 (5), 1648–1652. 10.1083/jcb.97.5.1648.

(46) Nakano, T.; Okaie, Y.; Kinugasa, Y.; Koujin, T.; Suda, T.; Hiraoka, Y.; Haraguchi, T. Roles of Remote and Contact Forces in Epithelial Cell Structure Formation. Biophysical Journal 2020, 118 (6), 1466–1478. 10.1016/j.bpj.2020.01.037.

(47) Rosines, E.; Johkura, K.; Zhang, X.; Schmidt, H. J.; DeCambre, M.; Bush, K. T.; Nigam, S. K. Constructing Kidney-like Tissues from Cells Based on Programs for Organ Development: Toward a Method of *In Vitro* Tissue Engineering of the Kidney. Tissue Engineering Part A 2010, 16 (8), 2441–2455. 10.1089/ten.tea.2009.0548.

(48) Royce, L. S.; Kibbey, M. C.; Mertz, P.; Kleinman, H. K.; Baum, B. J. Human Neoplastic Submandibular Intercalated Duct Cells Express an Acinar Phenotype When Cultured on a Basement Membrane Matrix. Differentiation 1993, 52 (3), 247–255.

(49) Sakurai, H.; Barros, E. J.; Tsukamoto, T.; Barasch, J.; Nigam, S. K. An in Vitro Tubulogenesis System Using Cell Lines Derived from the Embryonic Kidney Shows Dependence on Multiple Soluble Growth Factors. Proceedings of the National Academy of Sciences 1997, 94 (12), 6279–6284.

(50) Stopak, D.; Harris, A. K. Connective Tissue Morphogenesis by Fibroblast Traction: I. Tissue Culture Observations. Developmental biology 1982, 90 (2), 383–398.

(51) Vernon, R. B.; Sage, E. H. Between Molecules and Morphology. Extracellular Matrix and Creation of Vascular Form. 1995, 147 (4), 873–882.

(52) Vernon, R. B.; Angello, J. C.; Iruela-Arispe, M. L.; Lane, T. F.; Sage, E. H. Reorganization of Basement Membrane Matrices by Cellular Traction Promotes the Formation of Cellular Networks in Vitro. Lab Invest 1992, 66 (5), 536–547.

(53) Manoussaki, D.; Lubkin, S. R.; Vemon, R. B.; Murray, J. D. A Mechanical Model for the Formation of Vascular Networks in Vitro. Acta Biotheor 1996, 44 (3–4), 271–282. 10.1007/BF00046533.

(54) Murray, J.; Manoussaki, D.; Lubkin, S.; Vernon, R. A Mechanical Theory of in Vitro Vascular Network Formation. In Vascular morphogenesis: In vivo, in vitro, in mente; Springer, 1998; pp 173–188.

(55) Murray, J. D. Mathematical Biology II: Spatial Models and Biomedical Applications, 3rd ed.; Springer: Berlin, 2003; Vol. 18.

(56) Van Oers, R. F. M.; Rens, E. G.; LaValley, D. J.; Reinhart-King, C. A.; Merks, R. M. H. Mechanical Cell-Matrix Feedback Explains Pairwise and Collective Endothelial Cell Behavior In Vitro. PLoS Comput Biol 2014, 10 (8), e1003774. 10.1371/journal.pcbi.1003774.

(57) Bailles, A.; Gehrels, E. W.; Lecuit, T. Mechanochemical Principles of Spatial and Temporal Patterns in Cells and Tissues. Annu. Rev. Cell Dev. Biol. 2022, 38 (1), 321–347. 10.1146/annurev-cellbio-120420-095337.

(58) Van Helvert, S.; Storm, C.; Friedl, P. Mechanoreciprocity in Cell Migration. Nat Cell Biol 2018, 20 (1), 8–20. 10.1038/s41556-017-0012-0.

(59) Andrée, B.; Ichanti, H.; Kalies, S.; Heisterkamp, A.; Strauß, S.; Vogt, P.-M.; Haverich, A.; Hilfiker, A. Formation of Three-Dimensional Tubular Endothelial Cell Networks under Defined Serum-Free Cell Culture Conditions in Human Collagen Hydrogels. Scientific reports 2019, 9 (1), 5437.

(60) Arnaoutova, I.; George, J.; Kleinman, H. K.; Benton, G. The Endothelial Cell Tube Formation Assay on Basement Membrane Turns 20: State of the Science and the Art. Angiogenesis 2009, 12 (3), 267–274.

(61) Cimpean, A. M.; Raica, M. Historical Overview of in Vivo and in Vitro Angiogenesis Assays. Vascular Morphogenesis: Methods and Protocols 2020, 1–13.

(62) Iruela-Arispe, M. L.; Davis, G. E. Cellular and Molecular Mechanisms of Vascular Lumen Formation. Developmental cell 2009, 16 (2), 222–231.

(63) O’Connor, C.; Brady, E.; Zheng, Y.; Moore, E.; Stevens, K. R. Engineering the Multiscale Complexity of Vascular Networks. Nature Reviews Materials 2022, 7 (9), 702–716.

(64) Mederacke, M.; Conrad, L.; Doumpas, N.; Vetter, R.; Iber, D. Geometric Effects Position Renal Vesicles during Kidney Development. Cell reports 2023, 42 (12).

(65) Czirok, A.; Rongish, B. J.; Little, C. D. Vascular Network Formation in Expanding versus Static Tissues: Embryos and Tumors. Genes & Cancer 2011, 2 (12), 1072–1080. 10.1177/1947601911426774.

(66) Carpentier, G.; Berndt, S.; Ferratge, S.; Rasband, W.; Cuendet, M.; Uzan, G.; Albanese, P. Angiogenesis Analyzer for ImageJ—A Comparative Morphometric Analysis of “Endothelial Tube Formation Assay” and “Fibrin Bead Assay.” Scientific reports 2020, 10 (1), 11568.

(67) Sailem, H. Z.; Al Haj Zen, A. Morphological Landscape of Endothelial Cell Networks Reveals a Functional Role of Glutamate Receptors in Angiogenesis. Sci Rep 2020, 10 (1), 13829. 10.1038/s41598-020-70440-0.

(68) Gamba, A.; Ambrosi, D.; Coniglio, A.; Candia, A. de; Talia, S. D.; Giraudo, E.; Serini, G.; Preziosi, L.; Bussolino, F. Percolation, Morphogenesis, and Burgers Dynamics in Blood Vessels Formation. Phys. Rev. Lett. 2003, 90 (11), 118101. 10.1103/PhysRevLett.90.118101.

(69) Merks, R. M. H.; Perryn, E. D.; Shirinifard, A.; Glazier, J. A. Contact-Inhibited Chemotaxis in De Novo and Sprouting Blood-Vessel Growth. PLoS Comput Biol 2008, 4 (9), e1000163. 10.1371/journal.pcbi.1000163.

(70) Pereira, M.; Pinto, J.; Arteaga, B.; Guerra, A.; Jorge, R. N.; Monteiro, F. J.; Salgado, C. L. A Comprehensive Look at In Vitro Angiogenesis Image Analysis Software. IJMS 2023, 24 (24), 17625. 10.3390/ijms242417625.

(71) Merks, R. M. H.; Brodsky, S. V.; Goligorksy, M. S.; Newman, S. A.; Glazier, J. A. Cell Elongation Is Key to in Silico Replication of in Vitro Vasculogenesis and Subsequent Remodeling. Developmental Biology 2006, 289 (1), 44–54. 10.1016/j.ydbio.2005.10.003.

(72) Sun, J.; Jamilpour, N.; Wang, F.-Y.; Wong, P. K. Geometric Control of Capillary Architecture via Cell-Matrix Mechanical Interactions. Biomaterials 2014, 35 (10), 3273–3280. 10.1016/j.biomaterials.2013.12.101.

(73) Hanada, Y.; Halder, S.; Arima, Y.; Haruta, M.; Ogoh, H.; Ogura, S.; Shiraki, Y.; Nakano, S.; Ozeki, Y.; Fukuhara, S.; Uemura, A.; Murohara, T.; Nishiyama, K. Biomechanical Control of Vascular Morphogenesis by the Surrounding Stiffness. Nat Commun 2025, 16 (1), 6788. 10.1038/s41467-025-61804-z.

(74) Masson-Meyers, D. S.; Tayebi, L. Vascularization Strategies in Tissue Engineering Approaches for Soft Tissue Repair. J Tissue Eng Regen Med 2021, 15 (9), 747–762. 10.1002/term.3225.

(75) Serini, G.; Ambrosi, D.; Giraudo, E.; Gamba, A.; Preziosi, L.; Bussolino, F. Modeling the Early Stages of Vascular Network Assembly. EMBO J 2003, 22 (8), 1771–1779. 10.1093/emboj/cdg176.

(76) Ouyang, H.; Mou, L.; Luk, C.; Liu, N.; Karaskova, J.; Squire, J.; Tsao, M.-S. Immortal Human Pancreatic Duct Epithelial Cell Lines with Near Normal Genotype and Phenotype. The American Journal of Pathology 2000, 157 (5), 1623–1631. 10.1016/S0002-9440(10)64800-6.

(77) Broutier, L.; Andersson-Rolf, A.; Hindley, C. J.; Boj, S. F.; Clevers, H.; Koo, B.-K.; Huch, M. Culture and Establishment of Self-Renewing Human and Mouse Adult Liver and Pancreas 3D Organoids and Their Genetic Manipulation. Nat Protoc 2016, 11 (9), 1724–1743. 10.1038/nprot.2016.097.

(78) Greggio, C.; De Franceschi, F.; Figueiredo-Larsen, M.; Gobaa, S.; Ranga, A.; Semb, H.; Lutolf, M.; Grapin-Botton, A. Artificial Three-Dimensional Niches Deconstruct Pancreas Development in Vitro. Development 2013, 140 (21), 4452–4462. 10.1242/dev.096628.

(79) Mailleux, A. A.; Overholtzer, M.; Brugge, J. S. Lumen Formation during Mammary Epithelial Morphogenesis: Insights from in Vitro and in Vivo Models. Cell cycle 2008, 7 (1), 57–62.

(80) Randriamanantsoa, S. J.; Raich, M. K.; Saur, D.; Reichert, M.; Bausch, A. R. Coexisting Mechanisms of Luminogenesis in Pancreatic Cancer-Derived Organoids. Iscience 2024, 27 (7).

(81) Toivanen, R.; Shen, M. M. Prostate Organogenesis: Tissue Induction, Hormonal Regulation and Cell Type Specification. Development 2017, 144 (8), 1382–1398.

(82) Martens, S.; Coolens, K.; Van Bulck, M.; Arsenijevic, T.; Casamitjana, J.; Ruiz, A. F.; El Kaoutari, A.; de Villareal, J. M.; Madhloum, H.; Esni, F. Discovery and 3D Imaging of a Novel ΔNp63-Expressing Basal Cell Type in Human Pancreatic Ducts with Implications in Disease. Gut 2022, 71 (10), 2030–2042.

(83) Braga, V.; Pemberton, L.; Duhig, T.; Gendler, S. Spatial and Temporal Expression of an Epithelial Mucin, Muc-1, during Mouse Development. Development 1992, 115 (2), 427–437.

(84) Kopinke, D.; Murtaugh, L. C. Exocrine-to-Endocrine Differentiation Is Detectable Only Prior to Birth in the Uninjured Mouse Pancreas. BMC developmental biology 2010, 10 (1), 38.

(85) Bi, D.; Yang, X.; Marchetti, M. C.; Manning, M. L. Motility-Driven Glass and Jamming Transitions in Biological Tissues. Phys. Rev. X 2016, 6 (2), 021011. 10.1103/PhysRevX.6.021011.

(86) Bi, D.; Lopez, J. H.; Schwarz, J. M.; Manning, M. L. A Density-Independent Rigidity Transition in Biological Tissues. Nature Phys 2015, 11 (12), 1074–1079. 10.1038/nphys3471.

(87) Bi, D.; Lopez, J. H.; Schwarz, J. M.; Manning, M. L. Energy Barriers and Cell Migration in Densely Packed Tissues. Soft Matter 2014, 10 (12), 1885. 10.1039/c3sm52893f.

(88) Lemke, S. B.; Nelson, C. M. Dynamic Changes in Epithelial Cell Packing during Tissue Morphogenesis. Current Biology 2021, 31 (18), R1098–R1110.

(89) Park, J.-A.; Kim, J. H.; Bi, D.; Mitchel, J. A.; Qazvini, N. T.; Tantisira, K.; Park, C. Y.; McGill, M.; Kim, S.-H.; Gweon, B.; Notbohm, J.; Steward Jr, R.; Burger, S.; Randell, S. H.; Kho, A. T.; Tambe, D. T.; Hardin, C.; Shore, S. A.; Israel, E.; Weitz, D. A.; Tschumperlin, D. J.; Henske, E. P.; Weiss, S. T.; Manning, M. L.; Butler, J. P.; Drazen, J. M.; Fredberg, J. J. Unjamming and Cell Shape in the Asthmatic Airway Epithelium. Nature Mater 2015, 14 (10), 1040–1048. 10.1038/nmat4357.

(90) Staple, D. B.; Farhadifar, R.; Röper, J.-C.; Aigouy, B.; Eaton, S.; Jülicher, F. Mechanics and Remodelling of Cell Packings in Epithelia. Eur. Phys. J. E 2010, 33 (2), 117–127. 10.1140/epje/i2010-10677-0.

(91) Chieco, A. T.; Durian, D. J. Quantifying the Long-Range Structure of Foams and Other Cellular Patterns with Hyperuniformity Disorder Length Spectroscopy. Phys. Rev. E 2021, 103 (6), 062609. 10.1103/PhysRevE.103.062609.

(92) Damavandi, O. K.; Arzash, S.; Lawson-Keister, E.; Manning, M. L. Universality in the Mechanical Behavior of Vertex Models for Biological Tissues. PRX Life 2025, 3 (3), 033001. 10.1103/9ktk-6rqc.

(93) Allier, C.; Morel, S.; Vincent, R.; Ghenim, L.; Navarro, F.; Menneteau, M.; Bordy, T.; Hervé, L.; Cioni, O.; Gidrol, X. Imaging of Dense Cell Cultures by Multiwavelength Lens-free Video Microscopy. Cytometry Part A 2017, 91 (5), 433–442.

(94) Dolega, M. E.; Allier, C.; Vinjimore Kesavan, S.; Gerbaud, S.; Kermarrec, F.; Marcoux, P.; Dinten, J.-M.; Gidrol, X.; Picollet-D’Hahan, N. Label-Free Analysis of Prostate Acini-like 3D Structures by Lensfree Imaging. Biosensors and Bioelectronics 2013, 49, 176–183. 10.1016/j.bios.2013.05.001.

(95) Ghenim, L.; Allier, C.; Obeid, P.; Hervé, L.; Fortin, J.-Y.; Balakirev, M.; Gidrol, X. A New Ultradian Rhythm in Mammalian Cell Dry Mass Observed by Holography. Scientific Reports 2021, 11 (1). 10.1038/s41598-020-79661-9.

(96) Kesavan, S. V.; Momey, F.; Cioni, O.; David-Watine, B.; Dubrulle, N.; Shorte, S.; Sulpice, E.; Freida, D.; Chalmond, B.; Dinten, J. M.; Gidrol, X.; Allier, C. High-Throughput Monitoring of Major Cell Functions by Means of Lensfree Video Microscopy. Sci Rep 2014, 4 (1), 5942. 10.1038/srep05942.

(97) Hagberg, A.; Swart, P. J.; Schult, D. A. Exploring Network Structure, Dynamics, and Function Using NetworkX; Los Alamos National Laboratory (LANL), 2007.

(98) Ronneberger, O.; Fischer, P.; Brox, T. U-Net: Convolutional Networks for Biomedical Image Segmentation. In Medical Image Computing and Computer-Assisted Intervention – MICCAI 2015; Navab, N., Hornegger, J., Wells, W. M., Frangi, A. F., Eds.; Lecture Notes in Computer Science; Springer International Publishing: Cham, 2015; Vol. 9351, pp 234–241. 10.1007/978-3-319-24574-4_28.

(99) Loshchilov, I.; Hutter, F. Decoupled Weight Decay Regularization. arXiv preprint arXiv:1711.05101 2017.

(100) Ramos, D.; Franco-Pedroso, J.; Lozano-Diez, A.; Gonzalez-Rodriguez, J. Deconstructing Cross-Entropy for Probabilistic Binary Classifiers. Entropy 2018, 20 (3), 208.

(101) Lu, M.; Christensen, C. N.; Weber, J. M.; Konno, T.; Läubli, N. F.; Scherer, K. M.; Avezov, E.; Lio, P.; Lapkin, A. A.; Kaminski Schierle, G. S.; Kaminski, C. F. ERnet: A Tool for the Semantic Segmentation and Quantitative Analysis of Endoplasmic Reticulum Topology. Nat Methods 2023, 20 (4), 569–579. 10.1038/s41592-023-01815-0.

(102) Otsu, N. A Threshold Selection Method from Gray-Level Histograms. Automatica 1975, 11, 285–296.

(103) Renieblas, G. P.; Nogués, A. T.; González, A. M.; Gómez-Leon, N.; Del Castillo, E. G. Structural Similarity Index Family for Image Quality Assessment in Radiological Images. J. Med. Imag 2017, 4 (3), 035501. 10.1117/1.JMI.4.3.035501.

(104) Clark, J.; Holton, D. A. A First Look at Graph Theory; World Scientific: Teaneck, NJ, 1991.

(105) Diestel, R. Planar Graphs. In Graph Theory; Graduate Texts in Mathematics; Springer Berlin Heidelberg: Berlin, Heidelberg, 2017; Vol. 173, pp 89–118. 10.1007/978-3-662-53622-3_4.

(106) Baumgarten, W.; Ueda, T.; Hauser, M. J. B. Plasmodial Vein Networks of the Slime Mold *Physarum Polycephalum* Form Regular Graphs. Phys. Rev. E 2010, 82 (4), 046113. 10.1103/PhysRevE.82.046113.

(107) Du, Q.; Wang, D. The Optimal Centroidal Voronoi Tessellations and the Gersho’s Conjecture in the Three-Dimensional Space. Computers & Mathematics with Applications 2005, 49 (9–10), 1355–1373. 10.1016/j.camwa.2004.12.008.

(108) Gibson, M. C.; Patel, A. B.; Nagpal, R.; Perrimon, N. The Emergence of Geometric Order in Proliferating Metazoan Epithelia. Nature 2006, 442 (7106), 1038–1041. 10.1038/nature05014.

(109) Gibson, W. T.; Gibson, M. C. Chapter 4 Cell Topology, Geometry, and Morphogenesis in Proliferating Epithelia. In Current Topics in Developmental Biology; Elsevier, 2009; Vol. 89, pp 87–114. 10.1016/S0070-2153(09)89004-2.

(110) Honda, H. Description of Cellular Patterns by Dirichlet Domains: The Two-Dimensional Case. Journal of theoretical biology 1978, 72 (3), 523–543.

(111) Nagai, T.; Honda, H. A Dynamic Cell Model for the Formation of Epithelial Tissues. Philosophical Magazine B 2001, 81 (7), 699–719. 10.1080/13642810108205772.

(112) Rivier, N.; Schliecker, G.; Dubertret, B. The Stationary State of Epithelia. Acta Biotheor 1995, 43 (4), 403–423. 10.1007/BF00713562.

(113) Sánchez-Gutiérrez, D.; Sáez, A.; Pascual, A.; Escudero, L. M. Topological Progression in Proliferating Epithelia Is Driven by a Unique Variation in Polygon Distribution. PLoS ONE 2013, 8 (11), e79227. 10.1371/journal.pone.0079227.

(114) Sánchez-Gutiérrez, D.; Tozluoglu, M.; Barry, J. D.; Pascual, A.; Mao, Y.; Escudero, L. M. Fundamental Physical Cellular Constraints Drive Self-organization of Tissues. The EMBO Journal 2016, 35 (1), 77–88. 10.15252/embj.201592374.

(115) Weaire, D.; Rivier, N. Soap, Cells and Statistics—Random Patterns in Two Dimensions. Contemporary Physics 1984, 25 (1), 59–99.

(116) Weaire, D. L.; Hutzler, S. The Physics of Foams; Oxford University Press, 1999.

(117) Hufnagel, L.; Teleman, A. A.; Rouault, H.; Cohen, S. M.; Shraiman, B. I. On the Mechanism of Wing Size Determination in Fly Development. Proceedings of the National Academy of Sciences 2007, 104 (10), 3835–3840.

(118) Farhadifar, R.; Röper, J.-C.; Aigouy, B.; Eaton, S.; Jülicher, F. The Influence of Cell Mechanics, Cell-Cell Interactions, and Proliferation on Epithelial Packing. Current Biology 2007, 17 (24), 2095–2104. 10.1016/j.cub.2007.11.049.

(119) Gezer, F.; Aykroyd, R. G.; Barber, S. Statistical Properties of Poisson-Voronoi Tessellation Cells in Bounded Regions. Journal of Statistical Computation and Simulation 2021, 91 (5), 915–933. 10.1080/00949655.2020.1836184.

(120) Hinde, A. L.; Miles, R. E. Monte Carlo Estimates of the Distributions of the Random Polygons of the Voronoi Tessellation with Respect to a Poisson Process. Journal of Statistical Computation and Simulation 1980, 10 (3–4), 205–223. 10.1080/00949658008810370.

(121) Tanemura, M. Statistical Distributions of Poisson Voronoi Cells in Two and Three Dimensions. Forma-Tokyo- 2003, 18 (4), 221–247.

(122) Weaire, D.; Kermode, J. P.; Wejchert, J. On the Distribution of Cell Areas in a Voronoi Network. Philosophical Magazine B 1986, 53 (5), L101–L105. 10.1080/13642818608240647.

(123) Dunic, J. C.; Conner, J.; Anderson, S. C.; Thorson, J. T. Introducing the Generalized Gamma Distribution: A Flexible Distribution for Index Standardization. arXiv January 9, 2025. 10.48550/arXiv.2501.05618.

(124) Stacy, E. W. A Generalization of the Gamma Distribution. Ann. Math. Statist. 1962, 33 (3), 1187–1192. 10.1214/aoms/1177704481.

(125) Atia, L.; Bi, D.; Sharma, Y.; Mitchel, J. A.; Gweon, B.; A. Koehler, S.; DeCamp, S. J.; Lan, B.; Kim, J. H.; Hirsch, R.; Pegoraro, A. F.; Lee, K. H.; Starr, J. R.; Weitz, D. A.; Martin, A. C.; Park, J.-A.; Butler, J. P.; Fredberg, J. J. Geometric Constraints during Epithelial Jamming. Nature Phys 2018, 14 (6), 613–620. 10.1038/s41567-018-0089-9.

(126) Du, Q.; Emelianenko, M.; Ju, L. Convergence of the Lloyd Algorithm for Computing Centroidal Voronoi Tessellations. SIAM J. Numer. Anal. 2006, 44 (1), 102–119. 10.1137/040617364.

(127) Du, Q.; Faber, V.; Gunzburger, M. Centroidal Voronoi Tessellations: Applications and Algorithms. SIAM review 1999, 41 (4), 637–676.

(128) Lloyd, S. Least Squares Quantization in PCM. IEEE transactions on information theory 1982, 28 (2), 129–137.

(129) Torquato, S. Reformulation of the Covering and Quantizer Problems as Ground States of Interacting Particles. Phys. Rev. E 2010, 82 (5), 056109. 10.1103/PhysRevE.82.056109.

(130) Klatt, M. A.; Lovrić, J.; Chen, D.; Kapfer, S. C.; Schaller, F. M.; Schönhöfer, P. W.; Gardiner, B. S.; Smith, A.-S.; Schröder-Turk, G. E.; Torquato, S. Universal Hidden Order in Amorphous Cellular Geometries. Nature communications 2019, 10 (1), 811.

(131) Lim, S.; Vermot, J.; Lee, C. F. Vertex Model Mechanics Explain the Emergence of Centroidal Voronoi Tiling in Epithelia. arXiv 2025. 10.48550/ARXIV.2512.13116.

(132) Liu, A. J.; Nagel, S. R. The Jamming Transition and the Marginally Jammed Solid. Annu. Rev. Condens. Matter Phys. 2010, 1 (1), 347–369. 10.1146/annurev-conmatphys-070909-104045.

(133) Needleman, D.; Dogic, Z. Active Matter at the Interface between Materials Science and Cell Biology. Nat Rev Mater 2017, 2 (9), 17048. 10.1038/natrevmats.2017.48.

(134) Kim, S.; Pochitaloff, M.; Stooke-Vaughan, G. A.; Campàs, O. Embryonic Tissues as Active Foams. Nat. Phys. 2021, 17 (7), 859–866. 10.1038/s41567-021-01215-1.

(135) Noll, N.; Mani, M.; Heemskerk, I.; Streichan, S. J.; Shraiman, B. I. Active Tension Network Model Suggests an Exotic Mechanical State Realized in Epithelial Tissues. Nature Phys 2017, 13 (12), 1221–1226. 10.1038/nphys4219.

(136) Prost, J.; Jülicher, F.; Joanny, J.-F. Active Gel Physics. Nature Phys 2015, 11 (2), 111–117. 10.1038/nphys3224.

(137) Manning, M. L.; Foty, R. A.; Steinberg, M. S.; Schoetz, E.-M. Coaction of Intercellular Adhesion and Cortical Tension Specifies Tissue Surface Tension. Proc. Natl. Acad. Sci. U.S.A. 2010, 107 (28), 12517–12522. 10.1073/pnas.1003743107.

(138) Fletcher, A. G.; Osterfield, M.; Baker, R. E.; Shvartsman, S. Y. Vertex Models of Epithelial Morphogenesis. Biophysical Journal 2014, 106 (11), 2291–2304. 10.1016/j.bpj.2013.11.4498.

(139) Röper, K. Supracellular Actomyosin Assemblies: Master Coordinators of Development. Development 2025, 152 (16), dev204896. 10.1242/dev.204896.

(140) Clarke, D. N.; Martin, A. C. Actin-Based Force Generation and Cell Adhesion in Tissue Morphogenesis. Current Biology 2021, 31 (10), R667–R680. 10.1016/j.cub.2021.03.031.

(141) Miao, H.; Blankenship, J. T. The Pulse of Morphogenesis: Actomyosin Dynamics and Regulation in Epithelia. Development 2020, 147 (17), dev186502. 10.1242/dev.186502.

(142) Kim, J. M.; Jo, Y.; Jung, J. W.; Park, K. A Mechanogenetic Role for the Actomyosin Complex in Branching Morphogenesis of Epithelial Organs. Development 2021, 148 (6), dev190785. 10.1242/dev.190785.

(143) Wang, S.; Sekiguchi, R.; Daley, W. P.; Yamada, K. M. Patterned Cell and Matrix Dynamics in Branching Morphogenesis. Journal of Cell Biology 2017, 216 (3), 559–570. 10.1083/jcb.201610048.

(144) Salbreux, G.; Charras, G.; Paluch, E. Actin Cortex Mechanics and Cellular Morphogenesis. Trends in Cell Biology 2012, 22 (10), 536–545. 10.1016/j.tcb.2012.07.001.

(145) Straight, A. F.; Cheung, A.; Limouze, J.; Chen, I.; Westwood, N. J.; Sellers, J. R.; Mitchison, T. J. Dissecting Temporal and Spatial Control of Cytokinesis with a Myosin II Inhibitor. Science 2003, 299 (5613), 1743–1747. 10.1126/science.1081412.

(146) Ishizaki, T.; Uehata, M.; Tamechika, I.; Keel, J.; Nonomura, K.; Maekawa, M.; Narumiya, S. Pharmacological Properties of Y-27632, a Specific Inhibitor of Rho-Associated Kinases. Molecular pharmacology 2000, 57 (5), 976–983.

(147) Saitoh, M.; Ishikawa, T.; Matsushima, S.; Naka, M.; Hidaka, H. Selective Inhibition of Catalytic Activity of Smooth Muscle Myosin Light Chain Kinase. J Biol Chem 1987, 262 (16), 7796–7801.

(148) Surviladze, Z.; Waller, A.; Strouse, J. J.; Bologa, C.; Ursu, O.; Salas, V.; Parkinson, J. F.; Phillips, G. K.; Romero, E.; Wandinger-Ness, A.; Sklar, L. A.; Schroeder, C.; Simpson, D.; Nöth, J.; Wang, J.; Golden, J.; Aubé, J. A Potent and Selective Inhibitor of Cdc42 GTPase. In Probe Reports from the NIH Molecular Libraries Program; National Center for Biotechnology Information (US): Bethesda (MD), 2010.

(149) Quintanilla, M. A.; Hammer, J. A.; Beach, J. R. Non-Muscle Myosin 2 at a Glance. Journal of Cell Science 2023, 136 (5), jcs260890. 10.1242/jcs.260890.

(150) Zaidel-Bar, R.; Zhenhuan, G.; Luxenburg, C. The Contractome – a Systems View of Actomyosin Contractility in Non-Muscle Cells. Journal of Cell Science 2015, 128 (12), 2209–2217. 10.1242/jcs.170068.

(151) Pandya, P.; Orgaz, J. L.; Sanz-Moreno, V. Actomyosin Contractility and Collective Migration: May the Force Be with You. Current Opinion in Cell Biology 2017, 48, 87–96. 10.1016/j.ceb.2017.06.006.

(152) Heer, N. C.; Martin, A. C. Tension, Contraction and Tissue Morphogenesis. Development 2017, 144 (23), 4249–4260. 10.1242/dev.151282.

(153) Campbell, B. C.; Nabel, E. M.; Murdock, M. H.; Lao-Peregrin, C.; Tsoulfas, P.; Blackmore, M. G.; Lee, F. S.; Liston, C.; Morishita, H.; Petsko, G. A. mGreenLantern: A Bright Monomeric Fluorescent Protein with Rapid Expression and Cell Filling Properties for Neuronal Imaging. Proc. Natl. Acad. Sci. U.S.A. 2020, 117 (48), 30710–30721. 10.1073/pnas.2000942117.

(154) Nam, H.; Benezra, R. High Levels of Id1 Expression Define B1 Type Adult Neural Stem Cells. Cell Stem Cell 2009, 5 (5), 515–526. 10.1016/j.stem.2009.08.017.

(155) Ershov, D.; Phan, M.-S.; Pylvänäinen, J. W.; Rigaud, S. U.; Le Blanc, L.; Charles-Orszag, A.; Conway, J. R. W.; Laine, R. F.; Roy, N. H.; Bonazzi, D.; Duménil, G.; Jacquemet, G.; Tinevez, J.-Y. TrackMate 7: Integrating State-of-the-Art Segmentation Algorithms into Tracking Pipelines. Nat Methods 2022, 19 (7), 829–832. 10.1038/s41592-022-01507-1.

(156) Stringer, C.; Pachitariu, M. Cellpose3: One-Click Image Restoration for Improved Cellular Segmentation. Nat Methods 2025, 22 (3), 592–599. 10.1038/s41592-025-02595-5.

(157) Panaretos, V. M.; Zemel, Y. Statistical Aspects of Wasserstein Distances. Annual review of statistics and its application 2019, 6 (1), 405–431.

(158) Yam, P. T.; Wilson, C. A.; Ji, L.; Hebert, B.; Barnhart, E. L.; Dye, N. A.; Wiseman, P. W.; Danuser, G.; Theriot, J. A. Actin–Myosin Network Reorganization Breaks Symmetry at the Cell Rear to Spontaneously Initiate Polarized Cell Motility. The Journal of Cell Biology 2007, 178 (7), 1207–1221. 10.1083/jcb.200706012.

(159) Taneja, N.; Baillargeon, S. M.; Burnette, D. T. Myosin Light Chain Kinase-Driven Myosin II Turnover Regulates Actin Cortex Contractility during Mitosis. MBoC 2021, 32 (20), br3. 10.1091/mbc.E20-09-0608.

(160) Totsukawa, G.; Wu, Y.; Sasaki, Y.; Hartshorne, D. J.; Yamakita, Y.; Yamashiro, S.; Matsumura, F. Distinct Roles of MLCK and ROCK in the Regulation of Membrane Protrusions and Focal Adhesion Dynamics during Cell Migration of Fibroblasts. The Journal of Cell Biology 2004, 164 (3), 427–439. 10.1083/jcb.200306172.

(161) Niggli, V.; Schmid, M.; Nievergelt, A. Differential Roles of Rho-Kinase and Myosin Light Chain Kinase in Regulating Shape, Adhesion, and Migration of HT1080 Fibrosarcoma Cells. Biochemical and Biophysical Research Communications 2006, 343 (2), 602–608. 10.1016/j.bbrc.2006.03.022.

(162) Imai, M.; Furusawa, K.; Mizutani, T.; Kawabata, K.; Haga, H. Three-Dimensional Morphogenesis of MDCK Cells Induced by Cellular Contractile Forces on a Viscous Substrate. Sci Rep 2015, 5 (1), 14208. 10.1038/srep14208.

(163) Totsukawa, G.; Yamakita, Y.; Yamashiro, S.; Hartshorne, D. J.; Sasaki, Y.; Matsumura, F. Distinct Roles of ROCK (Rho-Kinase) and MLCK in Spatial Regulation of MLC Phosphorylation for Assembly of Stress Fibers and Focal Adhesions in 3T3 Fibroblasts. The Journal of cell biology 2000, 150 (4), 797–806.

(164) Haase, K.; Pelling, A. E. The Role of the Actin Cortex in Maintaining Cell Shape. Communicative & Integrative Biology 2013, 6 (6), e26714. 10.4161/cib.26714.

(165) Doss, B. L.; Pan, M.; Gupta, M.; Grenci, G.; Mège, R.-M.; Lim, C. T.; Sheetz, M. P.; Voituriez, R.; Ladoux, B. Cell Response to Substrate Rigidity Is Regulated by Active and Passive Cytoskeletal Stress. Proc. Natl. Acad. Sci. U.S.A. 2020, 117 (23), 12817–12825. 10.1073/pnas.1917555117.

(166) Oakes, P. W.; Beckham, Y.; Stricker, J.; Gardel, M. L. Tension Is Required but Not Sufficient for Focal Adhesion Maturation without a Stress Fiber Template. Journal of Cell Biology 2012, 196 (3), 363–374. 10.1083/jcb.201107042.

(167) Xue, R.; Kang, L.; Chen, Y.; Yang, H.; Jiang, H.; Gong, Z. Force Loading on Molecular Clutches Governs the Stability of Cell Lamellipodia. Proc. Natl. Acad. Sci. U.S.A. 2026, 123 (22), e2604349123. 10.1073/pnas.2604349123.

(168) Wakatsuki, T.; Wysolmerski, R. B.; Elson, E. L. Mechanics of Cell Spreading: Role of Myosin II. Journal of Cell Science 2003, 116 (8), 1617–1625. 10.1242/jcs.00340.

(169) Salhia, B.; Rutten, F.; Nakada, M.; Beaudry, C.; Berens, M.; Kwan, A.; Rutka, J. T. Inhibition of Rho-Kinase Affects Astrocytoma Morphology, Motility, and Invasion through Activation of Rac1. Cancer Research 2005, 65 (19), 8792–8800. 10.1158/0008-5472.CAN-05-0160.

(170) Marshall-Burghardt, S.; Migueles-Ramírez, R. A.; Lin, Q.; El Baba, N.; Saada, R.; Umar, M.; Mavalwala, K.; Hayer, A. Excitable Rho Dynamics Control Cell Shape and Motility by Sequentially Activating ERM Proteins and Actomyosin Contractility. Sci. Adv. 2024, 10 (36), eadn6858. 10.1126/sciadv.adn6858.

(171) Webb, D. J.; Donais, K.; Whitmore, L. A.; Thomas, S. M.; Turner, C. E.; Parsons, J. T.; Horwitz, A. F. FAK–Src Signalling through Paxillin, ERK and MLCK Regulates Adhesion Disassembly. Nature cell biology 2004, 6 (2), 154–161.

(172) Giannone, G.; Dubin-Thaler, B. J.; Döbereiner, H.-G.; Kieffer, N.; Bresnick, A. R.; Sheetz, M. P. Periodic Lamellipodial Contractions Correlate with Rearward Actin Waves. Cell 2004, 116 (3), 431–443. 10.1016/S0092-8674(04)00058-3.

(173) Giannone, G.; Dubin-Thaler, B. J.; Rossier, O.; Cai, Y.; Chaga, O.; Jiang, G.; Beaver, W.; Döbereiner, H.-G.; Freund, Y.; Borisy, G.; Sheetz, M. P. Lamellipodial Actin Mechanically Links Myosin Activity with Adhesion-Site Formation. Cell 2007, 128 (3), 561–575. 10.1016/j.cell.2006.12.039.

(174) Hong, L.; Kenney, S. R.; Phillips, G. K.; Simpson, D.; Schroeder, C. E.; Nöth, J.; Romero, E.; Swanson, S.; Waller, A.; Strouse, J. J.; Carter, M.; Chigaev, A.; Ursu, O.; Oprea, T.; Hjelle, B.; Golden, J. E.; Aubé, J.; Hudson, L. G.; Buranda, T.; Sklar, L. A.; Wandinger-Ness, A. Characterization of a Cdc42 Protein Inhibitor and Its Use as a Molecular Probe. Journal of Biological Chemistry 2013, 288 (12), 8531–8543. 10.1074/jbc.M112.435941.

(175) Kiwanuka, E.; Lee, C. C.; Hackl, F.; Caterson, E. J.; Junker, J. P.; Gerdin, B.; Eriksson, E. Cdc42 and p190RhoGAP Activation by CCN2 Regulates Cell Spreading and Polarity and Induces Actin Disassembly in Migrating Keratinocytes. International Wound Journal 2016, 13 (3), 372–381.

(176) Guan, L.-Y.; Lv, J.-Q.; Zhang, D.-Q.; Li, B. Collective Polarization of Cancer Cells at the Monolayer Boundary. Micromachines 2021, 12 (2), 112. 10.3390/mi12020112.

(177) Yoon, C.; Choi, C.; Stapleton, S.; Mirabella, T.; Howes, C.; Dong, L.; King, J.; Yang, J.; Oberai, A.; Eyckmans, J.; Chen, C. S. Myosin IIA–Mediated Forces Regulate Multicellular Integrity during Vascular Sprouting. MBoC 2019, 30 (16), 1974–1984. 10.1091/mbc.E19-02-0076.

(178) Guan, G.; Cannon, R. D.; Coates, D. E.; Mei, L. Effect of the Rho-Kinase/ROCK Signaling Pathway on Cytoskeleton Components. Genes 2023, 14 (2), 272. 10.3390/genes14020272.

(179) Cantat, I.; Cohen-Addad, S.; Elias, F.; Graner, F.; Höhler, R.; Pitois, O.; Rouyer, F.; Saint-Jalmes, A. Foams: Structure and Dynamics; OUP Oxford, 2013.

(180) Brakke, K. A. Surface Evolver Manual. Mathematics Department, Susquehanna Univerisity, Selinsgrove, PA 1994, 17870 (2.24), 20.

(181) Brakke, K. A. The Surface Evolver. Experimental mathematics 1992, 1 (2), 141–165.

(182) Howard, J.; Grill, S. W.; Bois, J. S. Turing’s next Steps: The Mechanochemical Basis of Morphogenesis. Nat Rev Mol Cell Biol 2011, 12 (6), 392–398. 10.1038/nrm3120.

(183) Varner, V. D.; Nelson, C. M. Cellular and Physical Mechanisms of Branching Morphogenesis. Development 2014, 141 (14), 2750–2759. 10.1242/dev.104794.

(184) Ingber, D. E. Mechanical Signaling and the Cellular Response to Extracellular Matrix in Angiogenesis and Cardiovascular Physiology. Circulation Research 2002, 91 (10), 877–887. 10.1161/01.RES.0000039537.73816.E5.

(185) Mammoto, A.; Connor, K. M.; Mammoto, T.; Yung, C. W.; Huh, D.; Aderman, C. M.; Mostoslavsky, G.; Smith, L. E. H.; Ingber, D. E. A Mechanosensitive Transcriptional Mechanism That Controls Angiogenesis. Nature 2009, 457 (7233), 1103–1108. 10.1038/nature07765.

(186) Kolega, J. Effects of Mechanical Tension on Protrusive Activity and Microfilament and Intermediate Filament Organization in an Epidermal Epithelium Moving in Culture. The Journal of cell biology 1986, 102 (4), 1400–1411. 10.1083/jcb.102.4.1400.

(187) Kourouklis, A. P.; Nelson, C. M. Modeling Branching Morphogenesis Using Materials with Programmable Mechanical Instabilities. Current Opinion in Biomedical Engineering 2018, 6, 66–73. 10.1016/j.cobme.2018.03.007.

(188) Szabo, A.; Mehes, E.; Kosa, E.; Czirok, A. Multicellular Sprouting In Vitro. Biophysical Journal 2008, 95 (6), 2702–2710. 10.1529/biophysj.108.129668.

(189) Debnath, J.; Muthuswamy, S. K.; Brugge, J. S. Morphogenesis and Oncogenesis of MCF-10A Mammary Epithelial Acini Grown in Three-Dimensional Basement Membrane Cultures. Methods 2003, 30 (3), 256–268. 10.1016/S1046-2023(03)00032-X.

(190) Ewald, A. J.; Brenot, A.; Duong, M.; Chan, B. S.; Werb, Z. Collective Epithelial Migration and Cell Rearrangements Drive Mammary Branching Morphogenesis. Developmental Cell 2008, 14 (4), 570–581. 10.1016/j.devcel.2008.03.003.

(191) Lancaster, M. A.; Knoblich, J. A. Generation of Cerebral Organoids from Human Pluripotent Stem Cells. Nat Protoc 2014, 9 (10), 2329–2340. 10.1038/nprot.2014.158.

(192) Azizoglu, D. B.; Braitsch, C.; Marciano, D. K.; Cleaver, O. Afadin and RhoA Control Pancreatic Endocrine Mass via Lumen Morphogenesis. Genes & development 2017, 31 (23–24), 2376–2390.

(193) Debnath, J.; Mills, K. R.; Collins, N. L.; Reginato, M. J.; Muthuswamy, S. K.; Brugge, J. S. The Role of Apoptosis in Creating and Maintaining Luminal Space within Normal and Oncogene-Expressing Mammary Acini. Cell 2002, 111 (1), 29–40.

(194) Lubarsky, B.; Krasnow, M. A. Tube Morphogenesis: Making and Shaping Biological Tubes. Cell 2003, 112 (1), 19–28.

(195) Kaisani, A.; Delgado, O.; Fasciani, G.; Kim, S. B.; Wright, W. E.; Minna, J. D.; Shay, J. W. Branching Morphogenesis of Immortalized Human Bronchial Epithelial Cells in Three-Dimensional Culture. Differentiation 2014, 87 (3–4), 119–126. 10.1016/j.diff.2014.02.003.

(196) Montesano, R.; Ghzili, H.; Carrozzino, F.; Rossier, B. C.; Féraille, E. cAMP-Dependent Chloride Secretion Mediates Tubule Enlargement and Cyst Formation by Cultured Mammalian Collecting Duct Cells. American Journal of Physiology-Renal Physiology 2009, 296 (2), F446–F457.

(197) Montesano, R.; Soulié, P. Retinoids Induce Lumen Morphogenesis in Mammary Epithelial Cells. Journal of cell science 2002, 115 (23), 4419–4431.

(198) Jamieson, P. R.; Dekkers, J. F.; Rios, A. C.; Fu, N. Y.; Lindeman, G. J.; Visvader, J. E. Derivation of a Robust Mouse Mammary Organoid System for Studying Tissue Dynamics. Development 2017, 144 (6), 1065–1071.

(199) Lee, B. H.; Fuji, K.; Petzold, H.; Seymour, P.; Yennek, S.; Schewin, C.; Lewis, A.; Riveline, D.; Hiraiwa, T.; Sano, M.; Grapin-Botton, A. Permeability-Driven Pressure and Cell Proliferation Control Lumen Morphogenesis in Pancreatic Organoids. Nat Cell Biol 2026, 28 (1), 113–124. 10.1038/s41556-025-01832-5.

(200) Mae, S.-I.; Ryosaka, M.; Sakamoto, S.; Matsuse, K.; Nozaki, A.; Igami, M.; Kabai, R.; Watanabe, A.; Osafune, K. Expansion of Human iPSC-Derived Ureteric Bud Organoids with Repeated Branching Potential. Cell Reports 2020, 32 (4).

(201) Papargyriou, A.; Najajreh, M.; Cook, D. P.; Maurer, C. H.; Bärthel, S.; Messal, H. A.; Ravichandran, S. K.; Richter, T.; Knolle, M.; Metzler, T. Heterogeneity-Driven Phenotypic Plasticity and Treatment Response in Branched-Organoid Models of Pancreatic Ductal Adenocarcinoma. Nature biomedical engineering 2025, 9 (6), 836–864.

(202) Randriamanantsoa, S.; Papargyriou, A.; Maurer, H. C.; Peschke, K.; Schuster, M.; Zecchin, G.; Steiger, K.; Öllinger, R.; Saur, D.; Scheel, C.; Rad, R.; Hannezo, E.; Reichert, M.; Bausch, A. R. Spatiotemporal Dynamics of Self-Organized Branching in Pancreas-Derived Organoids. Nat Commun 2022, 13 (1), 5219. 10.1038/s41467-022-32806-y.

(203) Roos, F. J.; van Tienderen, G. S.; Wu, H.; Bordeu, I.; Vinke, D.; Albarinos, L. M.; Monfils, K.; Niesten, S.; Smits, R.; Willemse, J. Human Branching Cholangiocyte Organoids Recapitulate Functional Bile Duct Formation. Cell Stem Cell 2022, 29 (5), 776–794.

(204) Lu, L.; Fuji, K.; Guyomar, T.; Lieb, M.; André, M.; Tanida, S.; Nonomura, M.; Hiraiwa, T.; Alcheikh, Y.; Yennek, S.; Petzold, H.; Martin-Lemaitre, C.; Grapin-Botton, A.; Honigmann, A.; Sano, M.; Riveline, D. Generic Comparison of Lumen Nucleation and Fusion in Epithelial Organoids with and without Hydrostatic Pressure. Nat Commun 2025, 16 (1), 6307. 10.1038/s41467-025-60780-8.

(205) Yang, M.; Du, Z. To See and to Know: The Power of Live Imaging in Illuminating and Decoding Biological Complexity. Journal of Genetics and Genomics 2025, S1673852725002826. 10.1016/j.jgg.2025.10.003.

(206) Araújo, S. J.; Llimargas, M. Time-Lapse Imaging and Morphometric Analysis of Tracheal Development in Drosophila. In Cell Migration in Three Dimensions; Springer, 2023; pp 163–182.

(207) Löf-Öhlin, Z. M.; Nyeng, P.; Bechard, M. E.; Hess, K.; Bankaitis, E.; Greiner, T. U.; Ameri, J.; Wright, C. V.; Semb, H. EGFR Signalling Controls Cellular Fate and Pancreatic Organogenesis by Regulating Apicobasal Polarity. Nature cell biology 2017, 19 (11), 1313–1325.

(208) Mullapudi, S. T.; Boezio, G. L.; Rossi, A.; Marass, M.; Matsuoka, R. L.; Matsuda, H.; Helker, C. S.; Yang, Y. H. C.; Stainier, D. Y. Disruption of the Pancreatic Vasculature in Zebrafish Affects Islet Architecture and Function. Development 2019, 146 (21), dev173674.

(209) Packard, A.; Georgas, K.; Michos, O.; Riccio, P.; Cebrian, C.; Combes, A. N.; Ju, A.; Ferrer-Vaquer, A.; Hadjantonakis, A.-K.; Zong, H. Luminal Mitosis Drives Epithelial Cell Dispersal within the Branching Ureteric Bud. Developmental cell 2013, 27 (3), 319–330.

(210) Riccio, P.; Cebrian, C.; Zong, H.; Hippenmeyer, S.; Costantini, F. Ret and Etv4 Promote Directed Movements of Progenitor Cells during Renal Branching Morphogenesis. PLoS biology 2016, 14 (2), e1002382.

(211) Scheele, C. L.; Hannezo, E.; Muraro, M. J.; Zomer, A.; Langedijk, N. S.; Van Oudenaarden, A.; Simons, B. D.; Van Rheenen, J. Identity and Dynamics of Mammary Stem Cells during Branching Morphogenesis. Nature 2017, 542 (7641), 313–317.

(212) Shih, H. P.; Panlasigui, D.; Cirulli, V.; Sander, M. ECM Signaling Regulates Collective Cellular Dynamics to Control Pancreas Branching Morphogenesis. Cell reports 2016, 14 (2), 169–179.

(213) Balasubramani, V.; Kujawińska, M.; Allier, C.; Anand, V.; Cheng, C.-J.; Depeursinge, C.; Hai, N.; Juodkazis, S.; Kalkman, J.; Kuś, A.; Lee, M.; Magistretti, P. J.; Marquet, P.; Ng, S. H.; Rosen, J.; Park, Y. K.; Ziemczonok, M. Roadmap on Digital Holography-Based Quantitative Phase Imaging. J. Imaging 2021, 7 (12), 252. 10.3390/jimaging7120252.

(214) Moore, S. P.; Zou, A.; Zhang, X.; Jonathan, O. C.; Lang, D.; Zhang, C. VaMiAnalyzer: An Open Source, Python-Based Application for Analysis of 3D in Vitro Vasculogenic Mimicry Assays. BMC bioinformatics 2025, 26 (1), 1–11.

(215) Schüttler, M.; Doğan, L.; Kirchner, J.; Ergün, S.; Wörsdörfer, P.; Fischer, S. C. VESNA: An Open-Source Tool for Automated 3D Vessel Segmentation and Network Analysis. Bioinformatics March 10, 2025. 10.1101/2025.03.05.641600.

(216) Alves, A. P.; Mesquita, O. N.; Gómez-Gardeñes, J.; Agero, U. Graph Analysis of Cell Clusters Forming Vascular Networks. R. Soc. open sci. 2018, 5 (3), 171592. 10.1098/rsos.171592.

(217) Vicente-Munuera, P.; Gómez-Gálvez, P.; Tetley, R. J.; Forja, C.; Tagua, A.; Letrán, M.; Tozluoglu, M.; Mao, Y.; Escudero, L. M. EpiGraph: An Open-Source Platform to Quantify Epithelial Organization. Bioinformatics 2020, 36 (4), 1314–1316. 10.1093/bioinformatics/btz683.

(218) Bordeu, I.; Chatzeli, L.; Simons, B. D. Inflationary Theory of Branching Morphogenesis in the Mouse Salivary Gland. Nat Commun 2023, 14 (1), 3422. 10.1038/s41467-023-39124-x.

(219) Brown, A. I.; Westrate, L. M.; Koslover, E. F. Impact of Global Structure on Diffusive Exploration of Organelle Networks. Sci Rep 2020, 10 (1), 4984. 10.1038/s41598-020-61598-8.

(220) Lucas, M.; Bisot, C.; Petri, G.; Declerck, S.; Carletti, T. Minimal Branching and Fusion Morphogenesis Approaches Biological Multi-Objective Optimality. arXiv 2026. 10.48550/ARXIV.2601.03877.

(221) Stauffer, D. Scaling Theory of Percolation Clusters. Physics Reports 1979, 54 (1), 1–74. 10.1016/0370-1573(79)90060-7.

(222) Stauffer, D.; Aharony, A. Introduction to Percolation Theory; Taylor & Francis, 2018.

(223) Aon, M. A.; Cortassa, S.; O’Rourke, B. Percolation and Criticality in a Mitochondrial Network. Proc. Natl. Acad. Sci. U.S.A. 2004, 101 (13), 4447–4452. 10.1073/pnas.0307156101.

(224) Atia, L.; Fredberg, J. J.; Gov, N. S.; Pegoraro, A. F. Are Cell Jamming and Unjamming Essential in Tissue Development? Cells & Development 2021, 168, 203727. 10.1016/j.cdev.2021.203727.

(225) Cardy, J.; Täuber, U. C. Theory of Branching and Annihilating Random Walks. Phys. Rev. Lett. 1996, 77 (23), 4780–4783. 10.1103/PhysRevLett.77.4780.

(226) Fessel, A.; Oettmeier, C.; Bernitt, E.; Gauthier, N. C.; Döbereiner, H.-G. Physarum Polycephalum Percolation as a Paradigm for Topological Phase Transitions in Transportation Networks. Phys. Rev. Lett. 2012, 109 (7), 078103. 10.1103/PhysRevLett.109.078103.

(227) Kirkegaard, J. B.; Nielsen, B. F.; Trusina, A.; Sneppen, K. Self-Assembly, Buckling and Density-Invariant Growth of Three-Dimensional Vascular Networks. J. R. Soc. Interface. 2019, 16 (159), 20190517. 10.1098/rsif.2019.0517.

(228) Kudryashova, N.; Nizamieva, A.; Tsvelaya, V.; Panfilov, A. V.; Agladze, K. I. Self-Organization of Conducting Pathways Explains Electrical Wave Propagation in Cardiac Tissues with High Fraction of Non-Conducting Cells. PLoS Comput Biol 2019, 15 (3), e1006597. 10.1371/journal.pcbi.1006597.

(229) Noerr, P. S.; Zamora Alvarado, J. E.; Golnaraghi, F.; McCloskey, K. E.; Gopinathan, A.; Dasbiswas, K. Optimal Mechanical Interactions Direct Multicellular Network Formation on Elastic Substrates. Proc. Natl. Acad. Sci. U.S.A. 2023, 120 (45), e2301555120. 10.1073/pnas.2301555120.

(230) Petridou, N. I.; Corominas-Murtra, B.; Heisenberg, C.-P.; Hannezo, E. Rigidity Percolation Uncovers a Structural Basis for Embryonic Tissue Phase Transitions. Cell 2021, 184 (7), 1914–1928.e19. 10.1016/j.cell.2021.02.017.

(231) Szabo, A.; Perryn, E. D.; Czirok, A. Network Formation of Tissue Cells via Preferential Attraction to Elongated Structures. Phys. Rev. Lett. 2007, 98 (3), 038102. 10.1103/PhysRevLett.98.038102.

(232) Alt, S.; Ganguly, P.; Salbreux, G. Vertex Models: From Cell Mechanics to Tissue Morphogenesis. Phil. Trans. R. Soc. B 2017, 372 (1720), 20150520. 10.1098/rstb.2015.0520.

(233) Jaynes, E. T. The Maximum Entropy Formalism, edited by R. D. Levine and M. Tribus.; MIT Press, 1979.

(234) Ohlenbusch, H.; Aste, T.; Dubertret, B.; Rivier, N. The Topological Structure of 2D Disordered Cellular Systems. The European Physical Journal B-Condensed Matter and Complex Systems 1998, 2 (2), 211–220.

(235) Baule, A.; Morone, F.; Herrmann, H. J.; Makse, H. A. Edwards Statistical Mechanics for Jammed Granular Matter. Rev. Mod. Phys. 2018, 90 (1), 015006. 10.1103/RevModPhys.90.015006.

(236) Edwards, S. F.; Oakeshott, R. Theory of Powders. Physica A: Statistical Mechanics and its Applications 1989, 157 (3), 1080–1090.

(237) Choksi, R.; Lu, X. Y. Bounds on the Geometric Complexity of Optimal Centroidal Voronoi Tesselations in 3D. Commun. Math. Phys. 2020, 377 (3), 2429–2450. 10.1007/s00220-020-03789-y.

(238) Du, Q.; Gunzburger, M.; Ju, L. Advances in Studies and Applications of Centroidal Voronoi Tessellations. Numerical Mathematics: Theory, Methods and Applications 2010, 3 (2), 119–142.

(239) Spatial Tessellations: Concepts and Applications of Voronoi Diagrams, 2nd ed.; Okabe, A., Boots, B., Sugihara, K., Chiu, S. N., Eds.; Wiley series in probability and statistics; Wiley: Chichester, 2000.

(240) Newman, D. The Hexagon Theorem. IEEE Trans. Inform. Theory 1982, 28 (2), 137–139. 10.1109/TIT.1982.1056492.

(241) Lawson-Keister, E.; Manning, M. L. Jamming and Arrest of Cell Motion in Biological Tissues. Current Opinion in Cell Biology 2021, 72, 146–155. 10.1016/j.ceb.2021.07.011.

(242) Mao, Y.; Wickström, S. A. Mechanical State Transitions in the Regulation of Tissue Form and Function. Nat Rev Mol Cell Biol 2024, 25 (8), 654–670. 10.1038/s41580-024-00719-x.

(243) Tetley, R. J.; Staddon, M. F.; Heller, D.; Hoppe, A.; Banerjee, S.; Mao, Y. Tissue Fluidity Promotes Epithelial Wound Healing. Nat. Phys. 2019, 15 (11), 1195–1203. 10.1038/s41567-019-0618-1.

(244) Grandy, C.; Port, F.; Pfeil, J.; Gottschalk, K.-E. Influence of ROCK Pathway Manipulation on the Actin Cytoskeleton Height. Cells 2022, 11 (3), 430. 10.3390/cells11030430.

(245) Amano, M.; Nakayama, M.; Kaibuchi, K. Rho-kinase/ROCK: A Key Regulator of the Cytoskeleton and Cell Polarity. Cytoskeleton 2010, 67 (9), 545–554. 10.1002/cm.20472.

(246) Berndt, J. D.; Clay, M. R.; Langenberg, T.; Halloran, M. C. Rho-Kinase and Myosin II Affect Dynamic Neural Crest Cell Behaviors during Epithelial to Mesenchymal Transition in Vivo. Developmental Biology 2008, 324 (2), 236–244. 10.1016/j.ydbio.2008.09.013.

(247) Horváth, Á. I.; Gyimesi, M.; Várkuti, B. H.; Képiró, M.; Szegvári, G.; Lőrincz, I.; Hegyi, G.; Kovács, M.; Málnási-Csizmadia, A. Effect of Allosteric Inhibition of Non-Muscle Myosin 2 on Its Intracellular Diffusion. Sci Rep 2020, 10 (1), 13341. 10.1038/s41598-020-69853-8.

(248) Martens, J. C.; Radmacher, M. Softening of the Actin Cytoskeleton by Inhibition of Myosin II. Pflugers Arch - Eur J Physiol 2008, 456 (1), 95–100. 10.1007/s00424-007-0419-8.

(249) Rosenfeld, D.; Landau, S.; Shandalov, Y.; Raindel, N.; Freiman, A.; Shor, E.; Blinder, Y.; Vandenburgh, H. H.; Mooney, D. J.; Levenberg, S. Morphogenesis of 3D Vascular Networks Is Regulated by Tensile Forces. Proc. Natl. Acad. Sci. U.S.A. 2016, 113 (12), 3215–3220. 10.1073/pnas.1522273113.

(250) Ubukawa, K.; Guo, Y.-M.; Takahashi, M.; Hirokawa, M.; Michishita, Y.; Nara, M.; Tagawa, H.; Takahashi, N.; Komatsuda, A.; Nunomura, W.; Takakuwa, Y.; Sawada, K. Enucleation of Human Erythroblasts Involves Non-Muscle Myosin IIB. Blood 2012, 119 (4), 1036–1044. 10.1182/blood-2011-06-361907.

(251) Fischer, R. S.; Gardel, M.; Ma, X.; Adelstein, R. S.; Waterman, C. M. Local Cortical Tension by Myosin II Guides 3D Endothelial Cell Branching. Current Biology 2009, 19 (3), 260–265. 10.1016/j.cub.2008.12.045.

(252) Kroll, J.; Epting, D.; Kern, K.; Dietz, C. T.; Feng, Y.; Hammes, H.-P.; Wieland, T.; Augustin, H. G. Inhibition of Rho-Dependent Kinases ROCK I/II Activates VEGF-Driven Retinal Neovascularization and Sprouting Angiogenesis. American Journal of Physiology-Heart and Circulatory Physiology 2009, 296 (3), H893–H899. 10.1152/ajpheart.01038.2008.

(253) Wang, S.; Sun, J.; Xiao, Y.; Lu, Y.; Zhang, D. D.; Wong, P. K. Intercellular Tension Negatively Regulates Angiogenic Sprouting of Endothelial Tip Cells via Notch1-Dll4 Signaling. Adv. Biosys. 2017, 1 (1–2), 1600019. 10.1002/adbi.201600019.

(254) Ehrig, S.; Schamberger, B.; Bidan, C. M.; West, A.; Jacobi, C.; Lam, K.; Kollmannsberger, P.; Petersen, A.; Tomancak, P.; Kommareddy, K.; Fischer, F. D.; Fratzl, P.; Dunlop, J. W. C. Surface Tension Determines Tissue Shape and Growth Kinetics. Sci. Adv. 2019, 5 (9), eaav9394. 10.1126/sciadv.aav9394.

(255) Hilgenfeldt, S.; Erisken, S.; Carthew, R. W. Physical Modeling of Cell Geometric Order in an Epithelial Tissue. Proc. Natl. Acad. Sci. U.S.A. 2008, 105 (3), 907–911. 10.1073/pnas.0711077105.

(256) Lecuit, T.; Lenne, P.-F. Cell Surface Mechanics and the Control of Cell Shape, Tissue Patterns and Morphogenesis. Nat Rev Mol Cell Biol 2007, 8 (8), 633–644. 10.1038/nrm2222.

(257) Winklbauer, R. Cell Adhesion Strength from Cortical Tension – an Integration of Concepts. Journal of Cell Science 2015, 128 (20), 3687–3693. 10.1242/jcs.174623.

(258) Pérez-González, C.; Alert, R.; Blanch-Mercader, C.; Gómez-González, M.; Kolodziej, T.; Bazellieres, E.; Casademunt, J.; Trepat, X. Active Wetting of Epithelial Tissues. Nature Phys 2019, 15 (1), 79–88. 10.1038/s41567-018-0279-5.

(259) Drenckhan, W.; Hutzler, S. Structure and Energy of Liquid Foams. Advances in colloid and interface science 2015, 224, 1–16.

(260) Graner, F.; Glazier, J. A. Simulation of Biological Cell Sorting Using a Two-Dimensional Extended Potts Model. Physical review letters 1992, 69 (13), 2013.

(261) Popović, M.; Druelle, V.; Dye, N. A.; Jülicher, F.; Wyart, M. Inferring the Flow Properties of Epithelial Tissues from Their Geometry. New J. Phys. 2021, 23 (3), 033004. 10.1088/1367-2630/abcbc7.

(262) Bohn, S.; Douady, S.; Couder, Y. Four Sided Domains in Hierarchical Space Dividing Patterns. Phys. Rev. Lett. 2005, 94 (5), 054503. 10.1103/PhysRevLett.94.054503.

(263) Matos, I. S.; Vu, B.; Mann, J.; Xie, E.; Madhavan, S.; Sharma, S.; Niewiadomski, I.; Echevarria, A.; Tomaka, C.; Carlos, S.; Antonio, M.; Chu, A.; Scudder, M.; Yokota, N.; Park, H. J.; Vuong, N.; Boakye, M.; Duarte, M. A.; Pechuzal, C.; Aparecido, L. M. T.; Franco, M. B.; Wong, R. J.; Liu, J.; Guevara Heredia, E.; Boyle, B.; Ryan, M.; Cárdenas, R. E.; Enquist, B. J.; Erwin, D. M.; Forbes, H.; Dexter, K.; Fricker, M.; Blonder, B. W. Leaf Venation Network Evolution across Clades and Scales. Nat. Plants 2025, 11 (6), 1127–1141. 10.1038/s41477-025-02011-y.

(264) Mittler, F.; Obeïd, P.; Rulina, A. V.; Haguet, V.; Gidrol, X.; Balakirev, M. Y. High-Content Monitoring of Drug Effects in a 3D Spheroid Model. Frontiers in Oncology 2017, 7. 10.3389/fonc.2017.00293.

(265) Pachitariu, M.; Rariden, M.; Stringer, C. Cellpose-SAM: Superhuman Generalization for Cellular Segmentation. Bioinformatics May 1, 2025. 10.1101/2025.04.28.651001.

(266) Stringer, C.; Wang, T.; Michaelos, M.; Pachitariu, M. Cellpose: A Generalist Algorithm for Cellular Segmentation. Nature Methods 2021, 18 (1), 100–106. 10.1038/s41592-020-01018-x.

(267) Kesavan, S. V.; Momey, F.; Cioni, O.; David-Watine, B.; Dubrulle, N.; Shorte, S.; Sulpice, E.; Freida, D.; Chalmond, B. High-Throughput Monitoring of Major Cell Functions by Means of Lensfree Video Microscopy. Scientific reports 2014, 4 (1), 5942.

(268) Martin, A.; Godefroy, G.; Lemarchand, F.; Neri, J.; Cioni, O.; Padmanabhan, K.; Paviolo, C. He2Cl: A 2-Step Clustering Algorithm to Characterize Cellular Heterogeneity From Cell Morpho-Dynamics Behaviors. IEEE Trans. Signal Process. 2025, 73, 3394–3405. 10.1109/TSP.2025.3590564.

(269) Allier, C.; Hervé, L.; Paviolo, C.; Mandula, O.; Cioni, O.; Pierré, W.; Andriani, F.; Padmanabhan, K.; Morales, S. CNN-Based Cell Analysis: From Image to Quantitative Representation. Front. Phys. 2022, 9, 776805. 10.3389/fphy.2021.776805.

(270) Hervé, L.; Kraemer, D. C. A.; Cioni, O.; Mandula, O.; Menneteau, M.; Morales, S.; Allier, C. Alternation of Inverse Problem Approach and Deep Learning for Lens-Free Microscopy Image Reconstruction. Sci Rep 2020, 10 (1), 20207. 10.1038/s41598-020-76411-9.

(271) Otsu, N. A Threshold Selection Method from Gray-Level Histograms. IEEE Trans. Syst., Man, Cybern. 1979, 9 (1), 62–66. 10.1109/TSMC.1979.4310076.

(272) Bradley, D.; Roth, G. Adaptive Thresholding Using the Integral Image. Journal of Graphics Tools 2007, 12 (2), 13–21. 10.1080/2151237X.2007.10129236.

(273) Soille, P. Morphological Image Analysis; Springer Berlin Heidelberg: Berlin, Heidelberg, 2004. 10.1007/978-3-662-05088-0.

(274) Zuiderveld, K. Contrast Limited Adaptive Histogram Equalization. In Graphics Gems; Elsevier, 1994; pp 474–485. 10.1016/B978-0-12-336156-1.50061-6.

(275) Zhang, T. Y.; Suen, C. Y. A Fast Parallel Algorithm for Thinning Digital Patterns. Commun. ACM 1984, 27 (3), 236–239. 10.1145/357994.358023.

(276) Boeing, G. Topological Graph Simplification Solutions to the Street Intersection Miscount Problem. 2024. 10.48550/ARXIV.2407.00258.

(277) Viana, M. P.; Strano, E.; Bordin, P.; Barthelemy, M. The Simplicity of Planar Networks. Sci Rep 2013, 3 (1), 3495. 10.1038/srep03495.

(278) Newman, M. E. J. The Structure and Function of Complex Networks. SIAM Rev. 2003, 45 (2), 167–256. 10.1137/S003614450342480.

(279) Albert, R. Scale-Free Networks in Cell Biology. Journal of Cell Science 2005, 118 (21), 4947–4957. 10.1242/jcs.02714.

(280) Strogatz, S. H. Exploring Complex Networks. Nature 2001, 410 (6825), 268–276. 10.1038/35065725.

(281) Fischer, S. C.; Bassel, G. W.; Kollmannsberger, P. Tissues as Networks of Cells: Towards Generative Rules of Complex Organ Development. J. R. Soc. Interface. 2023, 20 (204), 20230115. 10.1098/rsif.2023.0115.

(282) Latora, V.; Marchiori, M. Efficient Behavior of Small-World Networks. Phys. Rev. Lett. 2001, 87 (19), 198701. 10.1103/PhysRevLett.87.198701.

(283) Buhl, C.; Gautrais, J.; Solé, R. V.; Kuntz, P.; Valverde, S.; Deneubourg, J. L.; Theraulaz, G. Efficiency and Robustness in Ant Networks Ofgalleries. Eur. Phys. J. B 2004, 42 (1), 123–129. 10.1140/epjb/e2004-00364-9.

(284) Edelsbrunner; Letscher; Zomorodian. Topological Persistence and Simplification. Discrete Comput Geom 2002, 28 (4), 511–533. 10.1007/s00454-002-2885-2.

(285) Ghrist, R. Barcodes: The Persistent Topology of Data. Bull. Amer. Math. Soc. 2007, 45 (1), 61–75. 10.1090/S0273-0979-07-01191-3.

(286) Torquato, S. Hyperuniform States of Matter. Physics Reports 2018, 745, 1–95. 10.1016/j.physrep.2018.03.001.

