## Supplementary Materials for "Space partitioning by self-organized epithelial networks"

*Short title: Epithelial Networks Partition Space*

Ronny Tonato Zambrano[1]*, Ayla Biallas[2], Frédérique Mittler[2], Alice Nicolas[3], Xavier Gidrol[2], Sophie Achard[4], Lionel Hervé[1], Guillaume Godefroy[1]*, Maxim Y. Balakirev[2]*

[1]*Univ. Grenoble Alpes, CEA-Leti, F-38000 Grenoble, France*
[2]*Univ. Grenoble Alpes, CEA, INSERM, IRIG, Large-Scale Biology, Biomics, F-38054 Grenoble, France*
[3]*Univ. Grenoble Alpes, CNRS, CEA-Leti-Minatec, Grenoble INP, F-38000 LTM, Grenoble, France*
[4]*Univ. Grenoble Alpes, CNRS, Inria, Grenoble INP, LJK, F-38000 Grenoble, France*

**This PDF file includes:**

Supplementary Table S1……………………………………………………………..page 2

Supplementary Figures S1 to S6 (with legends)………………………………………. page 3 - 8

Representative screenshots for Supplementary Videos S1 to S8 (with legends)………… page 9 - 16

**Other Supplementary Materials for this manuscript include the following:**

Supplementary Videos S1 to S8

| Name | Description | Lineage | Type | Network |
|---|---|---|---|---|
| ADSC-d2 | Adipose-derived stem cells | Stem cells | Primary | N |
| ADSC-d6 | Adipose-derived stem cells | Stem cells | Primary | N |
| IPSC | Induced pluripotent stem cells | Stem cells | Primary | N |
| MCS | Mesenchymal stem cells | Stem cells | Primary | N |
| CAF | Cancer-associated fibroblasts | Fibroblasts | Primary | N |
| hDF | Dermal fibroblast | Fibroblasts | Primary | N |
| hLF | Lung fibroblasts | Fibroblasts | Primary | N |
| HPASTEC | Pancreatic stellate cells | Fibroblasts | Primary | N |
| HPNE | Immortalized pancreatic nestin-expressing cells | Fibroblastoid | Cell line | N |
| HPDE | Immortalized pancreatic ductal epithelial cells H6c7 | Epithelial | Cell line | Y |
| HPPE | Primary pancreatic epithelial cells | Epithelial | Primary | Y/N |
| HEK | Immortalized embryonic kidney cells | Epithelial | Cell line | Y |
| MCF10a | Immortalized breast epithelial cells | Epithelial | Cell line | Y |
| RPE1 | Immortalized retinal pigment epithelial cells | Epithelial | Cell line | N |
| RWPE1 | Immortalized prostate epithelial cells | Epithelial | Cell line | Y |
| HaCaT | Immortalized keratinocytes | Keratinocytes | Cell line | N |
| HKPM | Primary skin keratinocytes | Keratinocytes | Primary | N |
| hTert-Kera | Immortalized keratinocytes | Keratinocytes | Cell line | N |
| HPaMVEC | Pancreatic microvascular endothelial cells | Endothelial | Primary | Y/N |
| HUVEC | Umbilical vein endothelial cells | Endothelial | Primary | Y |

**Supplementary Table S1 | Ability of human non-cancerous cell types to form supracellular networks in vitro.** (N), no network formation observed under the experimental conditions tested; (Y), stable supracellular network formation observed; (Y/N), partial or transient network formation observed.

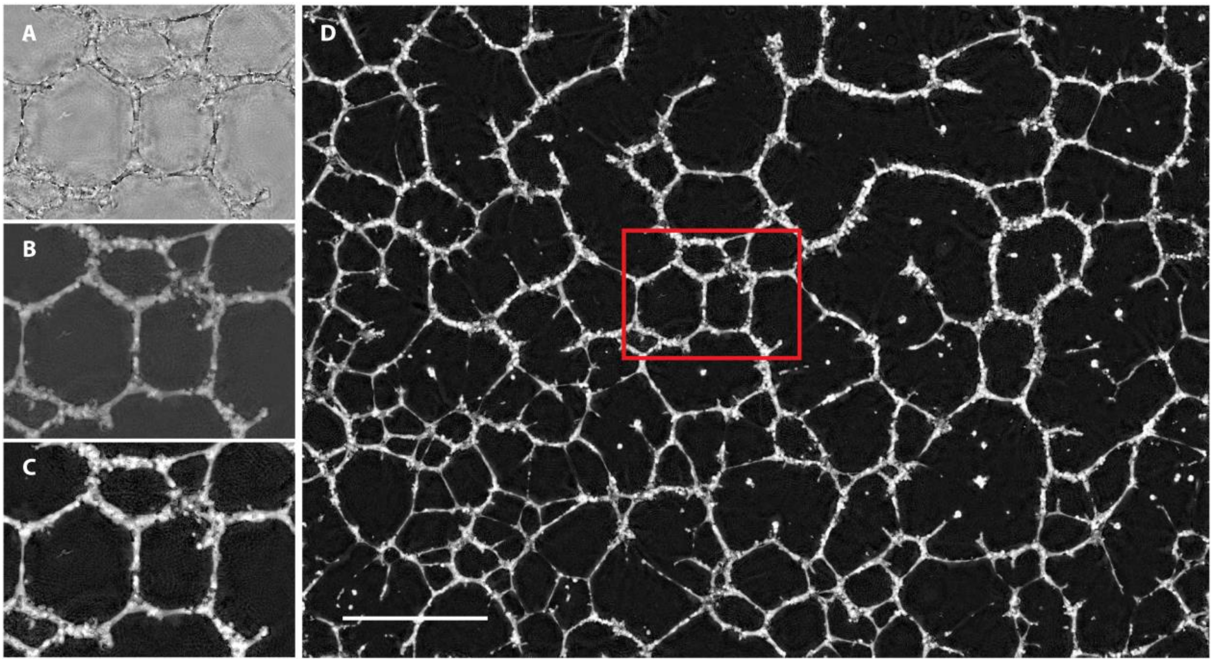


**Supplementary Figure S1 | Holographic reconstruction pipeline.** (**A**) Optical path difference (OPD) map obtained after solving the first inverse problem. (**B**) OPD map after inference using the phase-unwrapping convolutional neural network (CNN). (**C**) Final OPD map obtained after solving the second inverse problem. (**D**) Full-field reconstruction of the final OPD map. Scale bar, 1mm.

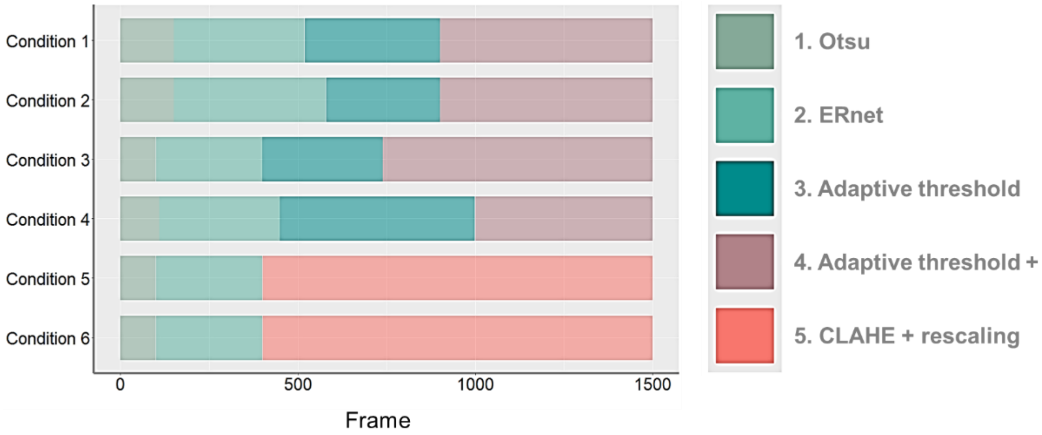


**Supplementary Figure S2 | Implementation of segmentation strategies.** Six phase-reconstructed time-lapse videos (~1500 frames per condition; conditions 1–6) were analyzed using five segmentation strategies.

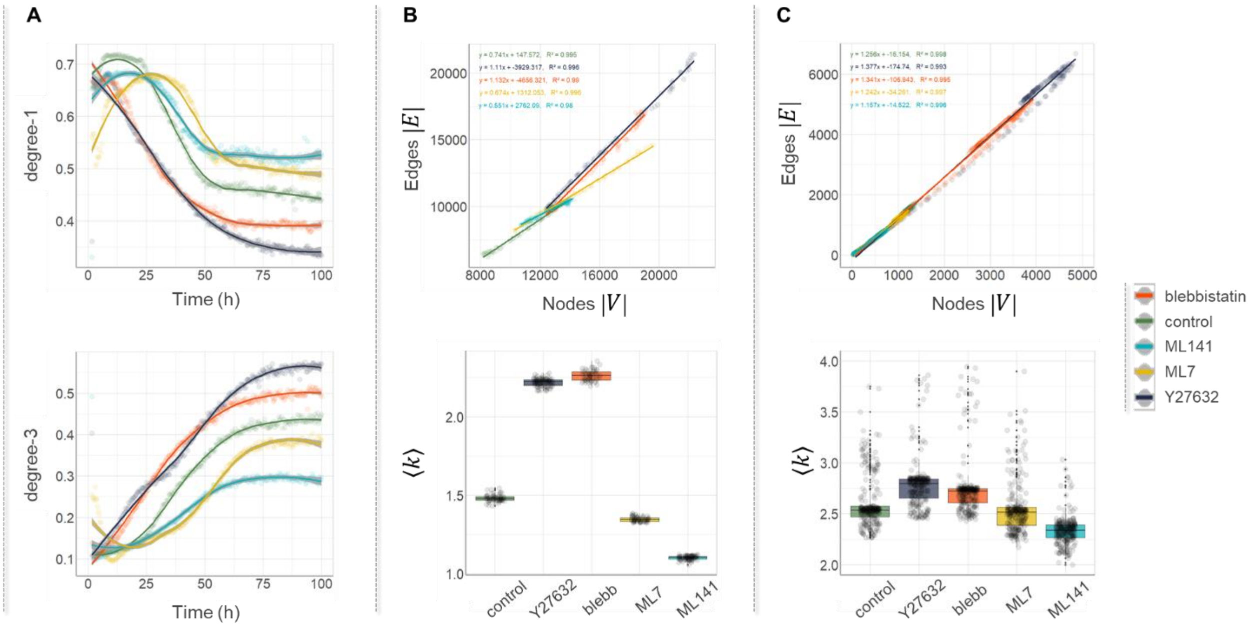


**Supplementary Figure S3 | Graph node degree analysis.** (**A**) Temporal evolution of the fractions of degree-1 and degree-3 graph nodes following pharmacological perturbations. Data points are color-coded by experimental condition; the same color scheme is used throughout all panels and is indicated on the right side of the figure. (**B**) Top panel: Number of edges ($|E|$) plotted as a function of the number of nodes ($|V|$) for complete EN graphs under different experimental conditions. Data are fitted by linear regressions, with the corresponding equations and $R^2$ values shown. Bottom panel: Boxplots of the mean node degree, $\langle k \rangle$, defined as the average number of edges incident on a node and calculated as $\langle k \rangle = 2|E|/|V|$. The mean node degree is directly related to the slope of the linear regressions shown in the top panel. (**C**) Same analysis as in **B**, but performed on pruned, angle-filtered simplified graphs (see Materials and Methods).

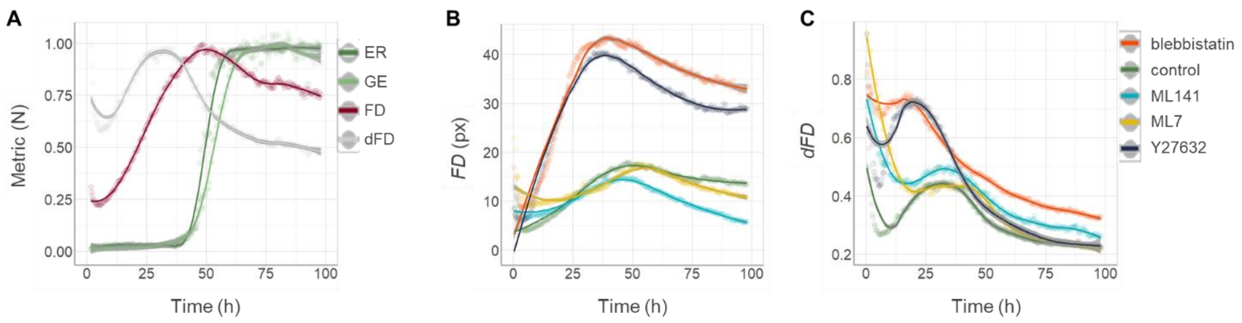


**Supplementary Figure S4 | Epithelial network dynamics.** (**A**) Time-resolved metrics of EN dynamics, normalized to their respective maximum values. Network dynamics were quantified using the frame-difference ($FD$) metric, calculated as the pixel-wise difference between consecutive binarized image frames using the Fiji Kymograph package. The normalized frame-difference metric ($dFD$) was obtained by dividing $FD$ by its corresponding moving average, as implemented in the same package. These metrics are compared with the corresponding graph edge ratio ($ER$) and global efficiency ($GE$) metrics. (**B**) Temporal evolution of $FD$ following pharmacological perturbations. Data points are color-coded according to experimental condition; the same color scheme is used in panel **C** and is indicated on the right side of the figure. (**C**) Temporal evolution of the normalized frame-difference metric ($dFD$) following pharmacological perturbations.

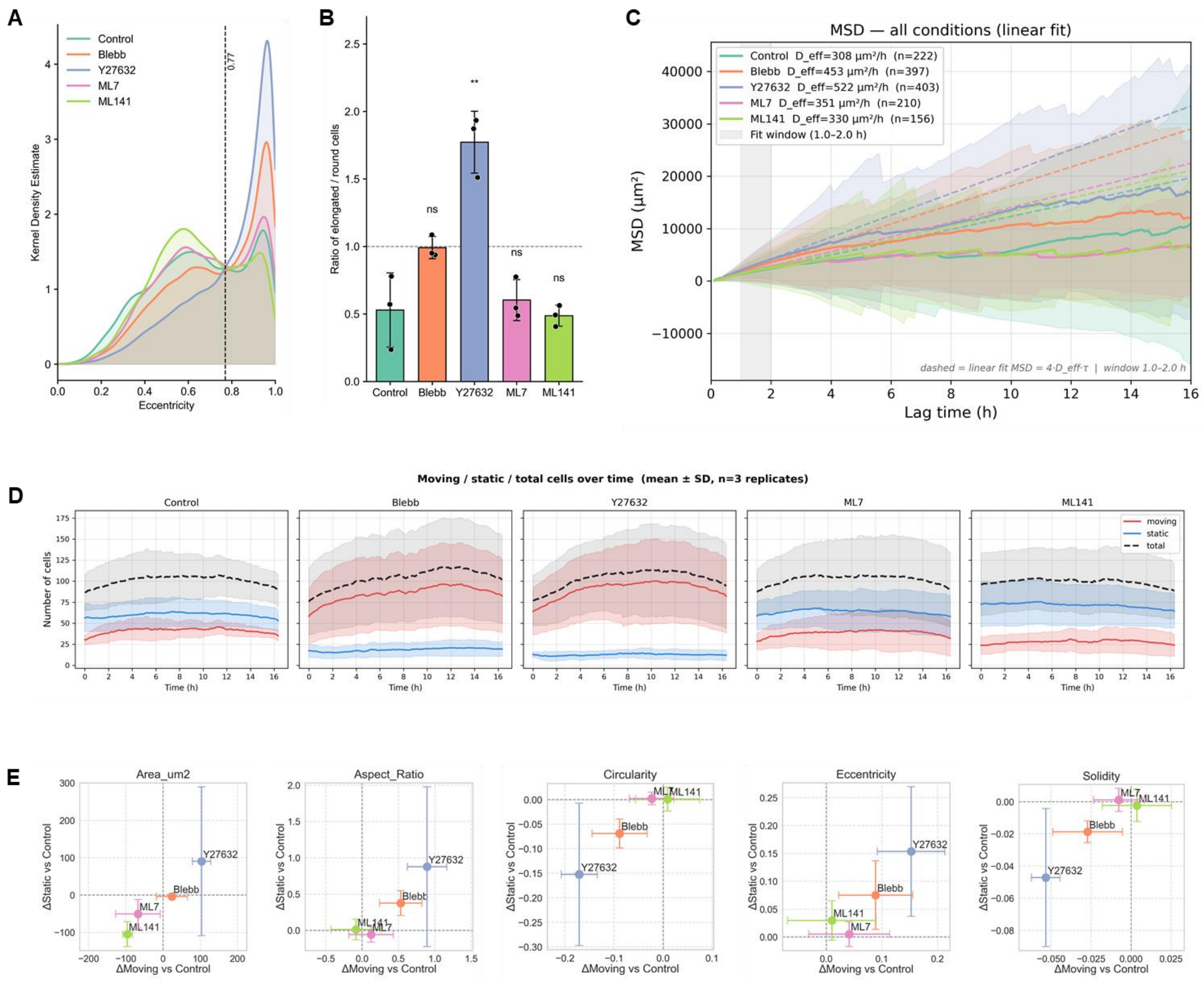


**Supplementary Figure S5 | Single-cell morphology and dynamics.** (**A**) Kernel density estimates (KDEs) of cell eccentricity distributions for migrating cells under different treatment conditions. Curves were generated from pooled measurements obtained from three independent biological replicates. The dashed vertical line indicates the eccentricity threshold (0.77) used to separate low- and high-eccentricity cell populations. (**B**) Ratio of high- to low-eccentricity cells, defined as the number of cells with eccentricity > 0.77 divided by the number of cells with eccentricity ≤ 0.77. Ratios were calculated independently for each biological replicate. Bars represent mean ± SD, and individual dots correspond to biological replicates (n = 3). Statistical significance was assessed using a t-test relative to the control condition (**P < 0.01). (**C**) Mean squared displacement (MSD) curves of migrating cells over the 16 h imaging period for all experimental conditions. Mean MSD values were calculated from all trajectories classified as moving. Dashed lines indicate linear fits performed over the 1–2 h lag-time interval, from which effective diffusion coefficients were derived. (**D**) Temporal evolution of the proportions of moving and static cells, together with the total number of tracked cells, over the 16 h acquisition period for each experimental condition. Cell trajectories were classified using custom movement criteria based on displacement, trajectory duration, and spatial spread. (**E**) Comparison of treatment-induced changes in cell morphology using the Wasserstein distance. For each morphological descriptor, Wasserstein distances were calculated relative to the control condition separately for moving and static cell populations. Distances for moving cells are plotted on the x-axis, whereas distances for static cells are plotted on the y-axis. Features located near the diagonal are affected similarly in both populations, whereas deviations from the diagonal indicate morphology changes preferentially associated with either moving or static cells.

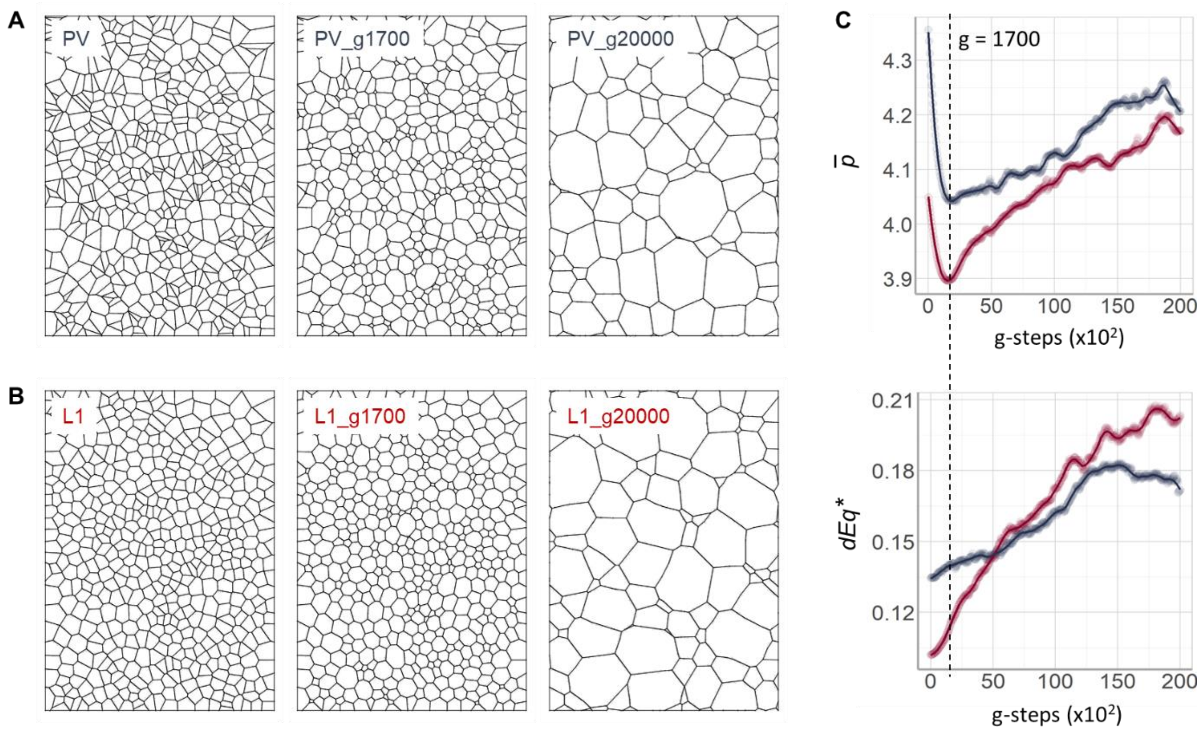


**Supplementary Figure S6 | Surface Evolver simulations using Voronoi tessellations.** (**A**) Evolution of a 500-generator Poisson–Voronoi (PV) tessellation after 0, 1700, and 20000 gradient-descent steps. (**B**) Evolution of a partially ordered Voronoi tessellation generated by one Lloyd iteration (L1) after 0, 1700, and 20000 gradient-descent steps. (**C**) Corresponding evolution of the mean mesh shape index, $\bar{p}$ (regularity), and normalized quantizer energy, $dEq^*$ (uniformity), for PV (blue) and L1 (purple) tessellations. Initially, network regularity increases, as indicated by a decrease in $\bar{p}$, whereas uniformity continuously decreases, reflected by an increase in $dEq^*$. Maximum regularity is reached at approximately $g \approx 1700$ gradient-descent steps, beyond which continued coarsening outweighs local edge-length homogenization, resulting in increasing mesh-size polydispersity and reduced regularity.

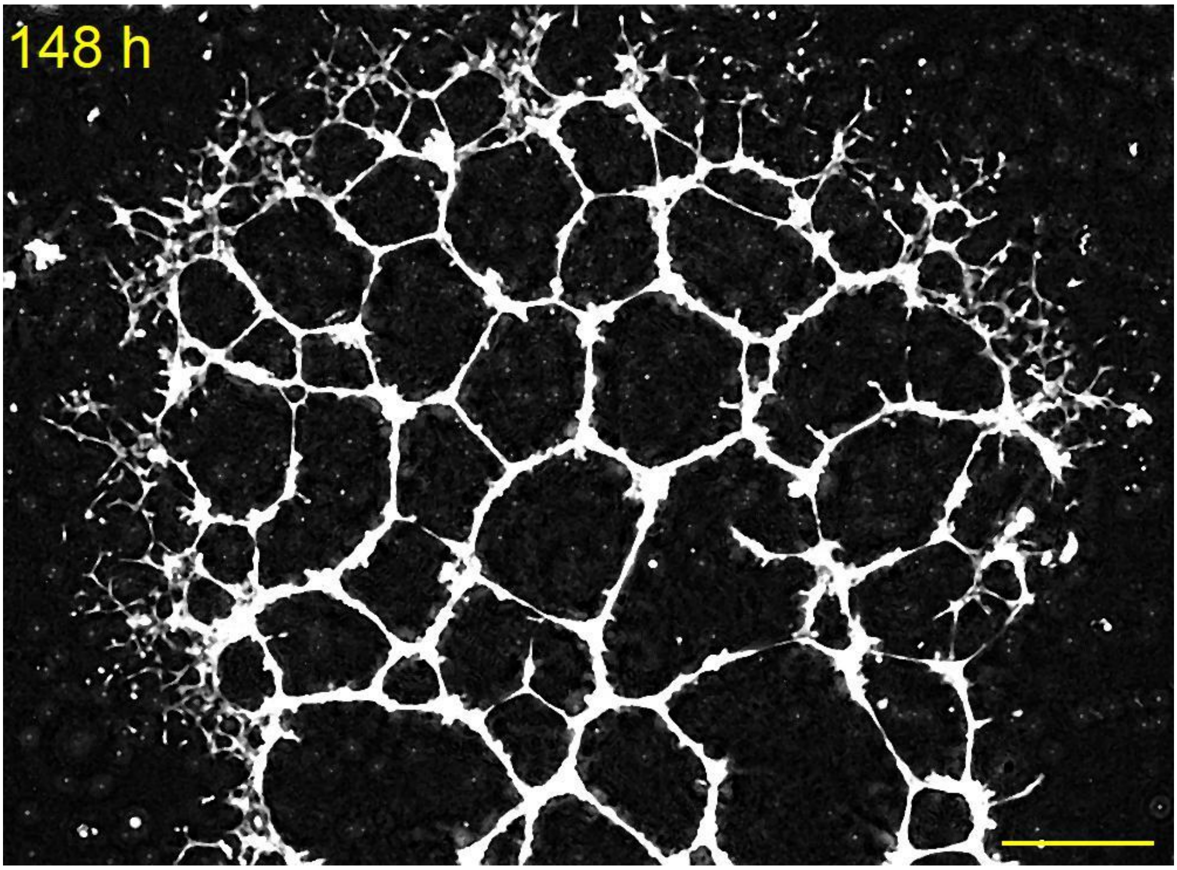


**Supplementary Video S1 | Formation of a quasi-2D epithelial network.** Cells seeded at low, spatially non-uniform density self-organize into hierarchically structured networks characterized by thick, fused branches in the central region and progressively finer branches toward the periphery, forming a self-similar architecture across scales. Network expansion occurs through a coordinated, wave-like front that invades initially unoccupied regions of the substrate. Scale bar: 0.5 mm.

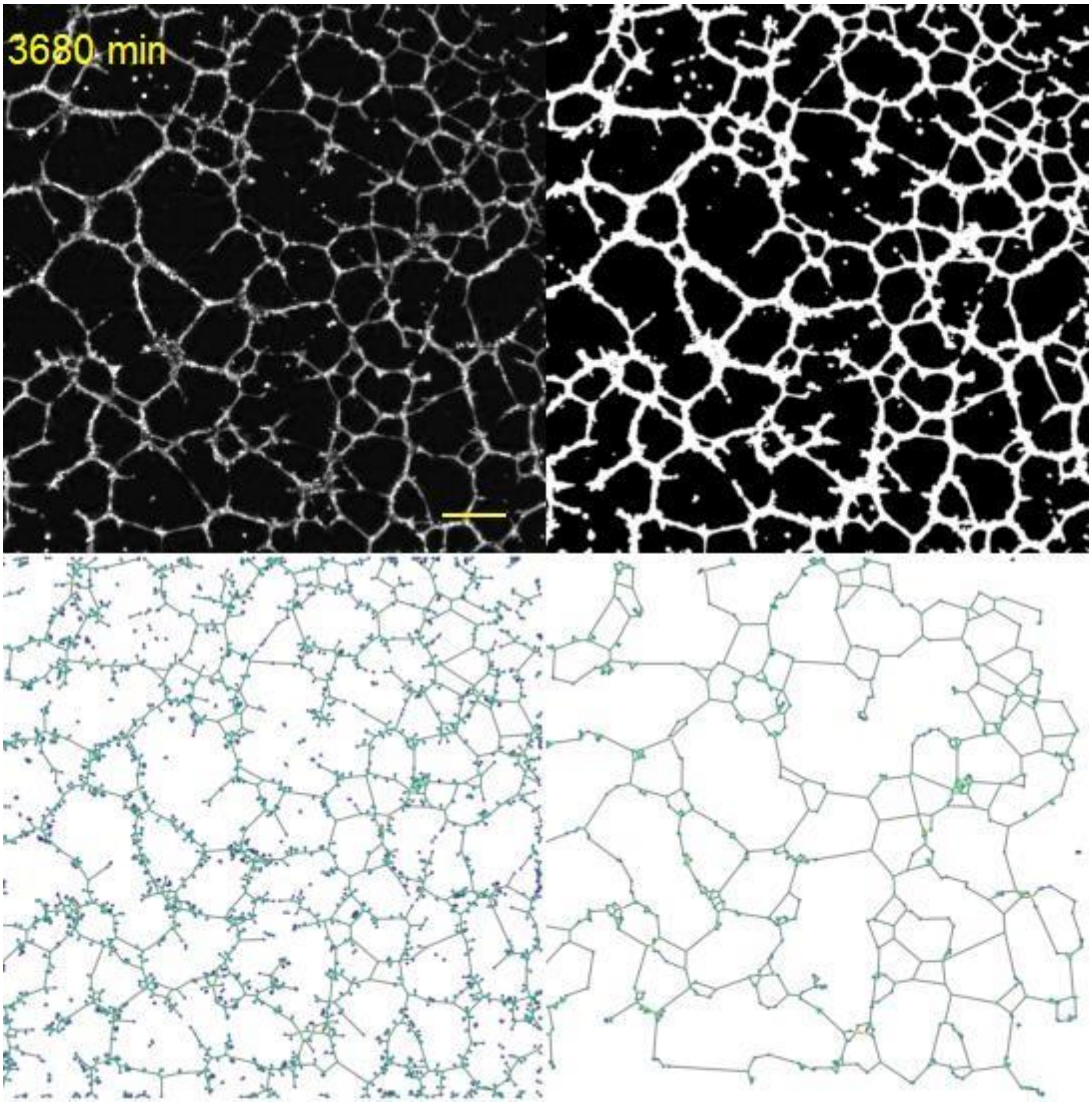


**Supplementary Video S2 | EpiNet analytical framework for epithelial network quantification.** Four-panel video illustrating the successive stages of the EpiNet workflow for quantitative spatiotemporal analysis of epithelial networks, from phase-reconstructed holographic images to CNN-based segmentation and binarized network representations, followed by extraction of planar graphs and their angle-filtered simplified counterparts (see Materials and Methods). Graph nodes are color-coded according to their degree: isolated nodes (violet), degree-1 nodes (dark blue), degree-2 nodes (blue), and degree-3 nodes (cyan). Scale bar: 0.5 mm.

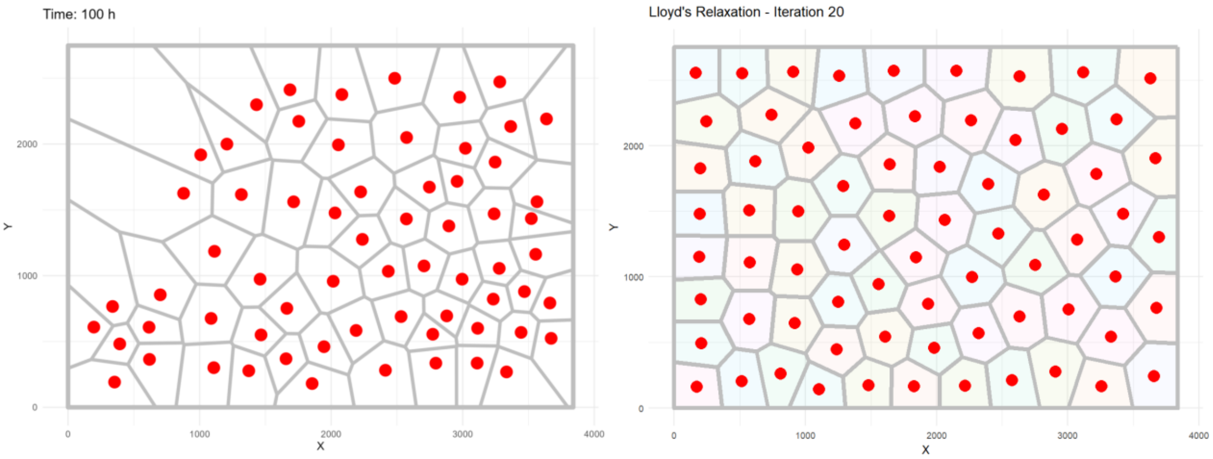


**Supplementary Video S3 | Comparison of EN relaxation dynamics with Lloyd's algorithm.** The right panel shows the evolution of experimental EN mesh centroids (red dots) during the network relaxation phase (T = 55–100 h), together with the corresponding centroidal Voronoi tessellations (CVTs). The left panel shows the evolution of a Voronoi tessellation generated from initially random Poisson-distributed generator points (red dots) undergoing 20 iterations of Lloyd's relaxation algorithm. The dimensions of the simulation domain (x, y) match the field of view of the lensless microscopy system and are expressed in image pixels.

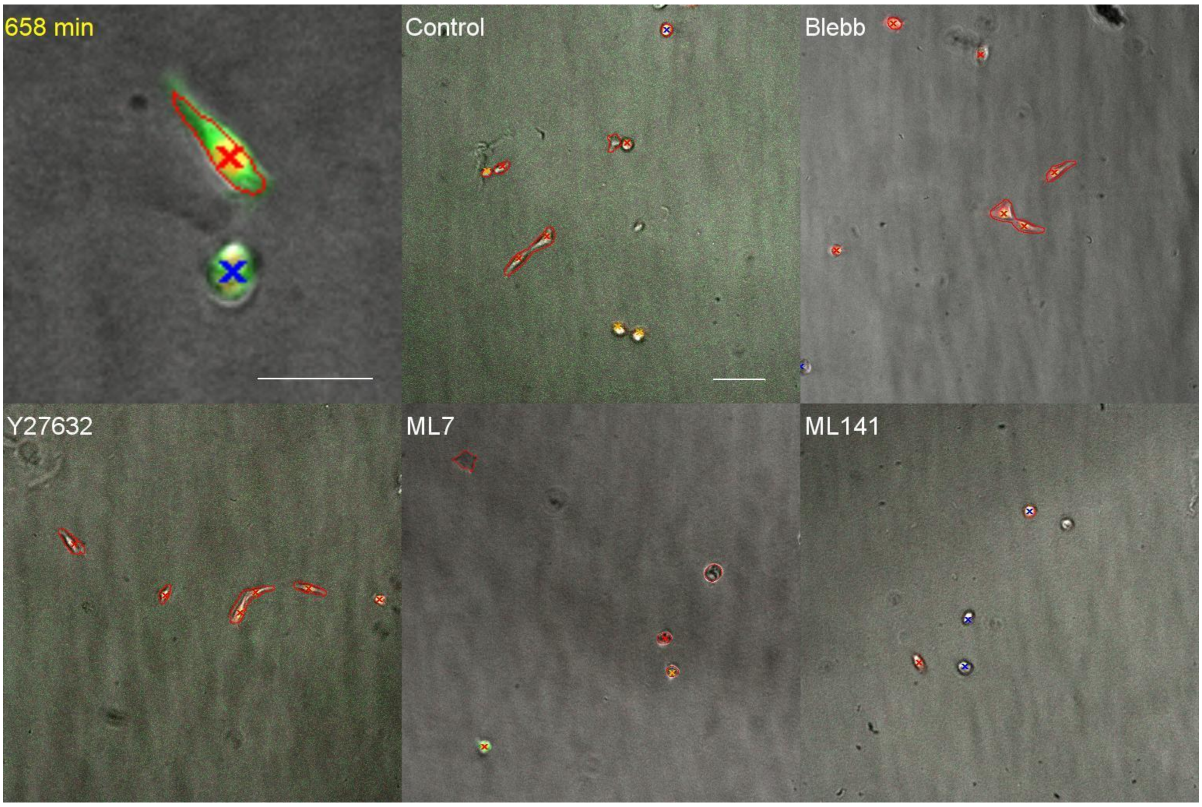


**Supplementary Video S4 | Single-cell morphology and dynamics.** Analysis of single-cell morphology and migration dynamics in reporter HPDE cells expressing membrane-localized Myr-mGL (green) and nuclear mCherry-H2B (red) fluorescent markers. Cell tracking was performed on mCherry-H2B fluorescence images using the Fiji TrackMate plugin for automated nuclear tracking. Extracted trajectories were subsequently processed using a custom Python-based classification pipeline to distinguish motile (red crosses) from static (blue crosses) cells and to remove trajectories arising from noise or tracking artefacts. Cell morphology was quantified from Cellpose-based segmentation masks (red contours) generated using a pre-trained model. Representative analyses of cells exposed to the indicated pharmacological treatments are shown. Scale bar: 50 µm.

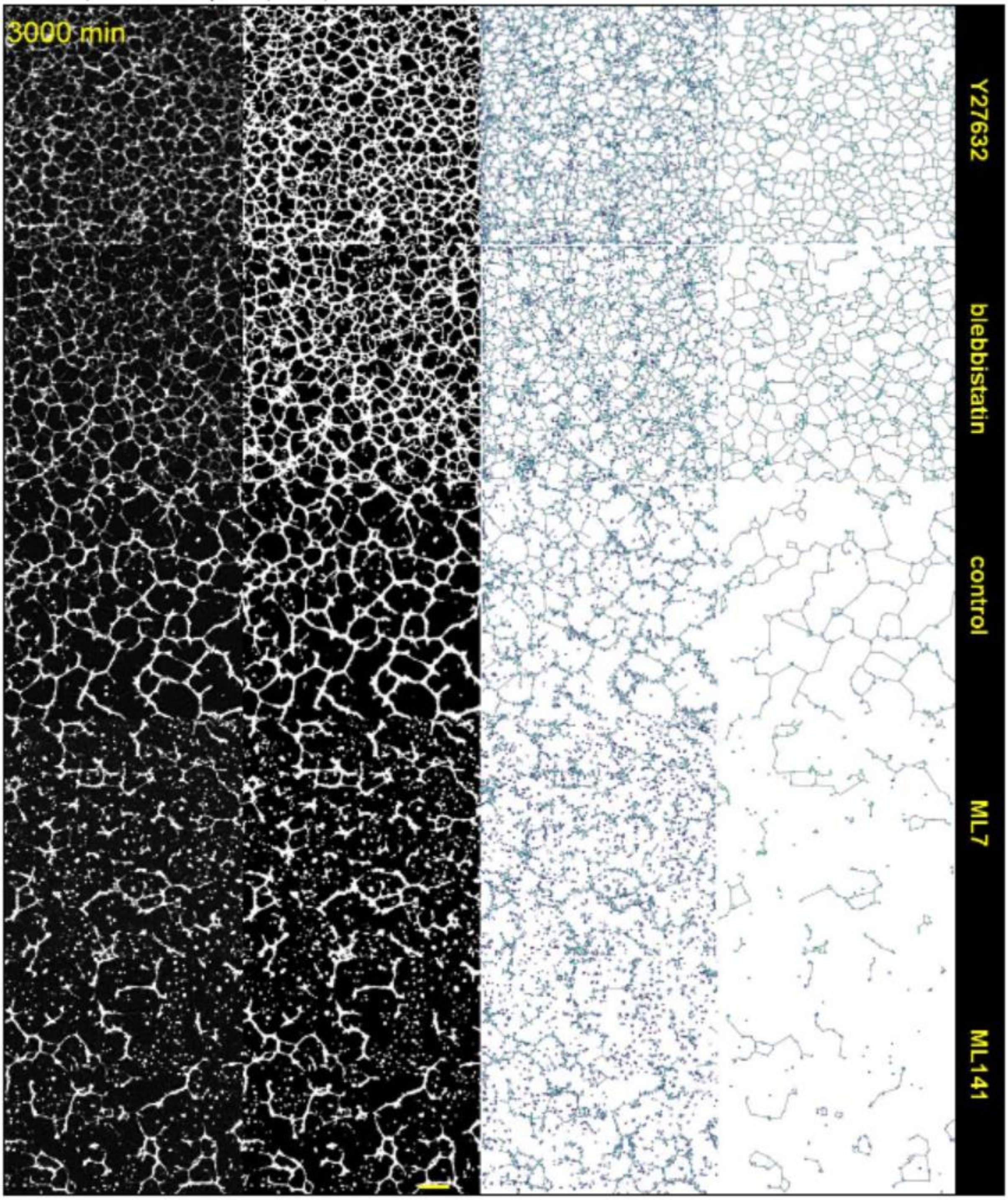


**Supplementary Video S5 | Effect of actomyosin modulation on epithelial network self-assembly.** For each pharmacological treatment condition (indicated on the right), the four-step EpiNet workflow is shown, enabling quantitative spatiotemporal characterization of epithelial network self-assembly. The workflow progresses from phase-reconstructed holographic images to CNN-based segmentation and binarized network representations, followed by extraction of planar graphs and their angle-filtered simplified counterparts (see Materials and Methods). Graph nodes are color-coded according to their degree: isolated nodes (violet), degree-1 nodes (dark blue), degree-2 nodes (blue), and degree-3 nodes (cyan). Scale bar: 0.5 mm.

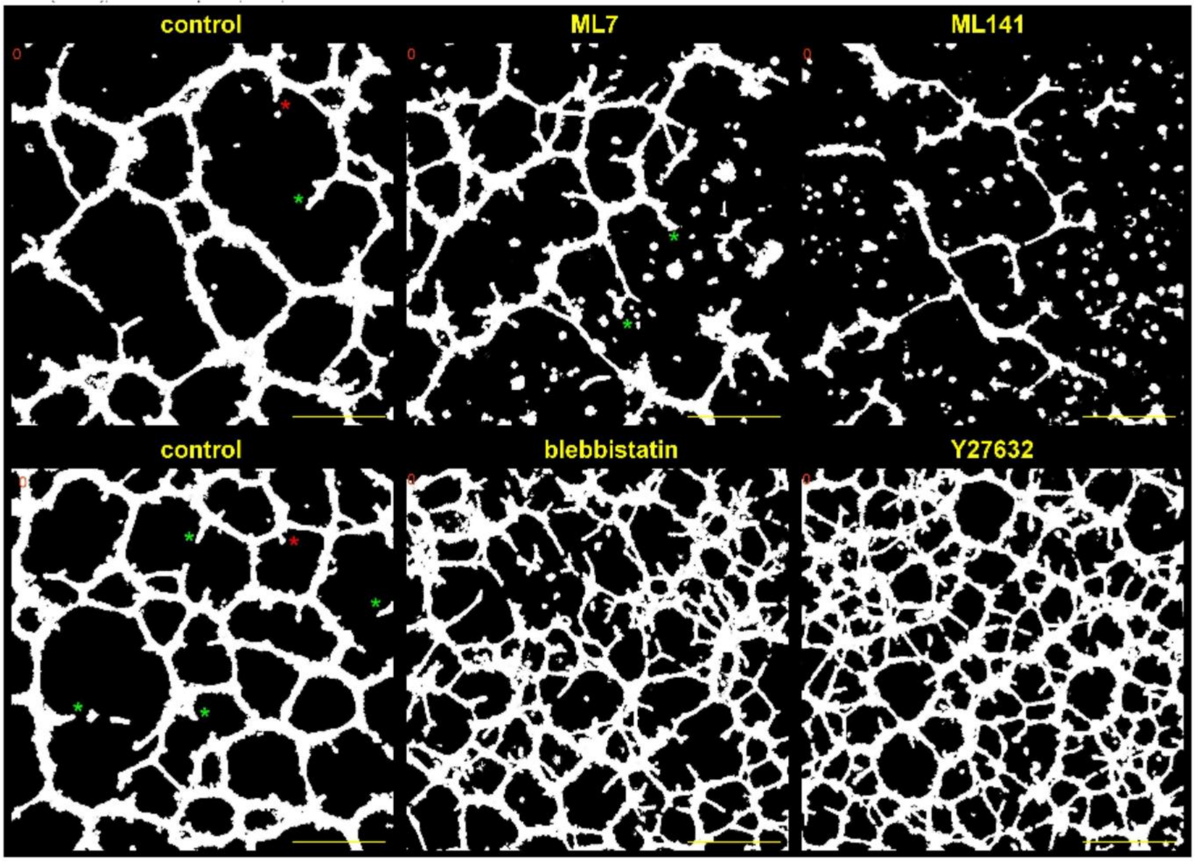


**Supplementary Video S6 | Topological rearrangements in epithelial networks.** Representative dynamics of epithelial networks showing frequent T0 topological transitions associated with mesh collapse (highlighted in yellow prior to disappearance), as well as occasional T1 topological rearrangements (magenta). Additional dynamic processes include vertex-angle equilibration, edge-length fluctuations, and pruning of unstable branches, which in non-spanning networks can lead to tension-driven collapse of the entire tree (e.g., ML141 condition). Frequent edge nucleation events emerging from pre-existing edges and forming transient T-junctions are indicated by asterisks; these events either regress (red asterisks) or stabilize to partition existing domains (green asterisks). In highly dynamic, dense networks induced by blebbistatin and Y27632, only T0 transitions are displayed for clarity (yellow). Time is indicated in minutes in the upper-left corner of each panel. Time windows are not synchronized across conditions. Scale bar: 0.5 mm.

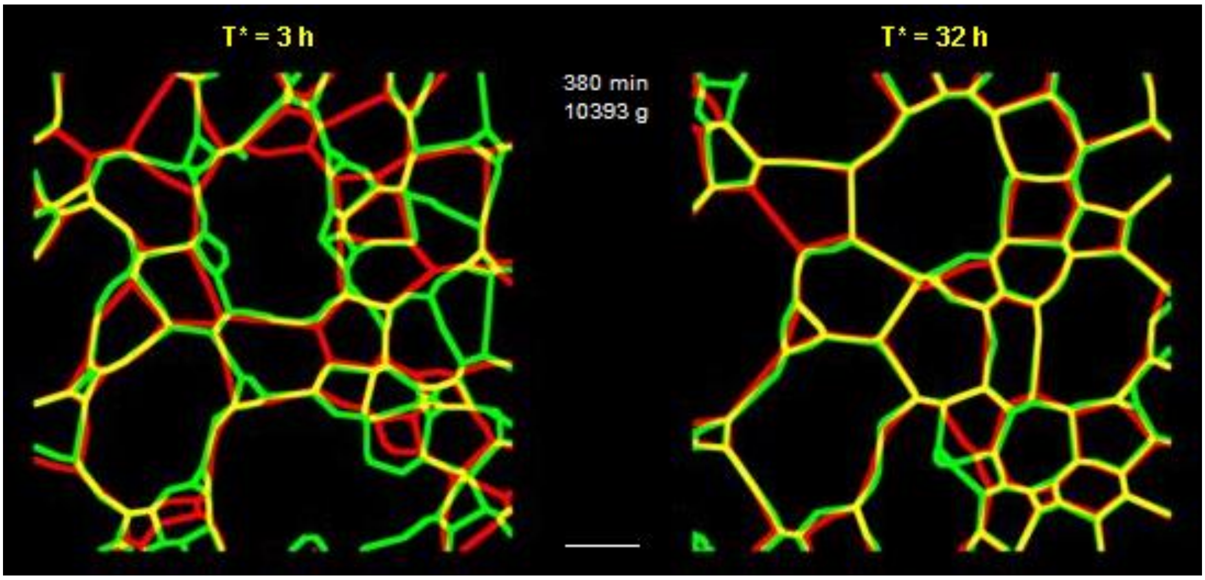


**Supplementary Video S7 | Prediction accuracy of the Surface Evolver STRING model.** Comparison of experimental (green) and simulated (red) network evolution, visualized through network overlap after rescaling gradient-descent steps to physical time. Progressive divergence between simulated and experimental geometries is indicated by the emergence of spatially segregated red and green regions. Prediction accuracy is lower at the onset of the relaxation stage (T* = 3 h, left panel) and increases substantially during later relaxation (T* = 32 h, right panel), suggesting that network dynamics become increasingly dominated by tension equilibration over time. Scale bar: 0.5 mm.

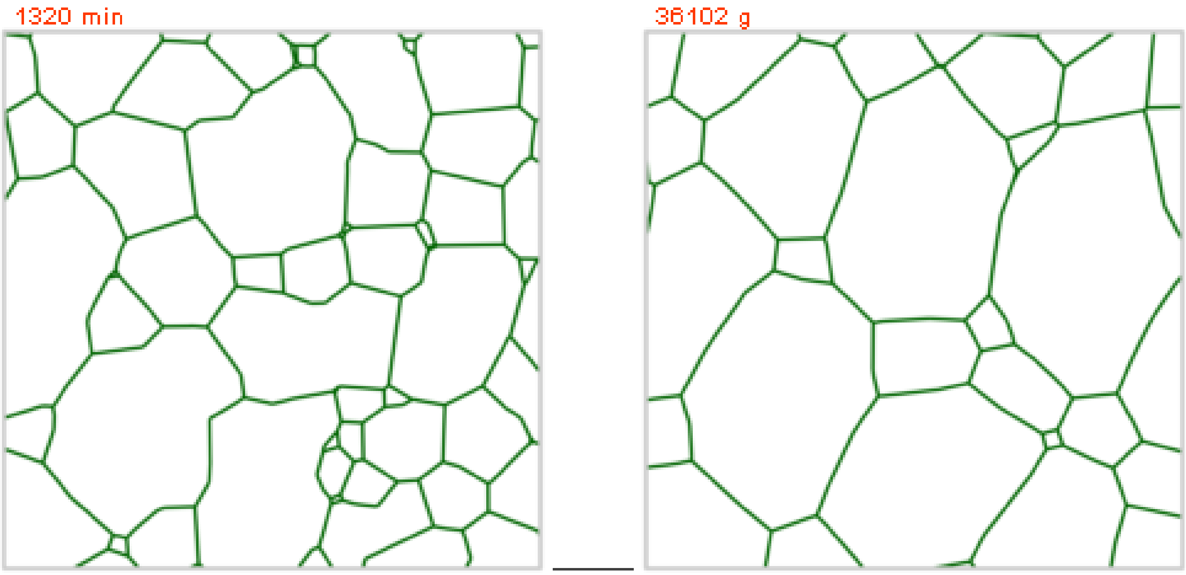


**Supplementary Video S8 | Limitations of the Surface Evolver STRING model**. Experimental networks (left panel) maintain an approximately constant mesh density while progressively increasing partition uniformity. This results from the interplay between tension-driven relaxation, which promotes force equilibration and geometric ordering, and domain-splitting events, which counteract coarsening and stabilize network scale. Together, these processes homogenize space partitioning while preserving a reticulate architecture. In contrast, without domain-splitting events, pure interfacial-energy minimization in the STRING model (right panel) drives network coarsening, characterized by the growth of larger domains at the expense of smaller ones, increased domain-size heterogeneity, and a reduction in mesh number with a corresponding increase in mean mesh area. Scale bar: 0.5 mm.
